# Type I TARPs regulate Kv7.2 potassium channels and susceptibility to seizures

**DOI:** 10.1101/2024.08.09.607194

**Authors:** Marina V. Rodrigues, Ângela S. Inácio, Maria Virginia Soldovieri, Sara Ribau, Telmo M. Leal, Theo Baurberg, Ilaria Mosca, Paolo Ambrosino, Luísa Amado, Nuno Beltrão, Gladys L. Caldeira, Luís F. Ribeiro, Ka Wan Li, Laurent Groc, Joana S. Ferreira, Maurizio Taglialatela, Ana Luisa Carvalho

**Author notes:** These authors contributed equally to this work (Sara Ribau, Telmo M. Leal).

## Abstract

The M-current is a low-threshold potassium current that modulates neuronal excitability and suppresses repetitive firing. However, the mechanisms regulating M-channel function remain unclear. We identified type I Transmembrane AMPA receptor Regulatory Proteins (TARPs) as M-channel Kv7.2 subunit interactors in cortical neurons, with their interaction increasing upon neuronal depolarization. Co-expression of TARPs with Kv7.2 increased channel surface expression and Kv7.2-mediated currents, while disrupting TARP-γ2 expression in neurons perturbed dendritic Kv7.2 nano-clusters and decreased M-currents. Knock-in mice with an intellectual disability-associated TARP-γ2 variant showed reduced hippocampal M-currents and increased seizure susceptibility, indicating that disrupting TARP-γ2 regulation of Kv7.2-M-channels is epileptogenic. These findings show that TARP-γ2, a synaptic protein crucial for excitatory transmission, also controls intrinsic excitability via M-channels. This discovery provides a link between synaptic transmission and neuronal excitability, with implications for disease, as the interplay between synaptic and intrinsic plasticity is pivotal to how the brain adapts to varying input signals.

**Highlights:**

- Type I TARPs bind to Kv7.2-M-channels and enhance Kv7.2-mediated currents.
- TARP-γ2 governs the neuronal nano-organization and function of Kv7.2 channels.
- Intellectual disability-associated TARP-γ2 variant impairs M-currents and facilitates seizures.
- Type I TARPs can serve as molecular integrators of synaptic and intrinsic excitability.

## Introduction

The integration of small stimuli simultaneously arriving at a neuron can generate a cumulative input signal that may trigger a fast depolarization event. Neuronal excitability is shaped by the duration and nature of these inputs. Intrinsic properties, such as resting membrane potential, the threshold for action potential generation, and input resistance, significantly contribute to defining neuronal excitability. Neurons use both synaptic and intrinsic plasticity to regulate network activity and maintain dependable computational capability, ensuring circuit stability while allowing for flexibility^1,2^. This prevents neuronal circuits from becoming excessively silent or overly excitable. Even though neuronal circuit function depends on the computational capacity of neurons and imbalances in excitability are linked to neurodevelopmental and psychiatric disorders^3–8^, the mechanisms governing neuronal excitability and responsiveness to synaptic inputs remain largely unknown.

The α-amino-3-hydroxy-5-methyl-4-isoxazolepropionic acid receptors (AMPAR) mediate most of the fast excitatory synaptic transmission in the brain. The dynamic nature of these receptors enables them to change in number, distribution, and functional properties in response to synaptic activity, which is crucial for the ability of the brain to adapt and learn^9^. One of the main protein families responsible for regulating these mechanisms is the Transmembrane AMPA Receptor Regulatory Proteins (TARPs) family. This family of proteins is subdivided into type I (TARP-γ2, -γ3, -γ4, and -γ8) and type II (TARP-γ5 and -γ7) TARPs based on whether they possess a typical or atypical PDZ-binding motif. The first protein identified in the TARP family was TARP-γ2, also known as stargazin. It was identified following the characterization of the stargazer mouse, which lacks TARP-γ2 expression and exhibits cerebellar ataxia, abnormal head tossing, absence seizures, and cognitive impairment^10^. TARP-γ2 regulates AMPAR cellular traffic and gating and alters receptor pharmacological properties^11^. Other members of the TARP family have been identified and studied in the context of their role in AMPAR regulation^11–13^, although their specific functions remain relatively unexplored. Despite showing decreased AMPAR-mediated excitatory transmission, the stargazer mouse presents unexplained cortical hyperexcitability, suggesting that other molecular players apart from AMPAR may play a role in this phenotype.

We have investigated the interactome of TARP-γ2 in cortical neurons by immunoprecipitating this protein and identifying its binding partners through mass spectrometry and identified the Kv7.2 M-channel subunit as a new putative interactor of TARP-γ2. M-channels are ubiquitously expressed in the brain and play a pivotal role in regulating neuronal excitability by delaying the onset of and/or hastening the termination of a burst of spikes, thereby causing spike frequency adaptation^14,15^. They are slow-activating voltage-gated potassium channels that do not inactivate, becoming active below the threshold for generating action potentials, a range where few other voltage-gated channels are operative^16^. Pathogenic loss-of-function variants of the *KCNQ2* gene, encoding the Kv7.2 subunit, have been repeatedly linked to a wide phenotypic spectrum of mostly early-onset epilepsies^17^ and developmental delay^18^. In rodents, *Kcnq2* deletion is perinatally lethal^19^, whereas *Kcnq3* (which encodes the Kv7.3 subunit) knock-out mice survive to adulthood^20,21^. Expression of dominant negative Kv7.2 subunit in mice decreases M-currents in CA1 pyramidal neurons by 75%, resulting in spontaneous epileptiform activity^22^. Interestingly, these animals present frequent head and neck extensions reminiscent of the stargazer behavior^22^. Additionally, conditional deletion of Kv7.2 from cortical pyramidal neurons results in abnormal electrocorticogram activity, increased excitability, and decreased medium afterhyperpolarization, whereas near-normal excitability was found in pyramidal neurons lacking Kv7.3 channels^23^. Moreover, a knock-in mouse model carrying one of the most recurrent variants in humans recapitulates at least some of the EEG and behavioral changes occurring in the most severe cases of KCNQ dysfunction^24^.

Despite their crucial role in regulating neuronal excitability, the mechanisms dictating the targeted localization and function of Kv7.2-M-channels are poorly characterized. Here, we report that in addition to regulating AMPARs and synaptic plasticity, type I TARPs also influence neuronal excitability by interacting with the Kv7.2 subunit of M-channels and enhancing Kv7.2-mediated currents. Genetic manipulation of TARP-γ2 expression in cortical neurons disturbed the nanoscale organization of Kv7.2 in dendrites and impaired neuronal M-currents. Moreover, an intellectual disability-associated variant of TARP-γ2 (V143L^25,26^) failed to modulate Kv7.2-mediated currents. Knock-in mice harboring this human mutation exhibited a decreased M-current density, and increased susceptibility to drug-induced seizures.

Altogether, our data provide seminal evidence for type I TARPs as regulators of Kv7.2 homomeric M-channels in the brain. Our findings support a dual role for TARP-γ2 in network activity control, serving as an auxiliary protein for both AMPARs and Kv7.2-M-channels. Thus, this discovery establishes a new connection between synaptic transmission and modulation of intrinsic neuronal excitability.

## Results

### Type I TARPs bind to the Kv7.2 subunit of M-channels

M-channels were first identified in the peripheral nervous system as heteromeric complexes composed of Kv7.2 and Kv7.3 subunits, primarily enriched in the axon initial segment (AIS) and Ranvier nodes^27–31^, which are crucial sites for regulating membrane potential and facilitating action potential generation and propagation. Recent evidence suggests that homomeric Kv7.2-M-channels play a more significant functional role in the somatodendritic compartment^20,32–34^. However, their physiological relevance remains largely unexplored and presents experimental challenges^15,35^. We have studied the interactome of TARP-γ2 in cortical neurons (Table S1) and identified the M-channel subunit Kv7.2 as a potential binding partner. To validate this interaction and investigate whether other type I TARPs can also interact with Kv7.2, we immunoprecipitated TARP-γ2, -γ3, and -γ8 from HEK293T cells co-expressing Kv7.2 and one of the TARPs. We observed that Kv7.2 co-immunoprecipitated with all type I TARPs tested (Figure 1A). To evaluate these interactions in neurons and determine their cellular localization, we performed an *in situ* proximity ligation assay (PLA) to visualize endogenous TARPs and Kv7.2 that co-localized within 40 nm^36^. We immunolabeled hippocampal cultured neurons for Kv7.2 and either TARP-γ2, -γ3, or -γ8 and detected a positive PLA fluorescent signal in the soma and dendrites (Figure 1B), suggesting an interaction of Kv7.2 with type I TARPs. Similar to hippocampal neurons, dendrites of cortical neurons displayed a positive PLA signal for Kv7.2 and TARP-γ2 (Figure S1A), indicating that these proteins interact in the somatodendritic compartment. A negligible PLA signal was detected when one of the primary antibodies was omitted from the reaction (Figure S1A, B), validating the specificity of the signal. In agreement with previous studies^37^, both Kv7.2 and TARP-γ2 were found to be endogenously expressed in all cellular compartments in cortical neurons, with Kv7.2 enriched at the AIS and TARP-γ2 at dendrites (Figure S1C). However, contrarily to dendrites, no PLA puncta were detected in the AIS or axons of cortical and hippocampal neurons at 15 days *in vitro* (DIV; Figure S1D, E). We then used dual-color direct stochastic optical reconstruction microscopy (dSTORM)^38^, a super-resolution imaging technique that allows for the simultaneous visualization of two different molecular species and enables the study of their co-localization at the nanoscale level. We expressed exogenous HA-tagged TARP-γ2 in cortical neurons and simultaneously labeled TARP-γ2 and Kv7.2. dSTORM imaging revealed dendritic nano-clusters of Kv7.2 that co-localized with TARP-γ2 (Figure 1C), with 35.6% of the Kv7.2 fluorescence signal overlapping the HA-TARP-γ2 signal (Figure 1D).

**Figure 1.**
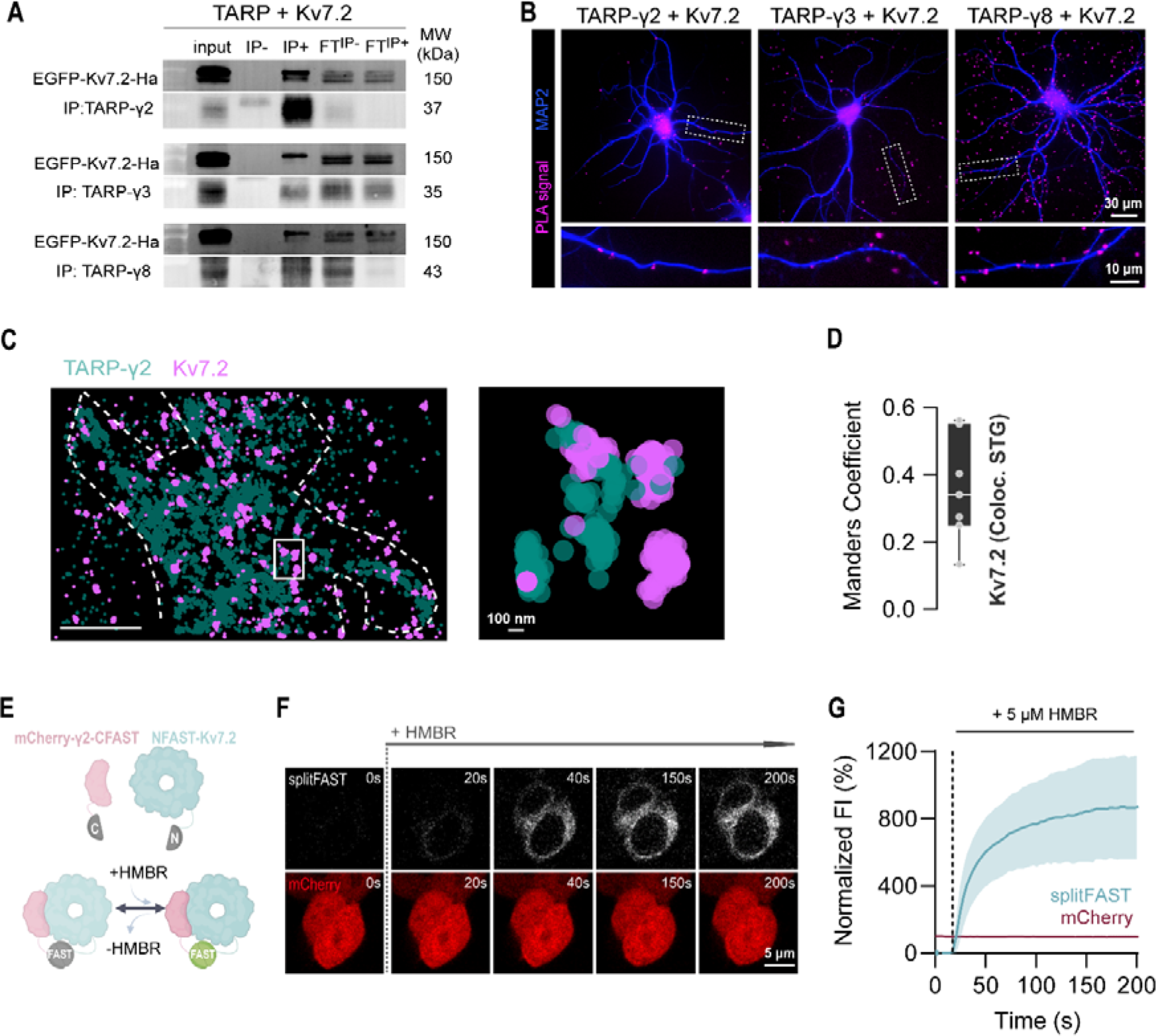
Type I TARPs interact with the Kv7.2 subunit of M-channels. A) Co-immunoprecipitation of Kv7.2 with TARP-γ2, -γ3 and -γ8 from co-transfected HEK293T cells. Immunoprecipitation (IP) assays were performed using either control non-immune rabbit IgGs (IP-) or specific rabbit antibodies against each TARP (IP+). Kv7.2 co-immunoprecipitated with all type I TARP tested (150 kDa band in the IP+ lane). The expression of each protein was confirmed in total lysates (input). FT^IP-^, control flow-through; FT^IP+^, IP flow-through. B) Representative images of proximity ligation assay (PLA) signal for type I TARPs and Kv7.2 in 15 DIV rat hippocampal neurons. PLA signal (pink) indicates spatial proximity between endogenously expressed TARP-γ2, -γ3, or -γ8 and Kv7.2. Neurons were also labelled for MAP2 (blue). C) Representative image of the dSTORM resolved nano-distribution of Kv7.2 and TARP-γ2 in 15 DIV cortical rat neurons, showing their co-localization in these cells. D) Manders’ co-localization coefficient between Kv7.2 and TARP-γ2 nano-clusters in cortical neurons overexpressing TARP-γ2. Data are presented as median ± interquartile range (IQR), with the box showing the 25^th^ and 75^th^ percentiles, whiskers ranging from the minimum to the maximum values, and the horizontal line indicating the median. N=4 independent experiments, n=7 neurons. E) Scheme depicting the experimental design of the splitFast assay to investigate the interaction between TARP-γ2 and Kv7.2 in live cells. CFast11 was fused to the C-terminal of TARP-γ2, while NFast11 was fused to the N-terminal of Kv7.2. The fluorescence signal requires the complementation of the Fast11 protein and depends on the binding of the HMBR fluorogen to the reconstructed Fast protein. F) Representative images showing the live interaction between TARP-γ2 and Kv7.2 in co-transfected HEK293T. The fluorescence signal (white) is only observed after adding HMBR, indicating that TARP-γ2 and Kv7.2 are physically interacting. The mCherry signal (red) was used to identify transfected cells. G) Temporal evolution of the normalized splitFAST and mCherry fluorescence intensities before and after adding HMBR. Data are shown as mean ± SD. N=2 independent experiments, n=55 cells. (See also Figure S1 and S2)

Next, we used splitFAST (split fluorescence-activating and absorption shifting tag) - a reversible fluorescence complementation system which specifically and reversibly binds fluorogenic hydroxybenzylidene rhodamine (HBR) analogues^39^ - to visualize the interaction between Kv7.2 and TARP-γ2 in live cells. We engineered mCherry-TARPγ2-CFAST-HA and NFAST-FLAG-Kv7.2 constructs (Figure S1F) that were co-expressed in HEK293T cells. Fluorescence was monitored before and after the addition of HMBR, which emits a green fluorescent signal when trapped inside the pocket of the reconstructed FAST (Figure 1E-G). The insertion of CFAST on the TARP-γ2 C-tail and of NFAST on the Kv7.2 N-tail did not affect the expression or trafficking of the proteins (Figure S1G). Expression of either portion of the FAST protein alone did not result in fluorescence emission (Figure S1H). However, the interaction between TARP-γ2 and Kv7.2 allowed FAST assembly and fluorescence emission upon HMBR binding (Figure 1G), further corroborating the co-immunoprecipitation data, and supporting a direct interaction between the two proteins.

Considering the function of M-currents in controlling hyperexcitability, we investigated whether neuronal depolarization could impact the endogenous interaction between TARP-γ2 and Kv7.2 in cultured cortical neurons. After 24 hours of treatment with 15 mM KCl, we observed a significant increase in the PLA signal for TARP-γ2 and Kv7.2 (Figure S2A, B). Additionally, we detected a significant distal shift of the AIS induced by chronic depolarization (Figure S2C, D), as previously reported^40^. These findings were independent of changes in AMPAR synaptic content or cell viability (Figure S2E-I). These observations imply that the interaction of TARP-γ2 with Kv7.2 increases in conditions of prolonged neuronal depolarization.

Taken together, these results indicate that Kv7.2 interacts with type I TARP family members. In neurons, Kv7.2 co-localizes with TARPs in the soma and dendrites. Moreover, the interaction between TARP-γ2 and Kv7.2 is strengthened during neuronal depolarization, indicating a dynamic interaction that may play a role in regulating the function or localization of Kv7.2 channels in response to changes in neuronal activity.

### Type I TARPs regulate Kv7.2-mediated currents and channel surface trafficking

To investigate the functional consequences of Kv7.2 interaction with TARPs, we performed whole-cell patch-clamp experiments to measure Kv7.2-mediated currents in transfected CHO cells, both in the absence and presence of each TARP. All type I TARPs (γ2, γ3, and γ4), except γ8, significantly increased the density of Kv7.2-mediated currents (Figure 2A, B). Co-expression of all TARPs resulted in Kv7.2 channel activation at less depolarized potentials (Figure 2C). The half activation potential (V_50_) for Kv7.2 was −22.72 ± −6.79 mV when expressed alone, shifting to values ranging from −26.10 to −32.31 mV in the presence of TARPs (Figure 2D). *KCNQ2* expression precedes that of *KCNQ3* during development, and most neonatal M-current is likely formed by Kv7.2 homomers^8,41^, but Kv7.2/7.3 heteromers are thought to comprise the majority of M-channels in the adult nervous system^27–31^. Hence, we investigated whether the impact of TARP-γ2 on M-current extended to heteromeric Kv7.2/7.3 heteromeric channels. Surprisingly, TARP-γ2 failed to modulate Kv7.2/Kv7.3-mediated currents, presenting itself as a specific regulator of Kv7.2 homomeric channels (Figure S3A-D).

**Figure 2.**
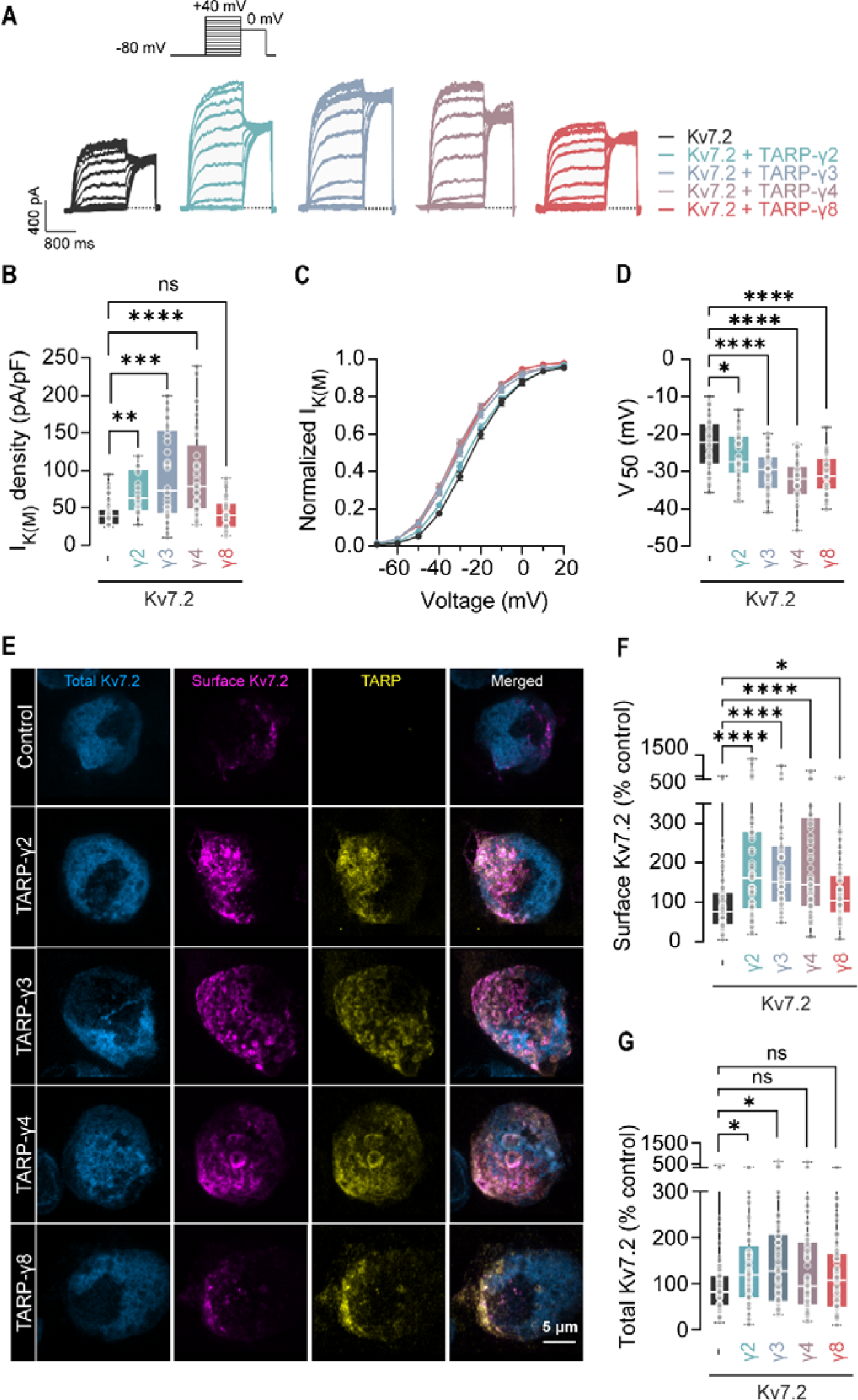
Type I TARPs differentially regulate Kv7.2 function and surface trafficking. A) Representative traces of Kv7.2-M–currents measured in CHO cells transfected with Kv7.2 alone or co-transfected with Kv7.2 and either TARP-γ2, -γ3, -γ4 or -γ8. To generate conductance–voltage curves, cells were held at −80 mV and depolarized for 1.5 seconds from −80 to +40 mV in 20 mV increments, followed by an 800 ms isopotential pulse at 0 mV. B) TARP-γ2, -γ3, and -γ4 co-expression increased currents mediated by Kv7.2 homomeric channels. Results are presented in whisker plots as median ± IQR (boxes show the 25^th^ and 75^th^ percentiles, whiskers range from the minimum to the maximum values, and the horizontal lines represent the median). Kruskal-Wallis (*p* < 0.0001) followed by Dunn’s multiple comparison test, \*\**p=*0.0021, \*\*\**p=*0.0001; \*\*\*\**p* < 0.0001, ns *p* > 0.9999. C) Current/voltage curves showing the impact of type I TARPs on Kv7.2 voltage-gating properties. Data are presented as mean ± SEM. D) The half-activation potentials (V_50_) were determined by fitting the current/voltage curves to a Boltzmann distribution. All type I TARPs significantly alter the gating properties of homomeric Kv7.2-M-channels, shifting the V_50_ to more negative potentials. Data are presented in whisker plots as median ± IQR. One-way ANOVA (*p* < 0.0001) followed by Dunnett’s multiple comparison test, \**p=*0.0307, \*\*\*\**p <* 0.0001. B and D) N ≥ 3 independent experiments, n=49 cells for Kv7.2, n=32 cells for Kv7.2 + TARP-γ2, n=32 cells for Kv7.2 + TARP-γ3, n=33 cells for Kv7.2 + TARP-γ4, n=34 cells for Kv7.2 + TARP-γ8. E) Representative fluorescence images showing the effect of type I TARPs on Kv7.2 total (blue) and surface (pink) expression levels. HEK293T cells were transfected with Kv7.2 alone or co-transfected with Kv7.2 and either TARP-γ2, -γ3, -γ4 or - γ8. An EGFP-Kv7.2-HA chimeric construct was used, and the extracellular hemagglutinin (HA) tag was labelled to visualize and quantify surface Kv7.2. The EGFP signal was used to determine the total levels of Kv7.2. TARPs (yellow) were immunolabeled with an anti-FLAG-tag antibody. F) Cell surface expression of Kv7.2 was significantly increased by TARP co-expression. Surface Kv7.2 was measured by quantifying the HA fluorescence intensity sum per cell, and data were normalized to the average intensity sum of the respective control (Kv7.2 alone). Kruskal-Wallis (p < 0.0001) followed by Dunn’s multiple comparison test, *p=0.0457, ****p < 0.0001. G) Total Kv7.2 expression was evaluated by quantifying the EGFP intensity sum per cell, and data were normalized to the average intensity sum of the respective control. Kruskal-Wallis (p=0.0296) followed by Dunn’s multiple comparison test, \**p=*0.0457 for TARP-γ2, \**p=*0.0162 for TARP-γ3, ns *p=*0.4397 for TARP-γ4, ns *p* > 0.9999 for TARP-γ8. F and G) Results are shown in whisker plots as median ± IQR. N=4 independent experiments, n=63 for Kv7.2, n=61 cells for Kv7.2 + TARP-γ2, n=62 cells for Kv7.2 + TARP-γ3, n=57 cells for Kv7.2 + TARP-γ4, n=6 cells for Kv7.2 + TARP-γ8. (See also Figure S3)

Given the role of TARPs in the surface trafficking of AMPAR^42–44^ and the observed increase in Kv7.2-mediated current density when co-expressed with type I TARPs (Figure 2A, B), we investigated whether TARPs affect the cell surface trafficking of Kv7.2. Co-expression of type I TARPs with Kv7.2 resulted in increased cell surface levels of the channel (Figure 2E, F). TARP-γ2 and -γ3 also induced an increase in the total expression of Kv7.2 (Figure 2G). In cortical neurons, overexpression of TARP-γ2 similarly augmented Kv7.2 levels (Figure S2E-G). Collectively, these data demonstrate that type I TARPs regulate both the current density and voltage-gating properties of Kv7.2 homomeric channels, as well as their trafficking to the cell surface.

### TARP-γ2 regulates dendritic Kv7.2 nano-clustering

While heterologous systems are convenient for clarifying specific aspects of the TARP-γ2/Kv7.2 interaction and its function, it is crucial to assess the neuronal significance of this interplay. Therefore, we used dSTORM to characterize the nanoscale topographic organization of native neuronal Kv7.2-containing M-channels, coupled with genetic manipulation of TARP-γ2 expression. Quantitative dSTORM has been successfully used to characterize the nano-organization of neurotransmitter receptors and ion channels in neuronal preparations^45–47^. To map all Kv7.2-containing M-channels, we labeled cortical neurons at 15 DIV using an antibody directed against an intracellular epitope of Kv7.2. As observed previously (Figure S1C), conventional epifluorescence imaging showed Kv7.2 immunoreactivity distributed along dendrites and axons (Figure 3A). Super-resolution dSTORM imaging revealed that Kv7.2-M-channel clusters were composed of several adjacent domains (Figure 3B) distributed across both dendrites and the AIS. Quantitative dSTORM imaging further elucidated the nanoscopic distribution of Kv7.2 clusters, revealing different nano-structural organizations in distinct compartments of cortical neurons. Notably, the size distribution of endogenous Kv7.2 nano-clusters observed in neurons (Figure S4A) closely resembled those previously observed in a heterologous system^48^. Segmentation of the super-resolved reconstructed images showed a higher density of clusters in dendrites than in the AIS (Figure 3D, E). These clusters were also larger (Figure 3F) and had a higher content of particle detection – local density - (Figure 3G). To our knowledge, this is the first report on the nanoscale cluster organization of native Kv7.2-containing M-channels in different neuronal compartments. The differential nanoscale mapping of Kv7.2 suggests different molecular mechanisms may operate to control dendritic- and AIS-localized channels.

**Figure 3.**
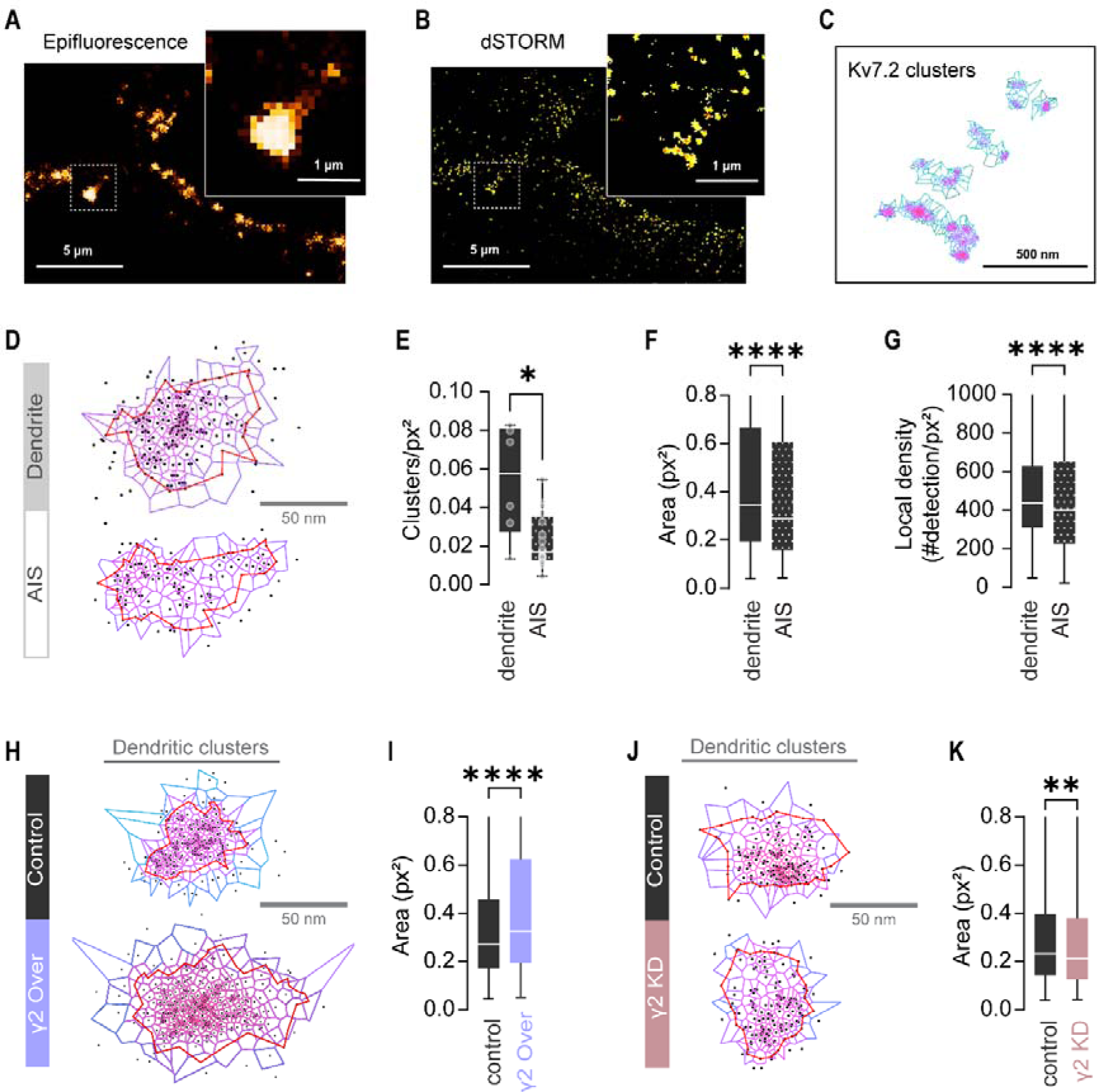
Differential neuronal compartmentalization of Kv7.2 at the nanoscale relies on balanced levels of TARP-γ2 expression. A) Representative image of 15 DIV rat cortical neurons immunolabeled for Kv7.2 and acquired using epifluorescence microscopy. B) The same neurons imaged with super-resolution dSTORM technique. C) Super-resolved Kv7.2 clusters from the zoomed image in B) segmented using SR-Tesseler analysis. D) Representative clusters of endogenously expressed Kv7.2 on dendrites and the axon initial segment (AIS) obtained with SR-Tesseler. MAP2 was used as a dendritic marker, while the AIS was identified as the soma-proximal region that was positive for EGFP but negative for MAP2 staining. Red lines delineate the cluster, with each dot representing an individual detection. Purple lines indicate the predicted polygons centered on each detection using the Voronoï diagram. E) Kv7.2 cluster density is higher in the dendritic compartment compared to the AIS. Data are shown in whisker plots as median ± IQR (boxes show the 25^th^ and 75^th^ percentiles, whiskers range from the minimum to the maximum values, and the horizontal lines represent the median). Unpaired t-test with Welch’s correction, p=0.0459, N ≥ 3 independent experiments, n=6 neurons for dendrites, n=17 neurons for the AIS. F) Dendritic Kv7.2 clusters have larger areas compared to Kv7.2 clusters in the AIS. Two-tailed Mann-Whitney test, p < 0.0001. G) Kv7.2 clusters in dendrites have a higher particle density than those in the AIS. Two-tailed Mann-Whitney test, *p* < 0.0001. F) and G) Data are shown as median ± IQR. N ≥ 3 independent experiments, n=1800 clusters randomly selected from all the acquisitions of the specific region. H) Representative clusters of endogenous Kv7.2 in the dendrites of cortical neurons following TARP-γ2 overexpression. Cells were transfected with either a control EGFP plasmid or a TARP-γ2-EGFP plasmid (γ2 Over), and transfected cells were identified by EGFP expression. I) Dendritic Kv7.2 cluster area in cortical neurons increases with TARP-γ2 overexpression. Two-tailed Mann-Whitney test, *p=*0.0001. Data are presented as median ± IQR. N=4 independent experiments, n=787 clusters randomly selected from all the acquisitions. J) Representative dendritic clusters of endogenous Kv7.2 following TARP-γ2 knockdown in cortical neurons. Cells were transfected with either a control EGFP plasmid or a TARP-γ2-specific EGFP-shRNA-encoding plasmid (γ2 KD). Transfected cells were identified by EGFP expression. K) TARP-γ2 knockdown decreases the area of Kv7.2 clusters in the dendrites of cortical neurons. Two-tailed Mann-Whitney test, *p=*0.0031. J) and K) Results are shown in whisker plots as median ± IQR. N=5 independent experiments, n=1050 clusters randomly selected from all the acquisitions. (See also Figure S4)

We then investigated whether the nanoscale organization of Kv7.2-containing M-channels in dendrites was governed by TARP-γ2 expression. The area of Kv7.2 dendritic clusters in cortical neurons overexpressing TARP-γ2 was significantly larger than that in clusters from control-transfected neurons (Figure 3H, I). Moreover, Kv7.2 clusters’ density – number of clusters per region of interest (ROI) - remained unchanged, while showing lower particle detection content (Figure S3B, C). In contrast, depletion of TARP-γ2 expression in neurons using an RNA interference strategy^26,49^ significantly diminished the area of endogenous Kv7.2 clusters (Figure 3J, K) despite no changes in cluster density or local particle detection (Figure S3D, E). All in all, our quantitative analysis of native Kv7.2 clusters using dSTORM imaging uncovered a role for TARP-γ2 in modulating the nanoscale organization of Kv7.2-M-channels within dendrites, particularly in influencing the size of Kv7.2 nano-clusters.

### Depleting TARP-γ2 in cortical neurons decreases M-currents

To investigate the functional significance of TARP-γ2 interaction with Kv7.2 in modulating neuronal M-currents, cortical neurons were genetically manipulated to suppress TARP-γ2 expression. Two different approaches were used to assess M-currents in whole-cell patch-clamp experiments (Figure 4). First, we perfused transfected neurons with retigabine, a specific activator of M-channels, to induce hyperpolarization of the resting membrane potential (Figure 4A), which was then measured in current-clamp recordings. Cortical neurons in which TARP-γ2 expression was silenced showed a significant decrease in the hyperpolarization induced by retigabine compared to control-transfected neurons (Figure 4B), supporting a role for endogenous TARP-γ2 in positively regulating M-currents. Similar to the findings described for the stargazer mouse^50^, there were no significant alterations detected in the resting membrane potential of TARP-γ2-depleted neurons compared to control neurons (Figure 4C).

**Figure 4.**
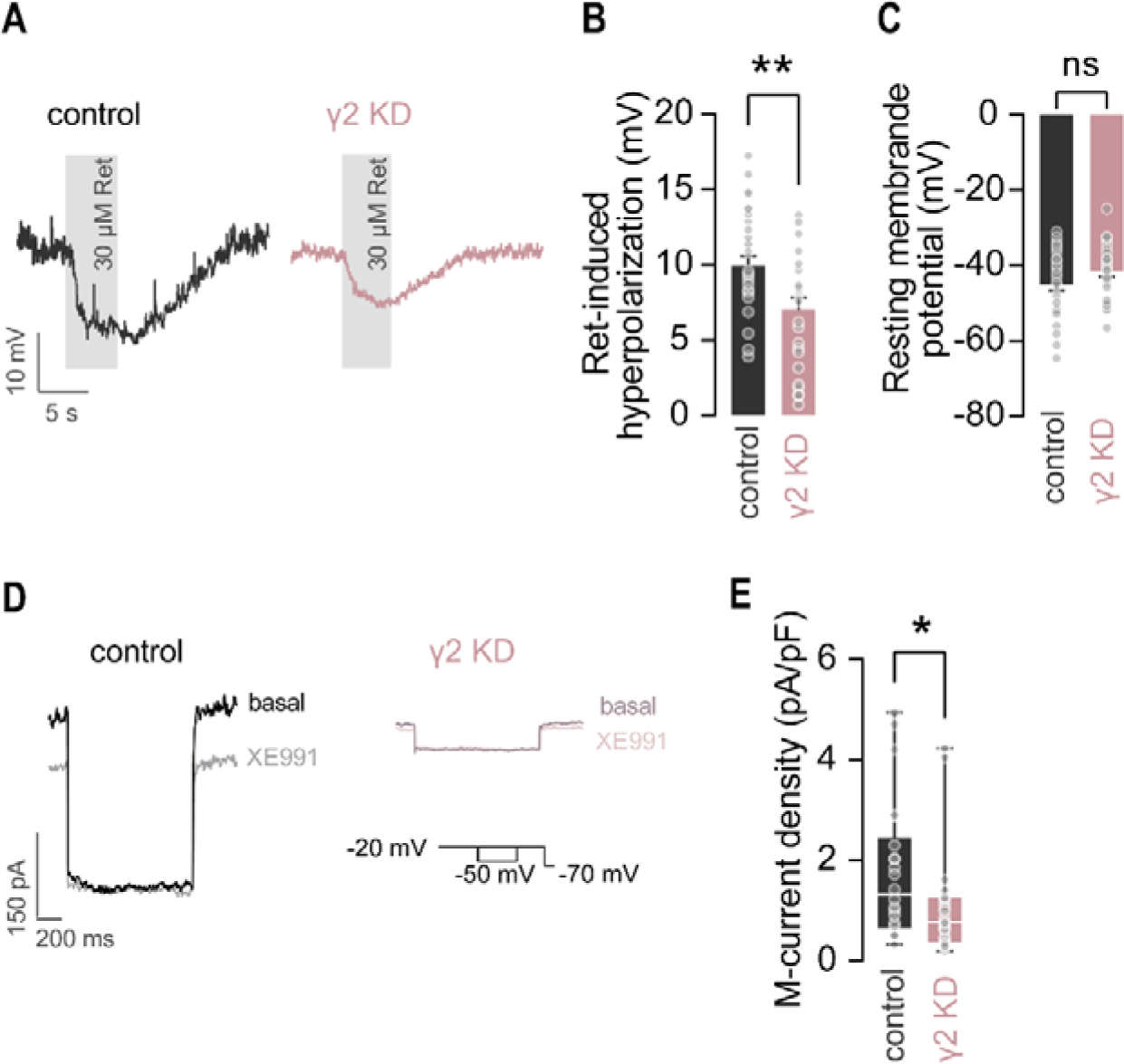
Silencing TARP-γ2 in cortical neurons decreases M-currents. A) Representative traces showing retigabine (Ret)-induced hyperpolarization of the resting membrane potential through activation of Kv7.2-containing M-channels in 15 DIV cortical neurons. Neurons transfected with a control plasmid, or a TARP-γ2-shRNA plasmid (γ2 KD) were held at 0 pA, and retigabine (30 µM) was perfused for 5 seconds, as indicated by the shaded grey area. B) Retigabine-induced hyperpolarization was significantly decreased in neurons following TARP-γ2 knockdown compared to control neurons. Two-tailed unpaired *t*-test, \*\**p=*0.0063. C) TARP-γ2 knockdown did not significantly alter the resting membrane potential in cortical neurons. Two-tailed unpaired *t*-test, ns *p=*0.0936. B and C) Data are presented as mean ± SEM. N=8 independent experiments, n=30 neurons for control, n=25 neurons for γ2 KD. D) Representative M-current traces from 15 DIV cortical neurons elicited by a 1-second repolarization step from −20 mV to −50 mV. M-currents in control neurons and neurons depleted of TARP-γ2 were measured as the XE991-sensitive component, calculated by subtracting the XE991-resistant component from the total current. E) Knockdown of TARP-γ2 resulted in a significant decrease of M-current density in cortical neurons. Two-tailed Mann-Whitney test, \**p=*0.0389, N=10 independent experiments, n=22 neurons for control, n=24 neurons for γ2 KD. Data are presented in whisker plots as median ± IQR (boxes show 25^th^ and 75^th^ percentiles, whiskers range from the minimum to the maximum values, and the horizontal lines represent the median).

Secondly, to directly measure M-currents, we analyzed the deactivation tail currents before and after perfusion with the M-channel blocker XE991^37,51,52^. The M-current, which corresponds to the XE991-sensitive component of the deactivation tail current, was significantly decreased in cortical neurons depleted of TARP-γ2 compared to control-transfected neurons (Figure. 4D, E). Altogether, our data clearly show that depleting the endogenous expression of TARP-γ2 in cortical neurons disrupts M-currents.

### Intellectual disability-associated TARP-γ2 variant decreases M-current function and facilitates seizures

The TARP-γ2 V143L variant, associated with intellectual disability^25^, has been shown to induce cognitive and social behavior alterations in mice, along with impairments in hippocampal synaptic transmission and plasticity^26^. Here, we investigated whether this mutation disrupts the functional regulation of Kv7.2 by TARP-γ2. Whole-cell patch clamp experiments were conducted to evaluate Kv7.2-mediated currents in CHO cells co-transfected with Kv7.2 and either wild-type TARP-γ2 or the intellectual disability-associated variant (Figure 5A-C). Unlike the wild-type protein, TARP-γ2 V143L did not induce potentiation of Kv7.2-mediated currents *in vitro* (Figure 5C).

**Figure 5.**
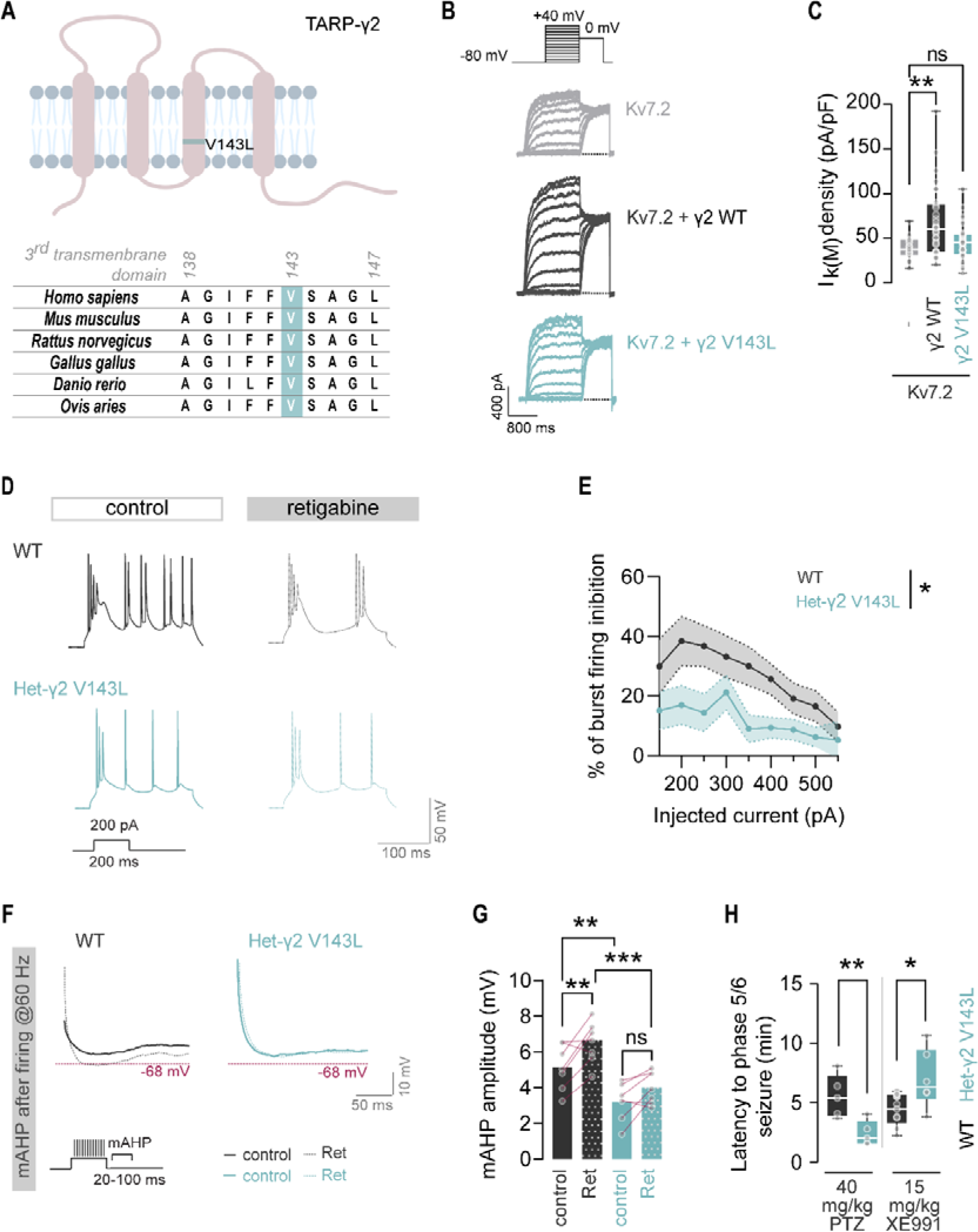
Intellectual disability-associated mutant of TARP-γ2 (V143L) impairs M-currents in subicular pyramidal neurons and increases seizure susceptibility. A) Schematic representation of TARP-γ2 highlighting an intellectual disability-linked missense mutation that leads to the replacement of valine 143 with leucine (V143L) in the third transmembrane domain^23^. This mutation is located at a highly conserved residue of the protein. B) Representative traces of Kv7.2 homomeric M-currents measured in CHO cells transfected with Kv7.2 alone or co-transfected with Kv7.2 and either wild-type TARP-γ2 (γ2 WT) or the intellectual disability-associated mutant form of TARP-γ2 (γ2 V143L). To generate conductance-voltage curves, cells were held at −80 mV and depolarized for 1.5 seconds from –80 to +40 mV in 20 mV increments, followed by an 800 ms isopotential pulse at 0 mV. C) Unlike wild-type TARP-γ2, the TARP-γ2 V143L variant did not potentiate Kv7.2-mediated currents. Data are presented in whisker plots as median ± IQR (boxes show 25^th^ and 75^th^ percentiles, whiskers range from the minimum to the maximum values, and the horizontal lines represent the median). Kruskal-Wallis (*p=*0.0060) followed by Dunn’s multiple comparison test, \*\**p=*0.0066; ns *p=*0,4811. N ≥ 3 independent experiments, n=18 cells for Kv7.2, n=41 cells for γ2 WT, n=22 cells for γ2 V143L. D) Representative traces of evoked action potentials in response to a 200 pA current step recorded from subicular burst-firing pyramidal neurons of wild-type (WT) and heterozygous TARP-γ2 V143L (Het-γ2 V143L) 8-week-old mice, both before and during perfusion with retigabine (30 µM). E) Subicular neurons from heterozygous TARP-γ2 V143L knock-in animals are less sensitive to retigabine compared to WT neurons. Data are presented as the mean ± SEM of the percentage of retigabine-induced firing inhibition. Two-way repeated measures ANOVA, *p =* 0.0387 (genotype effect). F) Representative traces of the medium after-burst hyperpolarization (mAHP) recorded from subicular burst-firing pyramidal neurons of WT and Het-γ2 V143L mice under control conditions and during perfusion with retigabine (Ret, 30 µM). The mAHP amplitude was calculated from action potential trains where the injected current induced firing at 60 Hz. G) Het-γ2 V143L mice showed a significant reduction in the amplitude of the mAHP in subicular pyramidal neurons compared to WT mice, and the mAHP amplitude in these animals did not change in response to retigabine. Data presented as mean ± SEM. Two-way repeated measures ANOVA (*p*=0.0011, genotype effect; *p=*0.0020, retigabine effect; p=0.3398, interaction effect) followed by Šídák’s multiple comparison test, \*\**p=*0.0090 control WT vs Ret WT, ns *p=*0.1156 control Het-γ2 V143L vs Ret Het-γ2 V143L, \*\**p=*0.0060 control WT vs control Het-γ2 V143L, \*\*\**p=*0.0006 Ret WT vs Ret Het-γ2 V143L. D-G) N ≥ 4 animals, n=7 neurons for WT, n=7 neurons for Het-γ2 V143L. H) 8-week-old Het-γ2 V143L mice presented increased susceptibility to seizures induced by a single dose of 40 mg/kg pentylenetetrazole (PTZ) and reduced susceptibility to seizures induced by a single dose of 15 mg/kg XE991, a blocker of M-channels. Latency to seizures was determined by the time required for each animal to reach phase 5/6 of adapted Racine’s scale of epileptogenesis after injection. Data presented as mean ± SEM. For the PTZ experiment N=5 animals for both genotypes. Two-tailed unpaired t-test, **p=0.0097. For the XE991 experiment N=9 animals for WT, N=6 animals for Het--γ2 V143L. Two-tailed unpaired t-test, *p=0.0166. (See also Figure S5)

To elucidate the significance of these findings, we assessed M-currents in the brains of TARP-γ2 V143L knock-in mice. Given the substantial co-enrichment of TARP-γ2 and Kv7.2 in the subiculum (Figure S5A), we characterized the intrinsic properties (Figure S5D-I) and measured M-currents in pyramidal neurons within this brain region. To investigate the impact of TARP-γ2 V143L on neuronal firing activity and the response to retigabine, cells were current-clamped at 0 pA for 120 ms, followed by depolarization for 200 ms with increasing steps of 50 pA, and then held at 0 pA for 4 seconds. This protocol was conducted under control conditions and repeated during retigabine perfusion, which activates M-channels and consequently decreases the firing rate. We observed that retigabine reduced burst activity in wild-type neurons (Figures 5D and S5J) but had minimal effect on neurons from heterozygous TARP-γ2 V143L mice (Figures 5D and S5K). The efficacy of retigabine, measured as the percentage of burst firing inhibition, was significantly reduced in TARP-γ2 V143L neurons compared to wild-type (Figures 5E and S5J, K). These findings suggest a reduced contribution of M-currents in attenuating burst activity in knock-in animals.

The M-current contributes to the medium and slow after-burst hyperpolarization (mAHP and sAHP)^20^. To further investigate the impact of the TARP-γ2 V143L variant on M-currents, we compared the mAHP in burst-firing pyramidal cells located in the subiculum of wild-type and heterozygous TARP-γ2 V143L mice (Figure 5F). The mAHP was defined as the amplitude of the afterhyperpolarization peak relative to the membrane resting potential, measured within the 20-100 ms interval following bursts fired at 50-60 Hz. Heterozygous TARP-γ2 V143L mice exhibited a significantly reduced basal mAHP compared to their wild-type littermates (Figure 5G). Furthermore, while wild-type subicular pyramidal neurons showed an increase in mAHP amplitude in response to retigabine, the mAHP in neurons from TARP-γ2 V143L heterozygous mice was not affected by retigabine perfusion (Figure 5G).

Our findings indicate a disturbance in the regulation of M-currents caused by the TARP-γ2 V143L variant. To examine the implications of this disruption in M-current function, we investigated the susceptibility of these mice to seizures. Notably, as per current knowledge, TARP-γ2 V143L knock-in mice do not have spontaneous seizures and even show adaptive changes in the intrinsic properties of subicular neurons (Figure S5D-I). To evaluate seizure susceptibility in these mice, we administered pentylenetetrazol (PTZ), a convulsive agent known to inhibit GABAergic transmission, to 8-week-old wild-type and TARP-γ2 V143L heterozygous mice. The latency to reach epileptogenic phase 5/6 was determined according to the PTZ-modified Racine scale^53^. Heterozygous TARP-γ2 V143L animals exposed to PTZ exhibited significantly shorter latencies to reach stage 5/6 seizures (Figure 5H), suggesting facilitated seizure susceptibility in these animals. We hypothesized that defective M-channel function in TARP-γ2 V143L knock-in mice might confer partial protection against seizure induction by an M-channel blocker. Indeed, the latency to seizures was prolonged in heterozygous TARP-γ2 V143L mice compared to wild-type littermates when they were injected with XE991, an M-channel blocker used to induce seizures (Figure 5H).

Multiple lines of evidence suggest dysfunctional M-channel activity in mice expressing the TARP-γ2 V143L variant. Compared to wild-type mice, heterozygous TARP-γ2 V143L animals exhibited: i) a reduced inhibitory effect of retigabine on burst firing in subicular pyramidal cells, ii) negligible potentiation of the mAHP by retigabine, and iii) lower sensitivity to XE991-induced seizures. Overall, our results support impaired control of burst activity due to compromised M-channel function in mice expressing the intellectual disability-associated V143L TARP-γ2 variant, correlating with increased susceptibility to PTZ-induced seizures.

## Discussion

Synaptic plasticity and intrinsic excitability are pivotal for shaping neural activity and information processing in the brain. However, how these mechanisms coordinate to regulate various aspects of network activity remains poorly understood. We have discovered an unexpected role for the type I TARP family of synaptic proteins, traditionally known as AMPAR auxiliary subunits, as regulatory proteins of Kv7.2-M-channels, playing crucial roles in regulating both their organization and function. Our data show that type I TARPs interact with Kv7.2, increase the channel surface expression, enhance Kv7.2-mediated-currents and modulate their voltage dependence. These findings support a dual role for TARPs in regulating network activity, not only through synaptic transmission and plasticity but also through intrinsic plasticity.

### Type I TARPs are regulatory subunits of Kv7.2-M-channels

In this work, we show that type I TARPs, which are well-described auxiliary subunits of AMPARs, interact with Kv7.2 and modulate its function. Biochemical and imaging assays identified TARP-γ2, -γ3, -γ4, and -γ8 as interaction partners of the Kv7.2 subunit of M-channels and revealed that they promote the surface trafficking of Kv7.2 homomeric channels. Furthermore, electrophysiology assays showed that type I TARPs enhance Kv7.2-mediated currents and modulate the gating properties of the channel. Importantly, endogenous neuronal TARP-γ2 regulates M-channels, as bi-directional manipulation of its expression in cortical neurons disrupted the nano-structure of Kv7.2 dendritic clusters, and TARP-γ2 depletion in these neurons decreased M-currents. Therefore, TARP-γ2 meets the criteria to be considered an auxiliary subunit of Kv7.2 channels^54^: i) it does not function as an active channel itself; ii) TARP-γ2 and Kv7.2 physically interact; iii) TARP-γ2 directly modulates the trafficking of Kv7.2; and iv) TARP-γ2 regulates Kv7.2 in native tissue. TARP-γ3 and -γ8 fulfill criteria i-iii and are strongly co-localized with Kv7.2 in the dendrites of hippocampal neurons. Although, due to technical limitations, we do not have data regarding criterion ii, TARP-γ4 also fulfills criteria i and iii. However, further studies are needed to clarify how TARP-γ3 and -γ8 regulate Kv7.2 in neurons. Given that TARP family members have distinct (albeit partially overlapping) expression patterns in the brain, different TARPs may be critical for Kv7.2 regulation across various brain regions, similar to how this family of proteins regulates AMPARs^55^.

Our results demonstrate that TARP-γ2 regulates ion channels from unrelated families, as we found that it cross-modulates Kv7.2, in addition to its well-known role as an auxiliary protein of AMPAR. Previous examples of cross-modulation by auxiliary proteins include the modulation of the biophysical properties of Kv1, Kv4.1, and Kv7 channels by Navβ1^56,57^, as well as the regulatory role of KCNE1, which modulates both cardiac KCNQ1 channels and TMEM16A chloride channels, belonging to two distinct ion channel superfamilies^58^. Our study adds to this body of evidence, supporting the concept of cross-modulation by auxiliary proteins.

### TARP-γ2 orchestrates Kv7.2 dendritic nano-clustering and regulates Kv7.2-M-currents

Our data support a role for type I TARPs in enhancing currents mediated by homomeric Kv7.2 channels but not by heteromeric Kv7.2/Kv7.3 channels. Heteromerization of M-channels is described as necessary for channel anchoring at the active sites of the AIS and axons^30,31,37,59–61^, suggesting that homomeric Kv7.2 channels are excluded from these compartments. Therefore, it is likely that dendritic Kv7.2 homomeric channels are preferentially, if not exclusively, regulated by type I TARPs. Indeed, the PLA signal for type I TARP-Kv7.2 interactions was found predominantly in the dendrites of hippocampal and cortical neurons. Furthermore, the dendritic compartment of cortical neurons exhibited a higher density of Kv7.2 clusters, which were larger and presented higher molecule content than those observed in the AIS, where heteromeric Kv7.2/Kv7.3 channels are more abundant. The nanoscale organization of dendritic Kv7.2 clusters is coordinated by TARP-γ2. Overexpression of TARP-γ2 led to larger Kv7.2 clusters and overall Kv7.2 increased expression, while the knockdown of TARP-γ2 decreased the area of Kv7.2 clusters and resulted in smaller M-currents in cortical neurons. These results raise the possibility that the decrease in cortical M-currents following TARP-γ2 knockdown may be a consequence of lower Kv7.2 homomeric currents.

### Disrupted M-currents and increased seizure susceptibility in mice with a point mutation in TARP-γ2 linked to intellectual disability

Seizures arise from disruptions in mechanisms that control neuronal excitability, and the M-current is one such mechanism^23,62,63^. Studies show a high comorbidity of epilepsy with most neuropsychiatric disorders, particularly intellectual disability^64–67^. Here, we show that a point mutation in TARP-γ2 linked to intellectual disability perturbs the functional regulation of Kv7.2-M-currents by TARP-γ2, ultimately resulting in increased susceptibility to seizures induced by GABAergic inhibition with pentylenetetrazol. Moreover, knock-in mice with this TARP-γ2 variant displayed decreased susceptibility to seizures induced by the M-channel blocker XE991 and reduced response to retigabine in the subiculum, supporting diminished M-currents in the presence of the mutant form of TARP-γ2.

Balanced neuronal excitability is particularly relevant in cerebral regions responsible for brain-wave oscillations and seizure control, which can act as backup brakes to suppress or facilitate convulsions. Interestingly, we found co-expression of TARP-γ2 and Kv7.2 in brain regions implicated in seizure suppression. Strong co-enrichment of TARP-γ2 and Kv7.2 was observed in the cerebellum, particularly in the cerebellar molecular layer, where stimulation of the cerebellar deep fastigial nucleus can block temporal lobe seizures^68,69^. Thus, the regulation of Kv7.2-mediated currents by TARP-γ2, and consequently synaptic excitability, may be of importance in Purkinje cell dendrites. Additionally, strong co-labeling of TARP-γ2 and Kv7.2 was found in the pallidum ventral subregions and the substantia nigra pars reticulata. Of note, deep stimulation of the ventral subregions of the pallidum attenuates pilocarpine-induced seizures^70^, and inhibition of substantia nigra pars reticulata neurons suppresses seizures in epilepsy-prone genetically engineered rat models^71^. Another region with significant enrichment of Kv7.2 and TARP-γ2 is the subiculum in the hippocampal formation. Subicular pyramidal cells are burst-firing neurons that contribute to the oscillatory activity of this brain area. This region has been reported to be particularly active in cases of focal epileptic disorders^72^. The functional modulation of Kv7.2 by TARP-γ2 in these brain regions may ensure a suitable level of excitability to coordinate the appropriate neuronal output.

Similar to stargazer mice^50,73^, subicular pyramidal neurons in TARP-γ2 V143L mice exhibited a decrease in the mAHP following a burst. A major contribution of Kv7.2-mediated currents to the mAHP has been suggested as conditional deletion of Kv7.2, but not Kv7.3, from CA1 pyramidal neurons results in a striking decrease in the mAHP^23^. Hence, we propose that the decreased mAHP observed in TARP-γ2 V143L mice may result from impaired Kv7.2 homomeric currents due to defective regulation by the TARP-γ2 V143L variant.

### Type I TARP-Kv7.2 interaction as a molecular integrator of synaptic input and intrinsic excitability

A recent study by Li and colleagues^74^ revealed an autoregulatory feedback loop in which chronic inactivity induces a homeostatic widening of action potentials. This homeostatic regulation of neuronal excitability relies on the control of the alternative splicing of BK channels through molecular mechanisms similar to Hebbian plasticity. Despite the growing evidence of cross-talk between homeostatic and Hebbian plasticity, as well as between synaptic transmission regulation and neuronal excitability, the molecular players facilitating this integration remain elusive. The role of TARP-γ2 as an auxiliary protein of both AMPARs and M-channels suggests that TARPs may function as molecular integrators of plasticity, synaptic transmission, and intrinsic neuronal excitability. Specific molecular modifications could potentially act as switches, directing TARPs towards the regulation of either AMPARs or Kv7.2 channels. The molecular mechanisms underlying these switches, which are likely responsive to neuronal activity, remain to be identified and warrant further investigation.

Recent studies have begun to unveil the interplay between glutamatergic transmission and Kv7.2-M-channels in the ventral hippocampus. The sustained antidepressant effects of ketamine were found to be potentiated by retigabine and to depend on increased M-currents through upregulation of Kv7.2 mRNA, but not Kv7.3, in this brain region^75^. Ketamine also increases AMPAR function in the ventral hippocampus, which is crucial for its fast antidepressant effect. This effect relies on the rapid recruitment of TARP-γ8 to synaptic sites, anchoring more AMPAR at PSD95 clusters in a CaMKIIα-dependent manner^76^. Based on these studies and our findings, it is tempting to suggest that a TARP-mediated mechanism, through the upregulation of Kv7.2 activity, may be involved in recruiting input-inhibition mechanisms to counteract ketamine-enhanced AMPAR function, thus preventing excessive neuronal activity.

Altogether, our findings establish a molecular link between synaptic function and intrinsic excitability. The concurrent modulation of AMPARs and Kv7.2 channels by TARPs at the dendritic level provides a mechanism for maintaining synaptic signals within physiological limits, particularly following synaptic plasticity. The slow braking activity of Kv7.2, promoted by TARPs, limits the depolarizing effects of AMPAR activation in a time-precise manner. In this scenario, the neuronal computing capacity is expanded while preserving both Hebbian and homeostatic net capabilities ready for further activity changes, thus facilitating complex cognitive processes. The pathways involved in such mechanisms require further investigation, as each type I TARP possesses differential regulatory effects, despite their convergent role in potentiating Kv7.2 function.

## STAR⍰Methods

### Key resources table

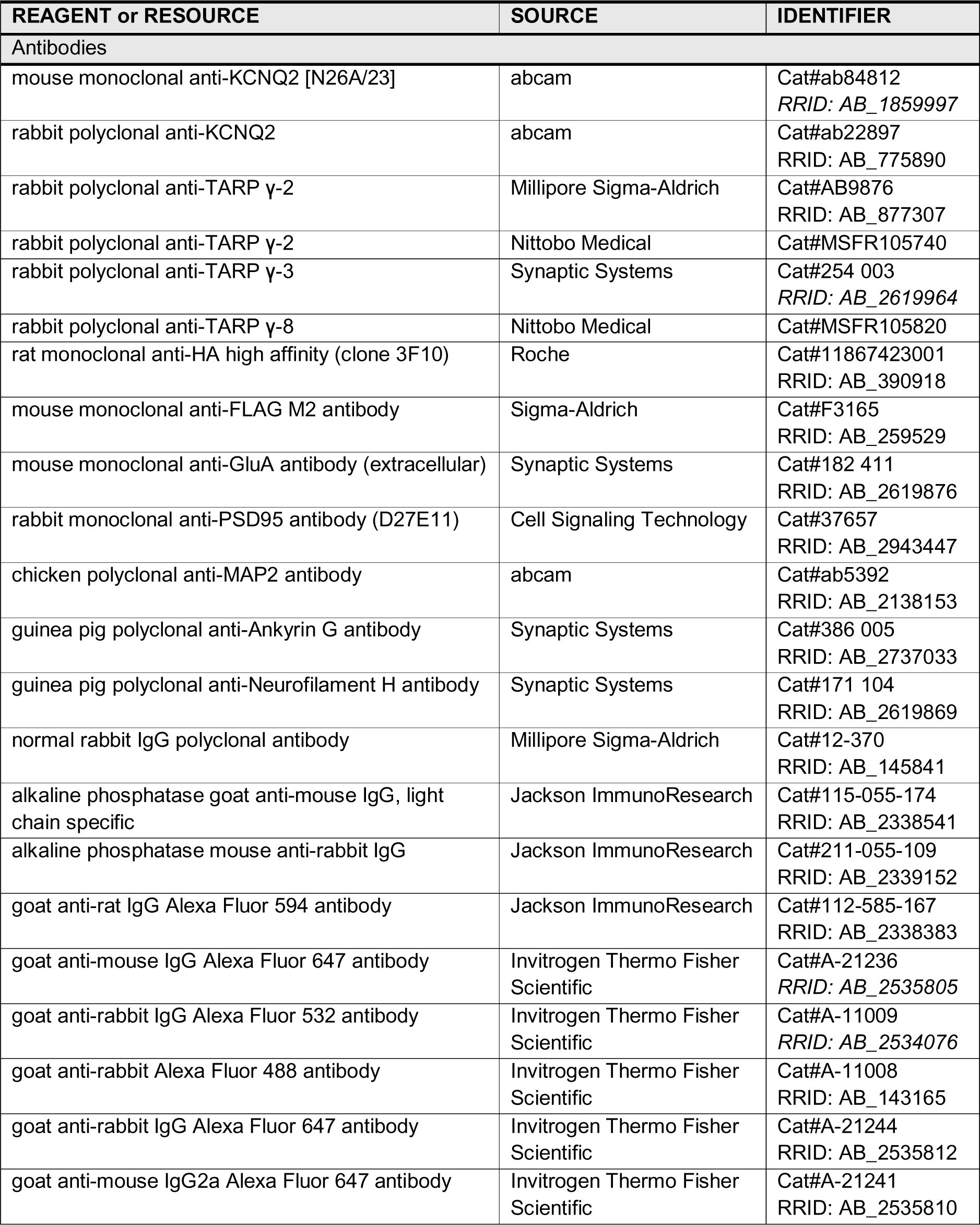
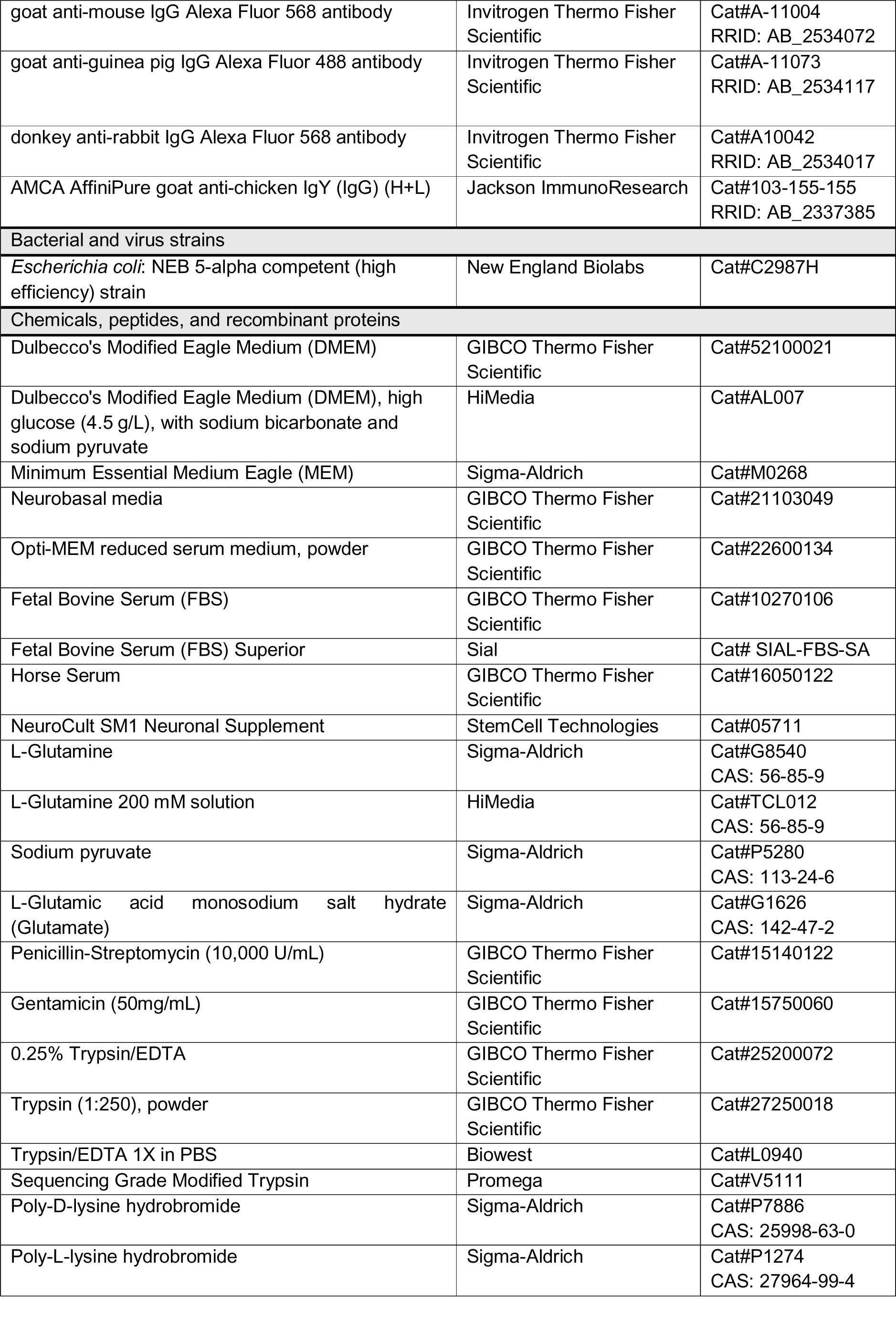
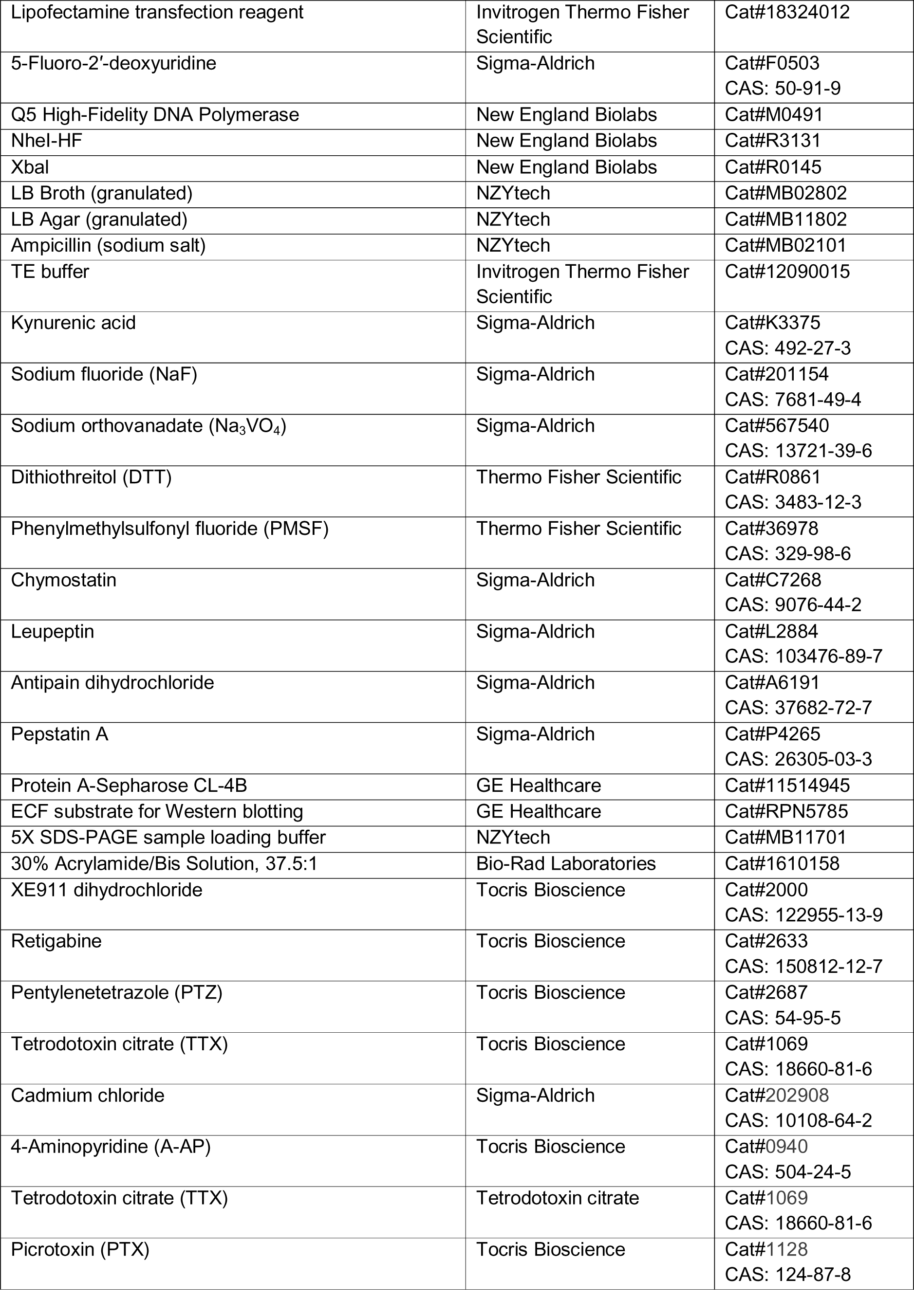
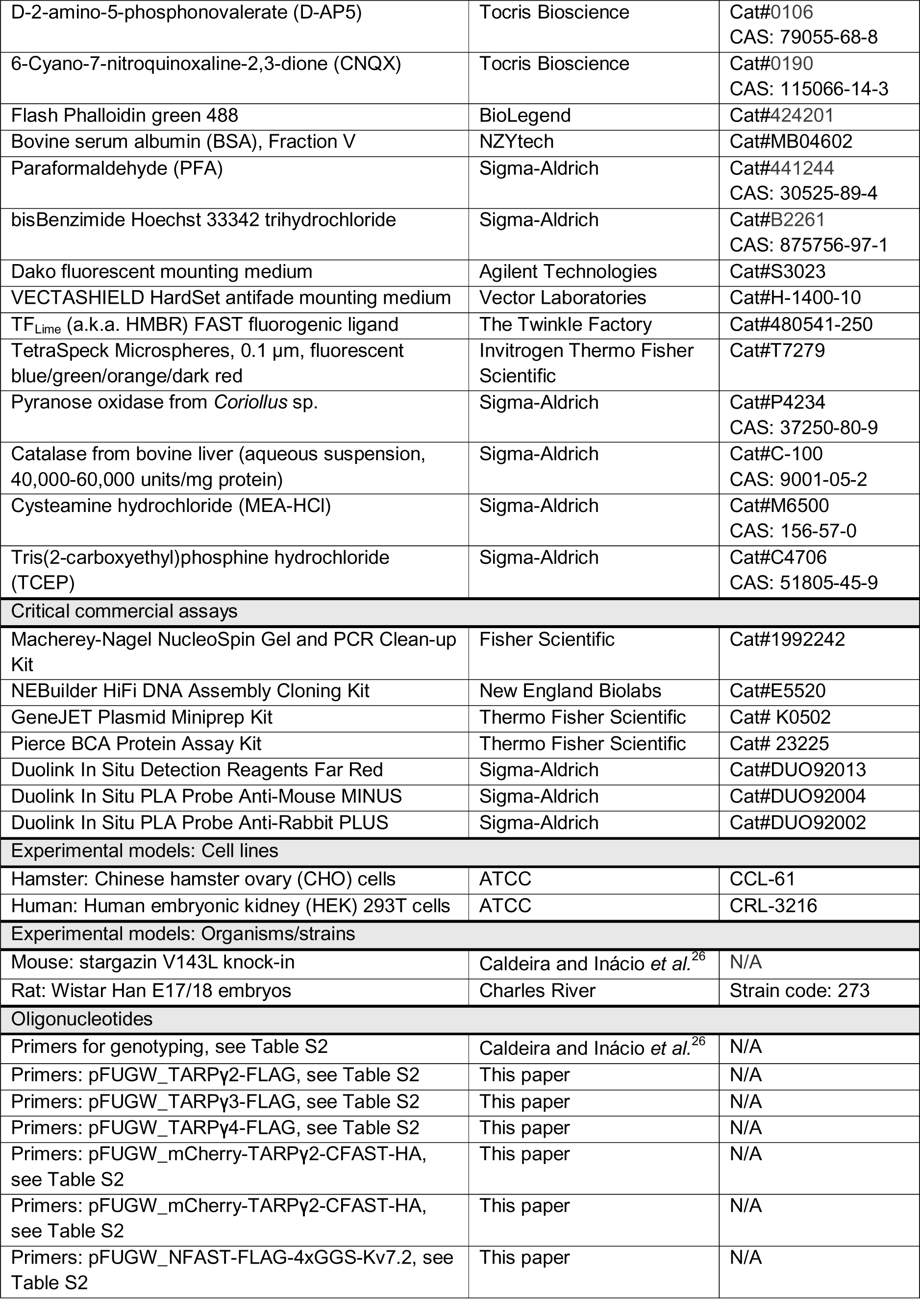
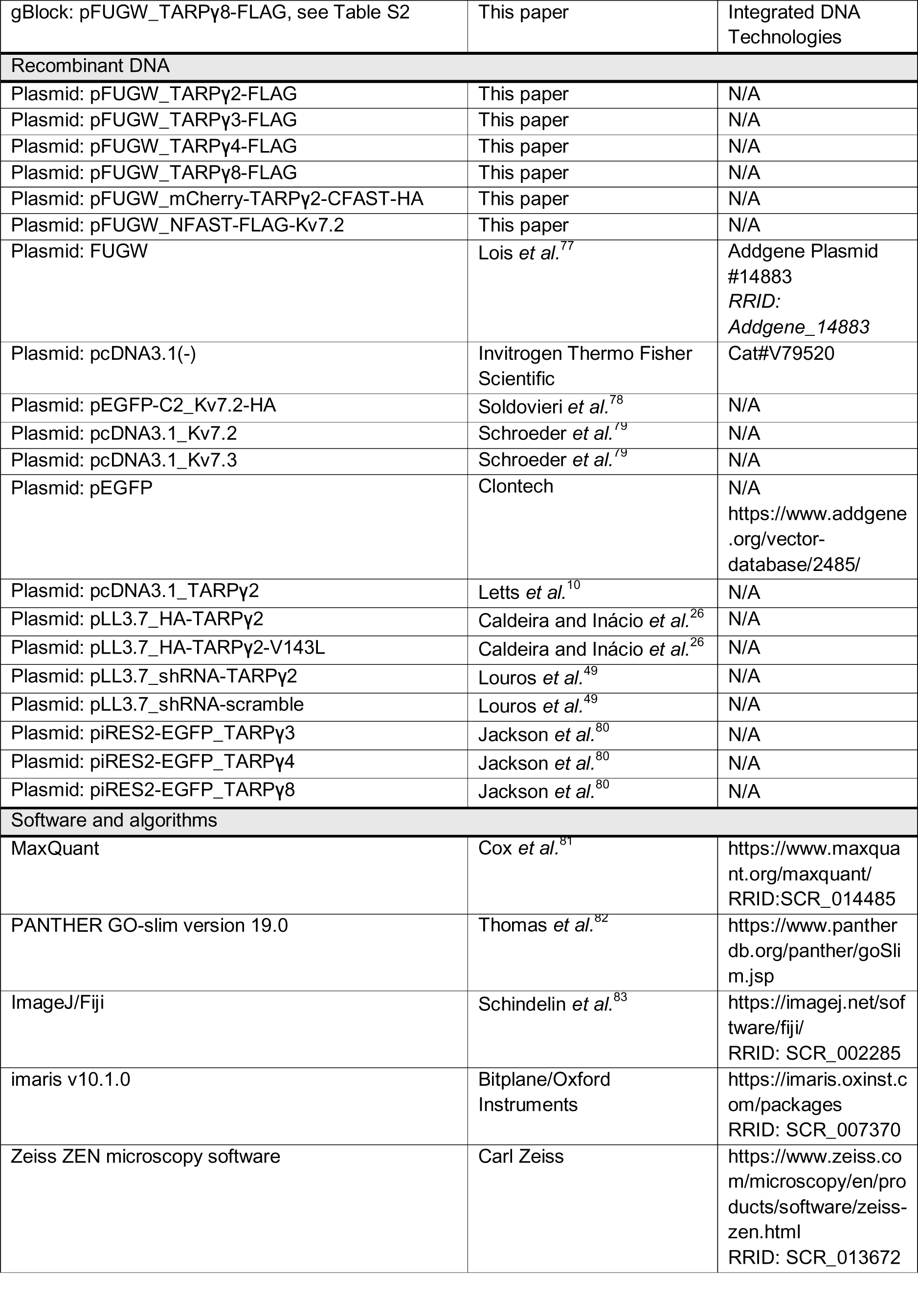
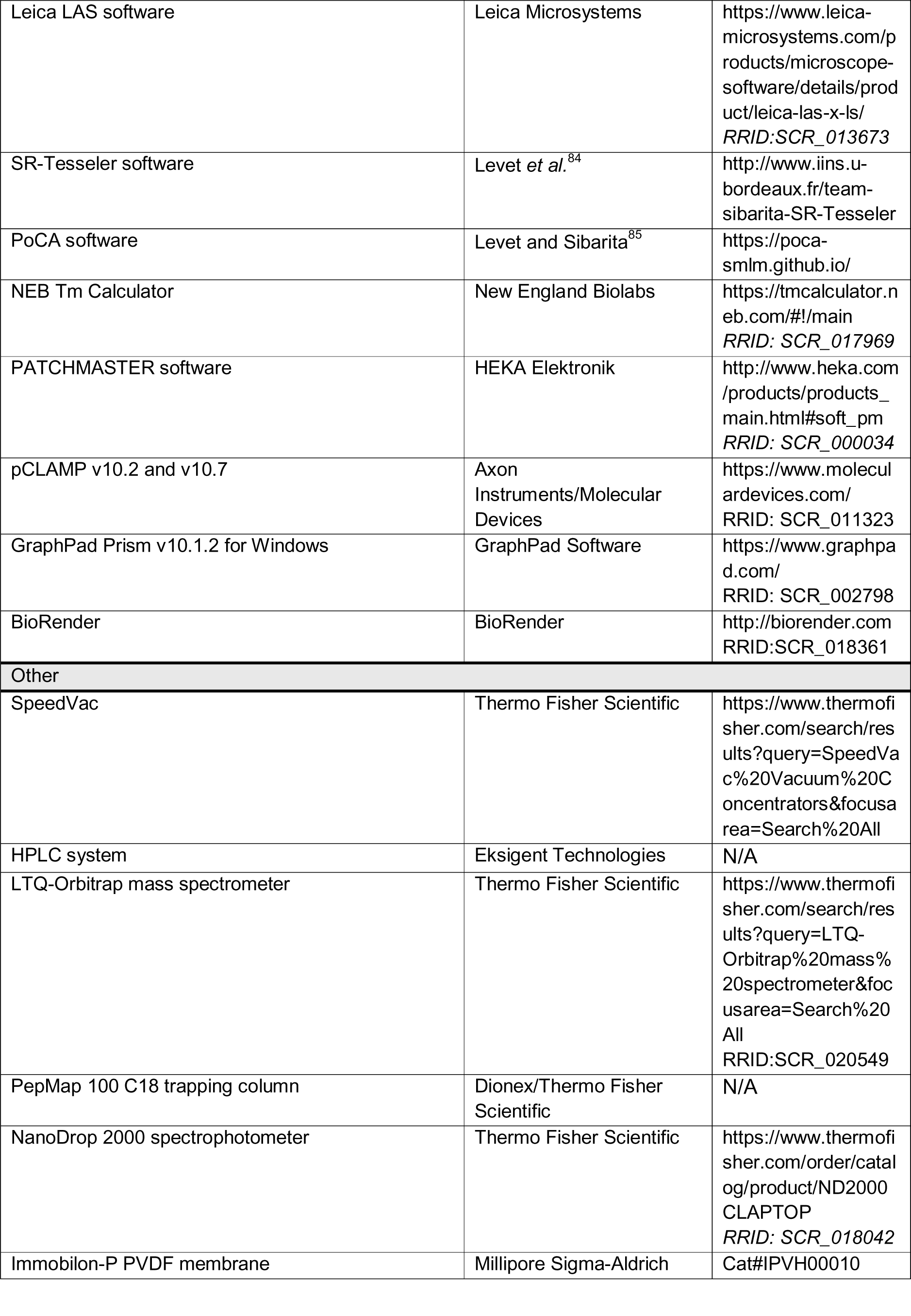
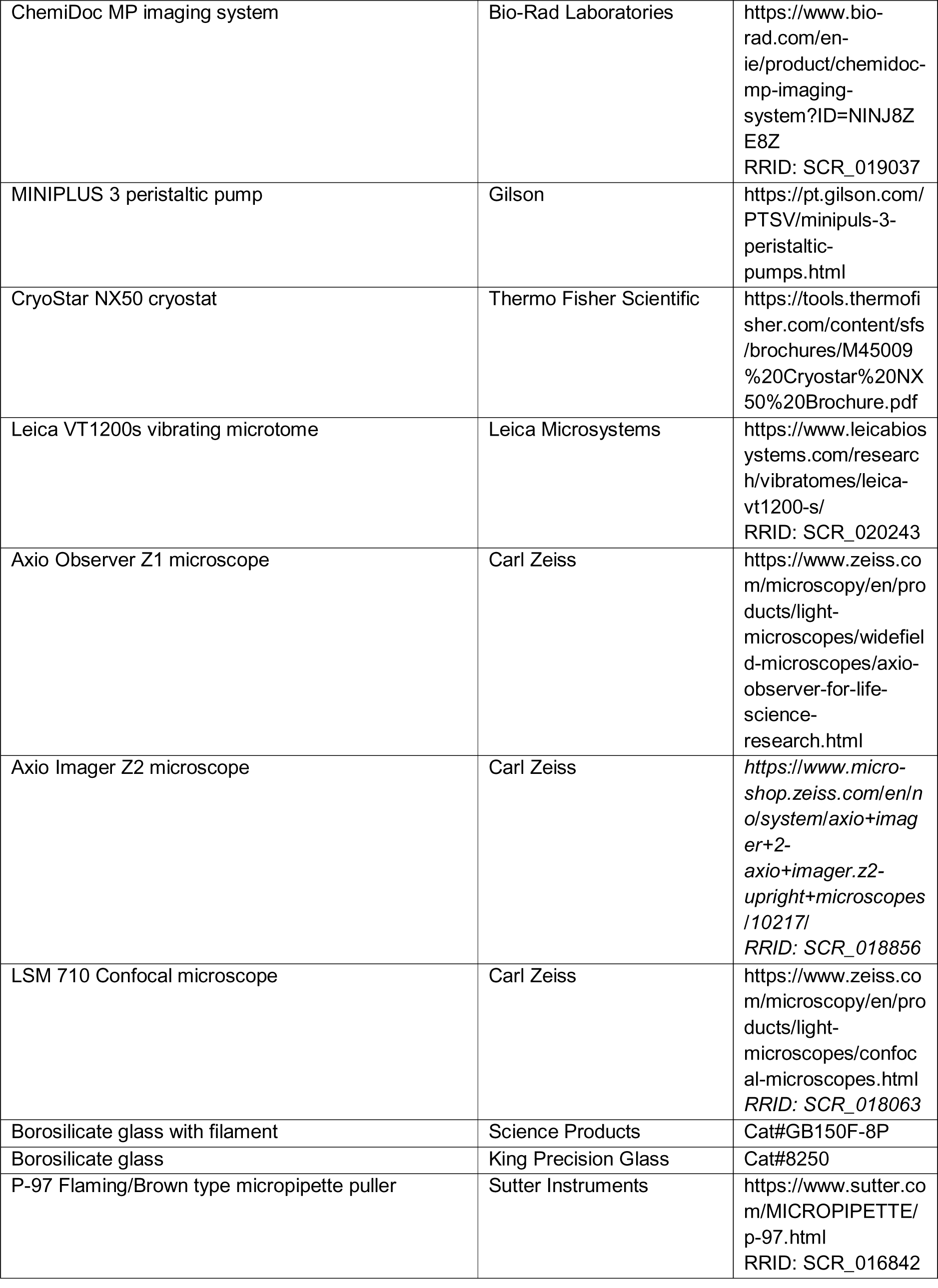
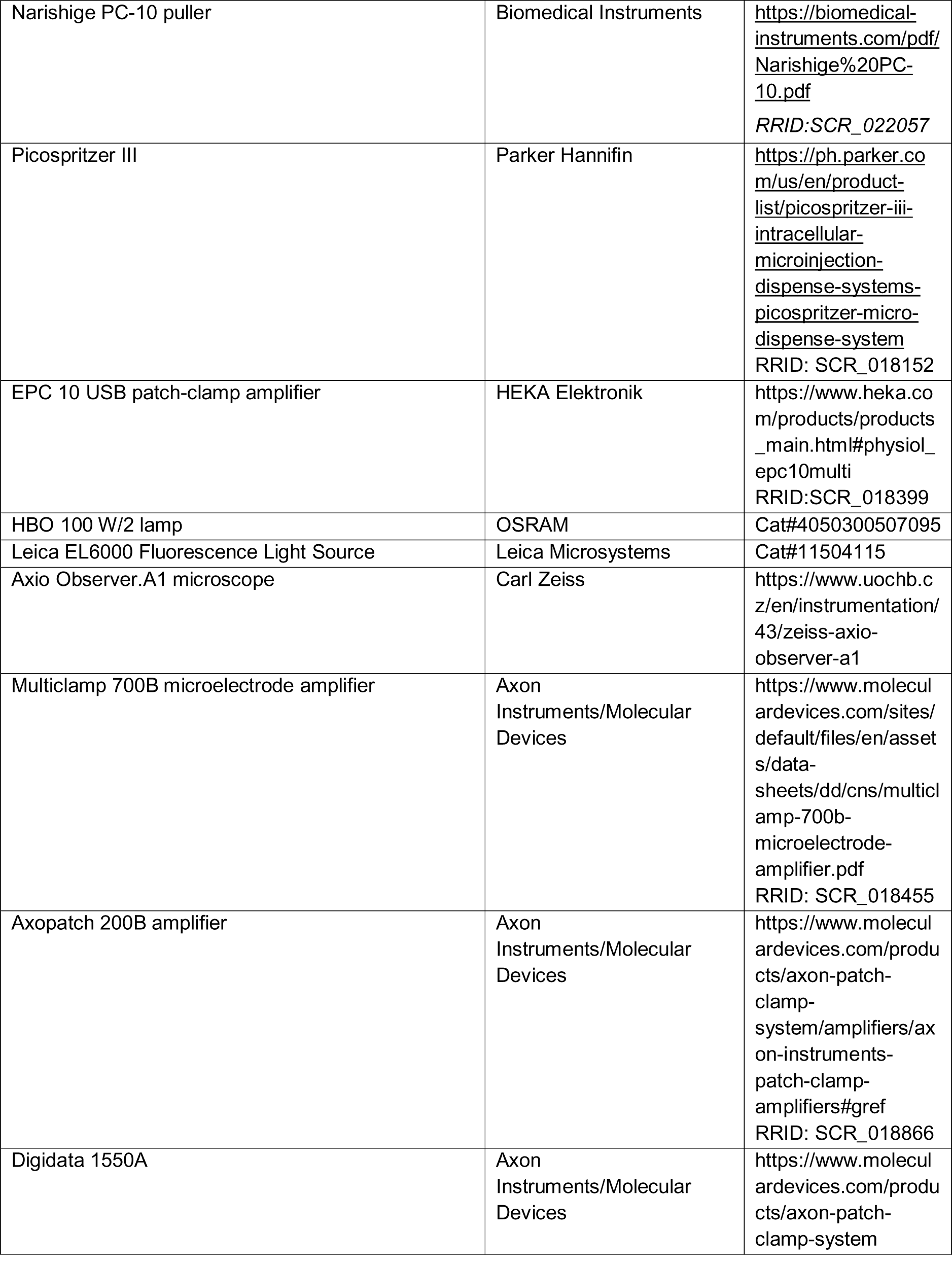
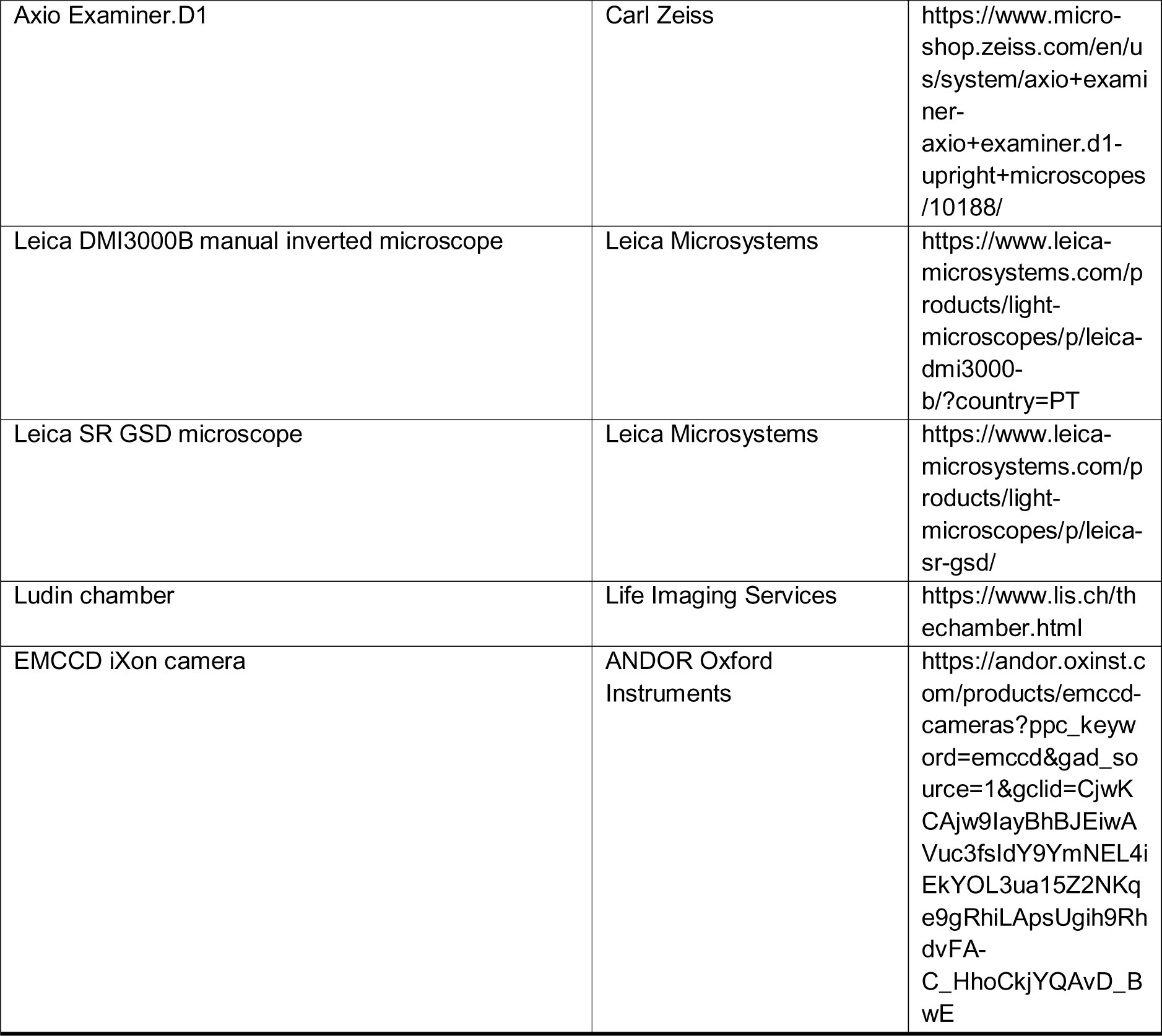

### Resource availability

#### Lead contact

Further information and requests for resources and reagents should be directed and will be fulfilled by the lead contact, Ana Luísa Carvalho.

#### Materials availability

Plasmids generated in this study are available from lead contact upon request.

#### Data and code availability

All data reported in this paper will be shared by the lead contact upon request. This paper does not report original code.

Any additional information required to reanalyze the data reported in this paper is available from the lead contact upon request.

## Experimental model and study participant details

### Cell line culture

#### Chinese hamster ovary cells (CHO)

CHO cells (ATCC, #CCL-61) were grown in DMEM (HiMedia, #AL007), supplemented with 10% inactivated FBS (Sial, #SIAL-FBS-SA), 1% penicillin/streptomycin (GIBCO Thermo Fisher Scientific, #15140122), and 1% L-glutamine (HiMedia, #TCL012), at 37°C in a humidified atmosphere with 5% CO_2_. Cells were split when reaching 80-90% confluence by enzymatic digestion using trypsin/EDTA (Biowest, #L0940) and were maintained for a maximum of 100 passages. For electrophysiological recordings, cells were plated at a 70% confluence on 40-mm plates containing 25 mm glass coverslips coated with 0.1 mg/mL poly-L-lysine (Sigma-Aldrich, #P1274) and transfected 24 hours later.

#### Human embryonic kidney 293T cells (HEK 293T)

HEK 293T cells (ATCC, #CRL-3216) were cultured in DMEM (GIBCO Thermo Fisher Scientific, #52100021) supplemented with 10% inactivated FBS (GIBCO Thermo Fisher Scientific, #10270106), and 1% penicillin/streptomycin (GIBCO Thermo Fisher Scientific, #15140122), at 37°C in a humidified atmosphere with 5% CO_2_. Cells were split when reaching 80-90% confluence by enzymatic digestion using 0.25% trypsin/EDTA (GIBCO Thermo Fisher Scientific, #25200072) and were maintained for a maximum of 35 passages. Cells were plated on 6-well plates for immunoprecipitation assays or on 18 mm glass coverslips coated with 0.1 mg/mL poly-D-lysine (Sigma-Aldrich, #P7886) for immunofluorescence assays, and they were transfected when reaching 50-60% confluence.

### Primary hippocampal and cortical neuronal cultures

Primary hippocampal and cortical neuronal cultures were prepared from E17/18 Wistar rat embryos. Briefly, hippocampi and cortices were dissected and collected in Ca^2+^- and Mg^2+^-free Hanks’ balanced salt solution (HBSS; 5.36 mM KCl, 0.44 mM KH_2_PO_4_, 137 mM NaCl, 4.16 mM NaHCO_3_, 0.34 mM Na_2_HPO_4_, 5 mM glucose, 1 mM sodium pyruvate, 10 mM HEPES, and 0.001% phenol red). Tissues were then digested with 0.02% trypsin (GIBCO Thermo Fisher Scientific, #27250018) in HBSS at 37°C for 12-15 minutes. After extensive washing with HBSS to remove trypsin, tissues were mechanically dissociated, and neurons were plated in MEM (GIBCO Thermo Fisher Scientific, #10270106) supplemented with 10% horse serum (GIBCO Thermo Fisher Scientific, #16050122), 0.6% glucose, and 1 mM sodium pyruvate (Sigma-Aldrich, #P5280) on 18 mm glass coverslips coated with 0.1 mg/mL poly-D-lysine (Sigma-Aldrich, #P7886). Cells were plated at an approximate density of 90,000 cells/cm^2^ for immunoprecipitation assays, 12,000 cells/cm^2^ for imaging, and 52,000 cells/cm^2^ for electrophysiology experiments. After cell attachment to the surface, approximately 2 hours after plating, the coverslips were inverted onto a confluent support layer of glia cells on 60 mm Petri dishes^86^ and maintained in Neurobasal medium (GIBCO Thermo Fisher Scientific, #21103049) supplemented with 1:50 NeuroCult SM1 neuronal supplement (StemCell Technologies, #05711), 0.5 mM L-glutamine (Sigma-Aldrich, #G8540), and 0.12 mg/mL gentamicin (GIBCO Thermo Fisher Scientific, #15750060), at 37°C in a humidified atmosphere with 5% CO_2_. In the case of hippocampal neuronal cultures, 25 µM glutamate (Sigma-Aldrich, #G1626) was also added to the supplemented Neurobasal medium. Three days after plating, cultures were treated with 10 µM 5-Fluoro-2′-deoxyuridine (Sigma-Aldrich, #F0503) to prevent the overgrowth of glial cells. Cells were fed every five days by replacing one-third of the culture medium with fresh supplemented Neurobasal medium without glutamate. When required, neuronal depolarization was induced by adjusting the KCl concentration of the conditioned culture medium to 15 mM 24 hours before use.

### Mice

All procedures involving animals adhered strictly to the guidelines established by the European Union Directive 2010/63/EU. All experiments were approved by the institutional animal welfare body (ORBEA) and the national competent authority (DGAV). Stargazin V143L mice were generated in our laboratory as previously described^26^, and were bred and maintained in the animal facility at the Faculty of Medicine at the University of Coimbra. Mice were housed in a 12-hour light/dark cycle at 22°C and 60% humidity. Food and water were provided *ad libitum*. Genotyping was performed by PCR from mouse ear or tail DNA using a forward primer for the WT allele (AAGGGACCCTCCGTCCTCTC) and a forward primer for the mutant allele (GGGCCCGGTGCAATACACGC), while the same reverse primer was used for both reactions (CATCGGGCATGGATCCTCAGTTC). Littermate WT mice were used as controls for all experiments. 8-week-old animals were used for electrophysiology recordings, and 10-12-week-old animals were used for immunohistochemistry experiments and seizure susceptibility testing. Both male and female mice were used for immunohistochemistry, and male mice were used for all other experiments. All tests and quantifications were performed blindly to the genotype. Sample size estimates were based on previous literature, and no randomization was applied.

## Method details

### Immunoprecipitation-mass spectrometry analysis

Total cell lysates were prepared from 11 days *in vitro* (DIV) high-density cultured cortical neurons. The cell culture medium was removed, and the cells were washed with ice-cold PBS, and scraped and resuspended in ice-cold TEEN buffer (25 mM Tris-HCl pH 7.4, 150 mM NaCl, 1 mM EDTA, 1 mM EGTA, 1% Triton X-100) supplemented with a cocktail of phosphatase and protease inhibitors (5 mM NaF, 0.1 mM Na_3_VO_4_, 1 mM okadaic acid, 1 mM DTT, 0.2 mM PMSF, 1 µg/mL CLAP – chymostatin, leupeptin, antipain, pepstatin). The cell lysates were kept on ice for 10 minutes before being sonicated for 45 seconds (4 pulses of 5 seconds) using an ultrasonic probe. After centrifugation at 21,100 g for 10 minutes at 4°C, the supernatant was collected, and the protein concentration was quantified using the bicinchoninic acid (BCA) assay (Thermo Fisher Scientific, #23225), following the protocol provided by the manufacturer. For TARP-γ2 immunoprecipitation, cell lysates were diluted with supplemented TEEN buffer to a final concentration of 2 µg/µL (approximately 1.6 mg of protein) and pre-cleared by incubation with 30 µL of protein-A Sepharose (GE Healthcare, #11514945) for 1 hour at 4°C under rotation. After centrifugation at 3,000 g for 1 minute at 4°C, the supernatant was collected and divided into two tubes. One was incubated with 2 µg of rabbit polyclonal Anti-TARP γ-2 (Millipore Sigma-Aldrich, #AB9876), and the other with 2 µg of species-matched non-specific IgGs (Millipore Sigma-Aldrich, #12-370), overnight at 4°C under rotation. After that, 80 µL of protein-A Sepharose beads were added, and the samples were incubated for 2 hours at 4°C under rotation. Following centrifugation at 3,000 g for 1 minute at 4°C, 30 µL of the supernatant was collected to evaluate the efficiency of the immunoprecipitation. An equal volume of 2X SDS-PAGE sample loading buffer (diluted from 5X SDS-PAGE sample loading buffer; NZYTech, #MB11701) was added to each sample. The Sepharose beads were washed six times: three washes with TEEN buffer, two with TEEN buffer supplemented with 1% Triton X-100, and one with TEEN buffer supplemented with 150 mM NaCl. This was followed by a final centrifugation at 3,000 g for 1 minute at 4°C. The supernatant was discarded, and proteins were eluted from the beads using 40 µL of 2X SDS-PAGE sample loading buffer (diluted from NZYTech 5X SDS-PAGE sample loading buffer). Samples were further incubated with 8 µL of 30% acrylamide (BioRad, #1610158) for 30 minutes at room temperature to alkylate cysteinyl residues and then resolved by SDS-PAGE in Tris-glycine-SDS buffer (25mM Tris, 192mM glycine, 0.1% SDS, pH 8.3) on an 11% polyacrylamide gel. Polyacrylamide gels were stained for 2 hours at room temperature with colloidal Coomassie blue, as previously described^87^, washed with water, and each gel lane was cut into five slices using a clean scalpel. Gel pieces were subsequently subjected to tryptic digestion as described elsewhere^88^ Briefly, the gel pieces were sequentially incubated with 50% acetonitrile in 50 mM NH_4_CO_3_, 100% acetonitrile, and 50 mM NH_4_CO_3_. The destaining cycle was repeated once. After dehydration in 100% acetonitrile, gel fragments were dried in a SpeedVac (Thermo Fisher Scientific), digested with 10 mg/mL trypsin (Promega, #V5111) in 50 mM NH_4_HCO_3_ overnight at 37°C. The peptides were extracted twice from the gel pieces with 200 µL of 50% acetonitrile in 0.1 % trifluoroacetic acid. The supernatants were collected, dried in a SpeedVac, and stored at - 20°C until further use. The peptides were re-dissolved in 20 µL 0.1% acetic acid and analyzed by liquid chromatography coupled with mass spectrometry (LC-MS/MS). Samples were injected into an LTQ-Orbitrap mass spectrometer (Thermo Electron, Thermo Fisher Scientific) equipped with an HPLC system (Eksigent Technologies). Peptides trapped on a 5 mm PepMap 100 C18 column (300 µm ID, 5 µm particle size; Dionex, Thermo Fisher Scientific) and then analyzed on a 200 mm Alltima C18 home-made column (100 µm ID, 3 µm particle size). Peptides were separated at a flow rate of 400 nL/min with a linear 40-minute gradient from 5% solvent A (5% acetonitrile, 94.9% H_2_O, 0.1% acetic acid) to 40% solvent B (95% acetonitrile, 4.9% H_2_O, 0.1% acetic acid). The mass spectrometer was operated in a data-dependent mode with a single MS full-scan survey (m/z range 330-2000), followed by MS/MS experiments of the five most abundant ions.

### Generation of DNA constructs

The engineering of the pFUGW_TARP-FLAG constructs consisted of inserting a FLAG-tag in the C-terminal tail of each TARP at the 304^th^ nucleotide, similar to the mCherry insertion in TARP γ-2 described by Sainlos and colleagues^89^. Except for TARP γ-8, primers (see Table S2) were designed to produce two DNA inserts: one containing the N-terminal and transmembrane domains coding sequence and the other containing the C-terminal coding sequence. In the case of TARP γ-8, a synthetic oligomer containing overhangs allowing for Gibson assembly (gBlock sequence in Table S2, Integrated DNA Technologies) was used. For the pFUGW_mCherry-TARPγ2-CFAST-HA construct, the CFAST11 sequence^39^ was inserted into the C-terminal tail of TARP γ-2 at the 304^th^ nucleotide, and mCherry and T2A sequences were cloned upstream of the TARP γ-2 sequence. An HA-tag was also inserted after CFAST11, before the C-terminal of TARP γ-2. Three DNA inserts were produced using the primers listed in Table S2. The pFUGW_NFAST-FLAG-4xGGS-Kv7.2 construct was engineered to insert the NFAST sequence^39^ followed by a FLAG-tag and a flexible spacer of four GSS repeats in the N-terminal of Kv7.2, cloned upstream of the Kv7.2 sequence similar to the EGFP insertion described by Soldovieri and colleagues^90^. Two DNA inserts were produced using the primers listed in Table S2. All DNA inserts were generated via PCR using Q5 High-Fidelity DNA Polymerase (New England Biolabs, #M0491), with 1 µg template DNA and 10 µM of each forward and reverse primer. Cycling parameters were an initial denaturation at 98°C for 30 seconds, followed by 35 amplification cycles of denaturation at 98°C for 10 seconds, primer annealing at 58-72°C for 60 seconds, and extension at 72°C for 30 seconds, with a final extension step at 72°C for 120 seconds. The primer annealing temperatures were defined based on predictions using the NEB Tm Calculator (New England Biolabs). The resulting amplicons were run on a 1% agarose gel at 90V for 40 minutes and then purified using the Macherey-Nagel NucleoSpin Gel and PCR Clean-up Kit (Fisher Scientific, #1992242), following the manufacturer’s instructions. Inserts were cloned into the FUGW plasmid (Addgene, #14883) via Gibson assembly^91^. The EGFP sequence was excised during the plasmid linearization step using the NheI-HF (New England Biolabs, #R3131) and XbaI (New England Biolabs, #R0145) enzymes. For Gibson assembly reactions, 100 ng of plasmid DNA was incubated with insert DNA at a 1:2 ratio, in 10 µL of 2X concentrated NEBuilder HiFi DNA Assembly Master Mix (New England Biolabs, #E5520). The final volume of the reaction was adjusted to 20 µL with nuclease-free water and incubated at 50°C for 1 hour. NEB 5-alpha competent *E. coli* strain (New England Biolabs, # C2987H) was used for subcloning. Briefly, 3 µL of the Gibson assembly reaction product was added to 50 µL of bacterial suspension and incubated on ice for 30 minutes, followed by 30 seconds at 42°C and a further 2 minutes on ice. Subsequently, 950 µL of SOC outgrowth medium (New England Biolabs, #E5520) was added and the bacteria were incubated at 37°C for 1 hour with agitation (250 rpm). At the end of the incubation, the bacteria were centrifuged at 21,100 g for 1 minute, and the pellet was resuspended in LB broth (NZYtech, #MB02802). Bacteria were then spread on LB agar (NZYtech, #MB11802) plates with 0.5 mg/mL ampicillin (NZYtech, #MB02101) and grown at 30°C for 20 hours. DNA constructs were purified from selected colonies using the GeneJET Plasmid Miniprep Kit (Thermo Fisher Scientific, #K0502), and DNA was eluted in TE buffer (Invitrogen Thermo Fisher Scientific, #12090015). The DNA concentration was measured by absorbance at 260 nm using a NanoDrop 2000 (Thermo Fisher Scientific), and the plasmids were stored at - 20°C. All DNA constructs were validated using Sanger sequencing (Eurofins Genomics).

### Cell transfection

#### Transfection of CHO cells

For electrophysiological recordings, CHO cells were transfected with 4 µg of plasmid DNA using the lipocationic reagent Lipofectamine (Invitrogen Thermo Fisher Scientific, #18324012). Cells were co-transfected with pcDNA3.1_Kv7.2 and either pcDNA3.1(-) empty vector, pcDNA3.1_TARPγ2, piRES2-EGFP_TARPγ3, piRES2-EGFP_TARPγ4, or piRES2-EGFP_TARPγ8, at different transfection ratios, totaling 3.6 µg of plasmid DNA. These plasmids were mixed with 0.4 µg of pEGFP, which contains the gene for Enhanced Green Fluorescent Protein (EGFP) used as a transfection marker. Briefly, plasmid DNAs and Lipofectamine were separately diluted in 250 µl of Opti-MEM medium (GIBCO Thermo Fisher Scientific, #22600134), mixed, and incubated for 20 minutes at room temperature to allow the formation of Lipofectamine-DNA complexes. Subsequently, 1 mL of Opti-MEM was added to the precipitates, and the total volume of the mixture was used to replace the DMEM medium in the culture dish. After 5 hours of incubation, the medium containing the precipitate was replaced with fresh complete DMEM medium.

For Kv7.2/Kv7.3 heteromeric current recordings, cells were co-transfected with pcDNA3.1_Kv7.2 and pcDNA3.1_Kv7.3, along with either pcDNA3.1(-) empty vector or pcDNA3.1_TARPγ2, at a 1:1:5 ratio, as described.

#### Transfection of HEK 293T cells

For immunoprecipitation and imaging assays, HEK 293T cells were transfected with 10 µg or 4 µg of plasmid DNA, respectively, using the calcium phosphate co-precipitation method. Briefly, plasmid DNA was diluted in TE buffer (Invitrogen Thermo Fisher Scientific, #12090015), and a CaCl_2_ solution (2 M CaCl_2_) was added dropwise to reach a final concentration of 246 mM, facilitating the formation of precipitates. These precipitates were subsequently combined with an equal volume of 2X concentrated HEPES-buffered saline (280 mM NaCl, 50 mM HEPES, 1.5 mM Na_2_HPO_4_, pH 7.1). The precipitate solution was added to the cells and incubated at 37°C for 4-6 hours in a humidified atmosphere with 5% CO_2_. Afterwards, the cell culture medium was replaced with fresh medium.

For immunoprecipitation assays, cells were co-transfected with pEGFP-C2_Kv7.2-HA and either pcDNA3.1_TARPγ2, piRES2-EGFP_TARPγ3 or piRES2-EGFP_TARPγ8, at a 1:1 ratio. Cells were lysed 48 hours after transfection.

For immunostaining, cells were co-transfected with pEGFP-C2_Kv7.2-HA and either pcDNA3.1(-) empty vector, pFUGW_TARPγ2-FLAG, FUGW_TARPγ3-FLAG, FUGW_TARPγ4-FLAG or FUGW_TARPγ8-FLAG, at a 1:1 ratio. Cells were fixed and immunostained 24 hours after transfection.

For the SplitFAST assay, cells were co-transfected with pFUGW_mCherry-TARPγ2-CFAST- HA and pFUGW_NFAST-FLAG-4xGGS-Kv7.2, at a 1:1 ratio. As a negative control, cells were transfected with each of the plasmids alone, and the total amount of DNA was kept constant using the pcDNA3.1(-) empty vector. Imaging was performed 24 hours after transfection. To assess the possible effects of the protein tags’ position on protein localization, cells were transfected with either TARPγ2-CFAST-HA, pcDNA3.1_TARPγ2, pFUGW_NFAST-FLAG-4xGGS-Kv7.2 or pcDNA3.1_Kv7.2, fixed, and immunostained 24 hours after transfection.

#### Transfection of cortical neurons

Rat cortical neurons were transfected with 3 µg of plasmid DNA at DIV 11 using the calcium phosphate co-precipitation method^92^. Plasmid DNA was diluted in TE buffer (1 mM Tris-HCl pH7.3, 1 mM EDTA), and a CaCl_2_ (2.5 mM CaCl^2^ in 10 mM HEPES) was added dropwise to reach a final concentration of 250 mM, facilitating the formation of precipitates. These precipitates were subsequently combined with an equal volume of 2X concentrated HEPES-buffered saline (274 mM NaCl, 42 mM HEPES, 1.4 mM Na_2_HPO_4_, 10 mM KCl, 11 mM glucose, pH 7.2). Neurons were transferred to 12-well plates with 200 µL of conditioned culture medium supplemented with 2 M kynurenic acid (Sigma-Aldrich, #K3375). Then, 100 µL of the precipitate solution was added to each well, and cells were incubated for 1-2 hours at 37°C in a humidified atmosphere with 5% CO_2_. Afterwards, the remaining precipitates were dissolved by incubating the neurons in fresh Neurobasal medium supplemented with 2 mM kynurenic acid for 15 minutes at 37°C in a humidified atmosphere with 5% CO_2_. Lastly, coverslips were transferred to the original culture dish with glia cells and kept for four days before being used for electrophysiological recordings or being fixed for imaging experiments. For electrophysiological recordings, cells were transfected with either pLL3.7_shRNA-scramble or pLL3.7_shRNA-TARPγ2. For dSTORM super-resolution imaging, neurons were transfected with either pLL3.7_shRNA-scramble, pLL3.7_shRNA-TARPγ2 or pLL3.7_HA-TARPγ2.

### Co-immunoprecipitation assays

HEK 293T cell lysates were prepared 48 hours after transfection. The cell culture medium was removed, and the cells were scraped and resuspended in ice-cold TEEN buffer (25 mM Tris-HCl pH 7.4, 150 mM NaCl, 1 mM EDTA, 1 mM EGTA, 1% Triton X-100) supplemented with a cocktail of phosphatase and protease inhibitors (5 mM NaF, 0.1 mM Na_3_VO_4_, 1 mM DTT, 0.2 mM PMSF, 1 µg/mL CLAP – chymostatin, leupeptin, antipain, pepstatin). The cell lysates were kept on ice for 10 minutes before being sonicated for 70 seconds (7 pulses of 5 seconds) using an ultrasonic probe. After centrifugation at 21,100 g for 10 minutes at 4°C, the supernatant was collected, and the protein concentration was quantified using the bicinchoninic acid (BCA) assay (Thermo Fisher Scientific, #23225), following the protocol provided by the manufacturer. For type I TARP immunoprecipitation, 500 µg of protein was incubated with 2 µg of either rabbit polyclonal Anti-TARP γ-2 (Millipore Sigma-Aldrich, #AB9876), rabbit polyclonal Anti-TARP γ-3 (Synaptic Systems, #254 003), rabbit polyclonal Anti-TARP γ-8 (Nittobo Medical, #MSFR105820), or normal rabbit polyclonal IgG (Millipore Sigma-Aldrich, #12-370) antibodies for 2 hours at 4°C under rotation. After that, 50 µL of protein-A Sepharose (GE Healthcare, #11514945) was added, and the samples were incubated overnight at 4°C under rotation. The samples were then centrifuged at 100 g for 1 minute at 4°C, and the flowthrough was collected. An equal volume of 2X concentrated denaturing buffer (62.5 mM Tris-HCl pH 6.8, 10% glycerol, 2% SDS, 0.01% bromophenol blue, 5% β-mercaptoethanol) was added to each sample. The pellet containing the Sepharose beads was washed first with 1 mL of TEEN buffer, followed by a wash with TEEN buffer supplemented with 150 mM NaCl, and then resuspended in 50 µL of 2X concentrated denaturing buffer. Samples were resolved by SDS-PAGE in Tris-glycine-SDS buffer (25 mM Tris, 192 mM glycine, 0.1% SDS, pH 8.3) on an 11% polyacrylamide gel. Following electrophoresis, proteins were transferred overnight (40 V, 4°C) to a PVDF membrane (Millipore Sigma-Aldrich, #IPVH00010) in Tris-glycine buffer (25 mM Tris, 192 mM glycine, pH8.3) containing 20% methanol. The membranes were then blocked with 5% non-fat dry milk in TBS buffer (20 mM Tris, 137 mM NaCl, pH 7.6) with 0.1% Tween-20 (TBS-T) at room temperature for 1 hour. Next, the primary antibodies against either Kv7.2 (mouse monoclonal #ab84812, abcam; 1:1000), TARP γ-2 (rabbit polyclonal #AB9876, Millipore Sigma-Aldrich; 1:750), TARP γ-3 (rabbit polyclonal #254 003, Synaptic Systems; 1:1000), or TARP γ-8 (rabbit polyclonal #MSFR105820, Nittobo Medical; 1:1000) were diluted in blocking solution and incubated for 2 hours at room temperature. The membranes were then washed three times in TBS-T for 10 minutes each and incubated for 1 hour at room temperature with the appropriate alkaline phosphatase-conjugated secondary antibodies (goat anti-mouse IgG light chain specific #115-055-174 or mouse anti-rabbit IgG #211-055-109, Jackson ImmunoResearch) diluted 1:10,000 in blocking solution. Following three 10-minute washes with TBS-T, the membranes were developed with ECF alkaline phosphatase substrate (GE Healthcare, # RPN5785), and the protein bands were detected using a ChemiDoc MP imaging system (Bio-Rad Laboratories).

### Immunostaining and imaging

#### Immunostaining of rat cortical neurons

Low-density cortical neurons, 15 days in vitro (DIV), were fixed in 4% paraformaldehyde (PFA; Sigma-Aldrich, #441244) and 4% sucrose in phosphate-buffered saline (PBS; 137 mM NaCl, 2.7 mM KCl, 10 mM Na_2_HPO_4_, 1.8 mM KH_2_PO_4_, pH 7.4) for 15 minutes at room temperature. After washing with PBS, neurons were permeabilized with ice-cold 0.25% Triton X-100 in PBS for 5 minutes at 4°C, followed by three washes with PBS. Next, nonspecific staining was blocked by incubation with 10% bovine serum albumin (BSA; NZYtech, #MB04602) in PBS for 30 minutes at 37°C in a humidified chamber. Neurons were then incubated overnight at 4°C with the appropriate primary antibodies diluted in 3% BSA in PBS. The following primary antibodies were used: monoclonal mouse anti-Kv7.2 antibody (1:200; abcam, #ab84812), polyclonal rabbit anti-TARP γ-2 (1:500; Millipore Sigma-Aldrich, #AB9876), polyclonal chicken anti MAP2 (1:5000; abcam, #ab5392), polyclonal guinea pig anti-Ankyrin G (1:1000; Synaptic Systems, #386 005), and polyclonal guinea pig anti-Neurofilament H (1:1000; Synaptic Systems, #171 104). Neurons were then extensively washed with PBS and incubated with the appropriate secondary Alexa (goat anti-mouse IgG Alexa Fluor 568 #A-11004, goat anti-rabbit IgG Alexa Fluor 647 #A-21244, goat anti-guinea pig IgG Alexa Fluor 488 #A-11073, Invitrogen Thermo Fisher Scientific) or AMCA (goat anti-chicken IgY (IgG) #103-155-155, Jackson ImmunoResearch) antibodies, diluted 1:500 or 1:200 respectively, in 3% BSA in PBSA for 45 minutes at 37°C in a humidified chamber. When required, phalloidin (BioLegend, #424201), diluted at a ratio of 1:70, was added to the secondary antibody solution to stain actin filaments. Nuclei were visualized by staining with 1 µg/mL Hoechst 33342 (Sigma-Aldrich, #B2261) in PBS for 5 minutes at room temperature. Following extensive washing with PBS, coverslips were mounted with Dako fluorescent mounting medium (Agilent Technologies, #S3023). Images were captured using an Axio Observer Z1 microscope (Carl Zeiss) equipped with a Plan-Apochromat 63X oil-immersion objective (NA 1.4) and the Zeiss ZEN software.

For labelling surface AMPA receptors, neurons were incubated with a monoclonal mouse anti-GluA extracellular epitope antibody (1:100; Synaptic Systems, #182 411) prior to cell permeabilization. Briefly, after fixation, cells were washed with PBS and blocked in 10% BSA in PBS for 30 minutes at 37°C in a humidified chamber. Neurons were then incubated with the primary antibody, followed by incubation with the secondary antibody (1:500; goat anti-mouse IgG2a Alexa 647 #A-2124, Invitrogen Thermo Fisher Scientific), as described above. Subsequently, cells were permeabilized and stained for PSD95 (1:200, monoclonal rabbit anti-PSD95 #37657, Cell Signaling Technology; 1:500, goat anti-rabbit IgG Alexa 488 #A-11008, Invitrogen Thermo Fisher Scientific) and MAP2 (1:5000, polyclonal chicken anti MAP2 #ab5392, abcam; 1:200, AMCA goat anti-chicken IgY (IgG) #103-155-155, Jackson ImmunoResearch). Nuclei were stained with 1 µg/mL Hoechst 33342, and coverslips were mounted with Dako fluorescent mounting medium.

For dSTORM imaging, neurons were immunostained as already described using the primary mouse monoclonal anti-Kv7.2 antibody (1:200; abcam, #ab84812) alone or in combination with a rabbit polyclonal anti-TARP γ-2 antibody (1:200; Nittobo Medical, #MSFR105740), and the respective secondary antibodies (1:500; goat anti-mouse IgG Alexa 647 #A-21236, goat anti-rabbit IgG Alexa 532 #A-11009 Invitrogen Thermo Scientific). The cells were postfixed for 10 minutes at room temperature in 4% PFA and 4% sucrose in PBS to preserve the fluorescence signal and then kept in PBS until imaging.

#### Total and surface staining of Kv7.2 in HEK 293T cells

HEK 293T cells were fixed in 4% PFA and 4% sucrose in PBS for 15 minutes at room temperature, 24 hours after transfection with a dual-tagged Kv7.2 chimeric construct. This construct had an EGFP tag at the cytoplasmic N-terminus of Kv7.2, which was used to visualize and quantify the total expression levels of the protein. Additionally, the surface expression of Kv7.2 was evaluated by staining the extracellular HA-tag inserted in the loop connecting the transmembrane domains S1 and S2^78^ before cell permeabilization. Briefly, after washing with PBS, cells were blocked in 10% BSA in PBS for 30 minutes at 37°C in a humidified chamber. Next, the cells were incubated overnight at 4°C with a primary monoclonal rat anti-HA antibody (1:500; Roche, #11867423001) diluted in 3% BSA in PBS. Cells were then extensively washed with PBS and incubated for 45 minutes at 37°C in a humidified chamber with the secondary anti-rat IgG Alexa 594 antibody (1:500; Jackson ImmunoResearch, #112-585-167) diluted in 3% BSA in PBS. After that, cells were washed with PBS and permeabilized with ice-cold 0.25% Triton X-100 in PBS for 5 minutes at 4°C, washed again with PBS, and blocked 10% BSA in PBS, as already described. To label type I TARPs, the intracellular FLAG-tag inserted on the C-terminal tail was immunostained using a primary monoclonal mouse Anti-FLAG antibody (1:500; Sigma-Aldrich, #F3165) and a secondary anti-mouse IgG2a Alexa 647 antibody (1:500; Invitrogen Thermo Fisher Scientific, #A-21241). Nuclei were visualized by staining with 1 µg/mL Hoechst 33342 (Sigma-Aldrich, #B2261) in PBS for 5 minutes at room temperature. Lastly, after extensive washing with PBS, coverslips were mounted with Dako fluorescent mounting medium (Agilent Technologies, #S3023). 16-bit z-stack images were acquired on a Zeiss LSM 710 laser scanning confocal microscope (Carl Zeiss) equipped with a Plan-Apochromat 63X oil-immersion objective (NA 1.4) using Zeiss ZEN software (Carl Zeiss).

#### TARP *γ*-2 and Kv7.2 staining in brain slices

10-12-week-old mice were deeply anesthetized with isoflurane and transcardially perfused with ice-cold PBS, followed by freshly prepared 4% PFA in PBS at a constant flow rate of 7 mL per minute using a peristaltic pump (Gilson). Brains were removed and kept in 4% PFA in PBS overnight at 4°C and then transferred to a 30% sucrose in PBS solution and stored at 4°C until they sank (24-48 hours). Brains were sliced using a Cryostar NX50 cryostat (Thermo Fisher Scientific) to obtain 50 µm coronal and sagittal slices. Free-floating slices were washed three times in PBS for 10 minutes and then permeabilized for 1 hour in 0.25% Triton X-100 and 5% horse serum (GIBCO Thermo Fisher Scientific, #16050122) in PBS at room temperature on an orbital shaker for continuous gentle agitation. After that, slices were incubated overnight at room temperature with continuous gentle agitation with a primary polyclonal rabbit anti-Kv7.2 antibody (1:500; abcam, #ab22897) or a polyclonal rabbit anti-TARP γ-2 antibody (1:200; Nittobo Medical, #MSFR105740) diluted in 0.25% Triton X-100 and 2% horse serum in PBS. After extensive washing with 0.25% Triton X-100 in PBS, slices were incubated with the secondary anti-rabbit IgG Alexa 568 antibody (1:500; Invitrogen Thermo Fisher Scientific, #A10042) diluted in 0.25% Triton X-100 and 2% horse serum in PBS for 2 hours at room temperature with continuous agitation. Brain sections were further incubated with 1 µg/mL Hoechst 33342 (Sigma-Aldrich, #B2261) in PBS for 5 minutes at room temperature to stain cell nuclei. Lastly, after three washes in PBS for 10 minutes with mild continuous agitation, brain slices were mounted in gelatinized glass slides using VECTASHIELD mounting medium (Vector Laboratories, #H-1400-10). Images were captured on an Axio Imager Z2 (Carl Zeiss) equipped with a Plan-Apochromat 20X air objective (NA 0.8) using the Zeiss ZEN software (Carl Zeiss).

### Proximity ligation assay (PLA)

Low-density 15 DIV hippocampal and cortical neurons were fixed for 15 minutes at room temperature in 4% PFA and 4% sucrose in PBS. After washing with PBS, cells were permeabilized for 5 minutes in 0.25% Triton X-100 in PBS at 4°C. Neurons were then blocked for 30 minutes in 10% BSA in PBS in a humidified chamber at 37°C and incubated overnight at 4°C with the appropriate combination of primary antibodies prepared in 3% BSA in PBS. To detect the proximity between Kv7.2 and type I TARPs, a mouse monoclonal anti-Kv7.2 antibody (1:200; abcam, #ab84812) was used in combination with either a rabbit polyclonal anti-TARP γ-2 antibody (1:200; Nittobo Medical, #MSFR105740), a rabbit polyclonal anti-TARP γ-3 antibody (1:100; Synaptic Systems, #254 003) or polyclonal anti-TARP γ-8 antibody (1:200; Nittobo Medical, #MSFR105820). TARP antibodies are knock-out validated^93^. Negative controls in which one of the primary antibodies was omitted from the reaction were also included to assess the specificity of the PLA signal. After washing with PBS, followed by three washes with buffer A (0.01 M Tris, 0.15 M NaCl, 0.05% Tween20)^94^, proximity ligation was performed using the Duolink *in situ* PLA Rabbit PLUS (Sigma-Aldrich, # DUO92002) and Mouse MINUS (Sigma-Aldrich, # DUO92004) probes, according to the manufacturer’s instructions, and the Duolink *in situ* Detection Reagents (Sigma-Aldrich, # DUO92013) were used for ligation, amplification, and label probe binding steps. Briefly, neurons were incubated with PLA probes diluted 1:5 in antibody diluent for 2 hours at 37°C in a humidified chamber, followed by several washes with was buffer A. Afterwards, cells were incubated for 30 minutes at 37°C in a humidified chamber with the ligation reaction prepared according to the manufacturer’s recommendations. Amplification and label probe binding were performed after extensive washing with buffer A. Neurons were incubated with the amplification reaction mixture for 1 hour and 45 minutes at 37°C in a humidified chamber. Amplification was stopped by two washes in buffer B (0.2 M Tris, 0.1 M NaCl, pH 7.5)^94^, followed by two washes in PBS. The cells were postfixed for 10 minutes at room temperature in 4% PFA and 4% sucrose in PBS to improve signal stability, washed with PBS, and further processed for staining with cell and nuclear markers as described above. Coverslips were mounted using Dako fluorescent mounting medium (Agilent Technologies, #S3023). Images were captured using an Axio Observer Z1 microscope (Carl Zeiss) equipped with a Plan-Apochromat 63X oil-immersion objective (NA 1.4), with the Zeiss ZEN software (Carl Zeiss).

### SplitFAST fluorescence complementation assay

Live imaging of transfected HEK 293T cells was performed 24 hours after transfection on a Zeiss LSM 710 laser scanning confocal microscope (Carl Zeiss) equipped with a Plan-Apochromat 63X oil-immersion objective (NA 1.4) and a heated stage for temperature control. Images were captured using Zeiss ZEN software (Carl Zeiss). To assess the interaction between Kv7.2 and TARP γ-2 the cells were imaged in an extracellular pH-buffered solution (119 mM NaCl, 5 mM KCl, 2 mM CaCl_2_, 2 mM MgCl_2_, 30 mM glucose, 10 mM HEPES, pH 7.4) at 37°C. mCherry signal was used to identify transfected cells. Samples were imaged continuously for 4 minutes at a frequency of 0.3 frames per second. The baseline signal was recorded for 20 seconds after which 5 µM HMBR FAST fluorogenic ligand (The Twinkle Factory, #480541-250) was added to the cells^39^.

### dSTORM imaging

Images were acquired using a Leica SR GSD microscope (Leica Microsystems) equipped with a Leica HC PL APO 160X oil-immersion TIRF objective (NA 1.43) and an EMCCD iXon camera (ANDOR Oxford Instruments) with a final pixel size of 100 nm, enabling the detection of single fluorophores. Samples were illuminated in TIRF mode, and images were captured using the Leica LAS software (Leica Microsystems) with an exposure time of 12 ms for up to 100,000 consecutive frames. Imaging was performed at room temperature within a sealed Ludin chamber (Life Imaging Services). An extracellular solution with adjusted pH (20 mM Tris-HCl, 50 mM NaCl, 10% glycerol, 10% glucose, pH7.5), supplemented with oxygen scavenger (0.1% catalase suspension #C100, 1 mg/mL pyranose oxidase #P4234, Sigma-Aldrich) and reducing agents (200 mM MEA-HCl #M6500, 4 mM TCEP #C4706, Sigma-Aldrich) was used. The collective fluorescence signal of Alexa Fluor 647 (Thermo Fisher Scientific, #A-21236) was converted into a dark state using half the potency of the 642 nm laser (500 mW). When the targeted count of individual fluorophores per frame was reached, the power of the 642 nm laser was reduced to 15% of its maximum potency. To maintain an optimal number of stochastically activated molecules per frame, the power of the 642 nm laser was continuously adjusted, up to a maximum of 30%. Likewise, the power of the 405 nm laser (30 mW) was continuously adjusted up to a maximum of 10% of its maximum potency. The particle detection threshold was set to 20 in the acquisition software. For dual-color imaging, the 642 nm and 532 nm lasers were used sequentially. After capturing the signal of Alexa Fluor 647, as described above, the fluorescence signal of Alexa Fluor 532 (Thermo Fisher Scientific, #A-11009) was converted into a dark state using the maximum power of the 532 nm laser (1000 mW) in epifluorescence mode. Once the desired number of single fluorophores per frame was reached, the power of the 532 nm laser was reduced to 15% of its maximum potency. To maintain an optimal number of stochastically activated molecules per frame, the power of the 532 nm laser was continuously adjusted, up to a maximum of 30%. Also, the power of the 405 nm laser (30 mW) was continuously adjusted up to a maximum of 10% of its maximum potency. The particle detection threshold was set to 20-50 in the acquisition software.

### Electrophysiological recordings

#### Patch-clamp recordings in CHO cells

Whole-cell patch-clamp recordings on CHO cells were performed 24 hours after transfection, as previously described^62^. The recording chamber was mounted on a Leica DMI3000B inverted microscope (Leica Microsystems) with a mobile stage and perfused with extracellular solution (138 mM NaCl, 2 mM CaCl_2_, 5.4 mM KCl, 1 mM MgCl_2_, 10 mM glucose, 10 mM HEPES, pH 7.4 adjusted with NaOH). Transfected cells were identified by EGFP expression using fluorescence illumination (Leica EL6000 Fluorescence Light Source, Leica Microsystems, #11504115), while transmitted light was used to visualize the selected cells during patch-clamp recordings. Patch pipette electrodes (3-5 mΩ), made from borosilicate glass (King Precision Glass, #8250), were prepared using a vertical microelectrode puller (Narishige PC-10 puller, Biomedical Instruments) and filled with a KCl-based intracellular solution (140 mM KCl, 2 mM MgCl_2_, 10 mM EGTA, 10 mM HEPES, 5 mM Mg-ATP, pH 7.4 adjusted with KOH). Voltage-clamp recordings were performed using an Axopatch 200B patch-clamp amplifier (Axon Instruments, Molecular Devices). Data were acquired at 0.5-2 kHz and filtered at 1-5 kHz with the 4-pole low-pass Bessel filter of the amplifier, using pCLAMP v10.2 software (Axon Instruments, Molecular Devices). No corrections were made for liquid junction potentials. Cells were excluded if the leak current exceeded 100 pA. Linear cell capacitance was determined by integrating the area under the whole-cell capacity transient, evoked by short (5-10 ms) pulses from −80 to −75 mV, with the whole-cell capacitance compensation circuit of the Axopatch 200B turned off. Current densities (expressed in pA/pF) were calculated as peak potassium currents divided by cell capacitance. To generate conductance/voltage (G/V) curves, cells were held at −80 mV and then depolarized for 1.5 seconds from −80 to +40 mV in 20 mV increments, followed by an isopotential pulse at 0 mV for 0.8 seconds. The current values recorded at the beginning of the 0 mV pulse were normalized and expressed as a function of the preceding voltages. The data were fit to a Boltzmann equation, y = max/[1+exp(V_1/2_ -V)/k], where *V* is the test potential, *V_1/2_* is the half-activation potential, and *k* is the slope factor. Currents were corrected offline for linear capacitance and leakage current using the standard subtraction routines of the Clampfit module of pClamp v10.2.

#### Patch-clamp recordings in rat cortical neurons

Whole-cell patch-clamp recordings from 15 DIV transfected cortical neurons plated on poly-D-lysine coated coverslips were performed at room temperature (∼22°C). The recording chamber was mounted on a mobile stage inverted Axio Observer.A1 microscope (Carl Zeiss) and perfused with extracellular solution (140 mM NaCl, 2.4 mM KCl, 10 mM HEPES, 10 mM glucose, 4 mM CaCl_2_, 4 mM MgCl_2_, pH 7.3, 300-310 mOsm), at a constant rate of 1-2 mL per minute. Transfected neurons were identified by EGFP expression using fluorescence illumination (HBO 100 W/2 lamp, OSRAM, #4050300507095), and transmission light differential contrast (DIC) was used to visualize the selected neurons during patch-clamp. Patch pipette electrodes (4–6 mΩ) made from borosilicate glass (Science Products, #GB150F-8P) were pulled with a horizontal microelectrode puller (P-97 Flaming/Brown type micropipette puller, Sutter Instruments), and filled with a K-gluconate-based solution (122 mM K-gluconate, 20 mM KCl, 20 mM HEPES, 13.6 mM NaCl, 3 mM Mg^2+^-ATP, 1 mM EGTA, 0.2 mM CaCl_2_, pH 7.3 adjusted with KOH, 295-300 mOsm). Both voltage- and current-clamp recordings were performed using an EPC 10 USB patch-clamp amplifier (HEKA Elektronik), low-pass filtered at 2.9 kHz using a built-in Bessel filter and digitized at 25 kHz with PATCHMASTER software (HEKA Elektronik). Cells were excluded if the leak current exceeded 200 pA, the series resistance (Rs) was above 25 MΩ or the Rs varied by more than 20%.

For gap-free recordings in current-clamp mode, neurons were clamped at 0 pA. A baseline was recorded for 5 seconds. Subsequently, 30 µM retigabine (Tocris Bioscience, #2687), prepared in extracellular recording solution, was perfused for 5 seconds using a Picospritzer III (Parker Hannifin). This was followed by a 10-second recovery period. The M-current was measured as the hyperpolarizing peak amplitude relative to the membrane resting potential induced by retigabine, an M-channel activator^95^.

For recording M-current in voltage-clamp mode, the extracellular solution was supplemented with 0.5 µM (Tocris Biosciences, #1069), 100 µM CdCl_2_ (Sigma-Aldrich, #202908), and 1 mM 4-AP (Tocris Biosciences, #0940). Neurons were clamped at −20 mV for 200 ms, followed by a series of 1-second-long repolarization steps from −30 mV to –80 mV, after which neurons were held at −20 mV for 500 ms. To pharmacologically isolate the M-current, the protocol was repeated while the cell was locally perfused with 10 µM of the specific M-channel blocker XE991, prepared in extracellular recording solution (Tocris Biosciences, #2000). The M-current was determined by subtracting the XE991 sensitive component from the total deactivating tail currents measured for the voltage step from −20 mV to −50 mV^37^.

#### Subicular pyramidal neuron recordings in hippocampal acute brain slices

8-week-old WT and stagazin V143L heterozygous male mice were deeply anesthetized with isoflurane and transcardially perfused with an ice-cold, oxygenated (95% O_2_ and 5% CO_2_) sucrose solution (217.7 mM sucrose, 2.6 mM KCl, 1.23 mM NaH_2_PO_4_, 26 mM NaHCO_3_, 10 mM glucose, 3 mM MgCl_2_, 1 mM CaCl_2_, pH 7.4, 300-320 mOsm). The brains were quickly removed and placed in ice-cold, oxygenated sucrose solution, and 300 µm hippocampal sagittal slices were prepared using a Leica VT1200s vibrating microtome (Leica Microsystems). The brain slices were transferred to a holding chamber with oxygenated artificial cerebrospinal fluid (aCSF; 125.1 mM NaCl, 2.5 mM KCl, 1.1 mM NaH_2_PO_4_, 25 mM NaHCO_3_, 25 mM glucose, 2 mM MgSO_4_, 2 mM CaCl_2_, pH 7.4, 300-310 mOsm), and kept at 32°C for 30 minutes, followed by 1 hour at room temperature to recover before recording.

Subicular pyramidal neurons were visualized under infrared-differential interference contrast (IR-DIC) microscopy using an upright Axio Examiner.D1 microscope (Carl Zeiss). Whole-cell patch-clamp recordings were performed using a Multiclamp 700B amplifier (Axon Instruments, Molecular Devices), low-pass filtered at 2 kHz, digitized at 20 kHz with Digidata 1550A (Axon Instruments, Molecular Devices), and acquired with the pCLAMP v10.7 software (Axon Instruments, Molecular Devices). Patch pipette electrodes (4–6 mΩ) made from borosilicate glass (Science Products, #GB150F-8P) were pulled using a horizontal microelectrode puller (P-97 Flaming/Brown type micropipette puller, Sutter Instruments). The electrodes were filled with a K-gluconate-based solution (122 mM K-gluconate, 20 mM KCl, 20 mM HEPES, 13.6 mM NaCl, 3 mM Mg^2+^-ATP, 1 mM EGTA, 0.2 mM CaCl_2_, pH 7.3 adjusted with KOH, 295-300 mOsm). Slices were kept at 30°C in a recording chamber perfused with oxygenated aCSF (2-3 mL per minute), supplemented with 100 µM PTX (Tocris Bioscience, #1128), 50 µM D-AP5 (Tocris Bioscience, #0106), and 10 µM CNQX (Tocris Bioscience, #0190) to block synaptic activity. Recordings were performed in the current-clamp mode at a holding current of 0 pA, and the resting membrane potential (RMP) was recorded. Next, a depolarizing current ramp injection from 0-200 pA was applied. The rheobase was determined as the minimum injected current required for the cell to fire an action potential (AP), and the threshold voltage was defined as the membrane potential at which the voltage deflection exceeded 20 mV/ms. Afterwards, the membrane input resistance and the membrane time constant (tau) were estimated from the response of the cell to a rectangular current pulse of −200 pA applied for 100 ms. The steady-state hyperpolarization was measured, and the membrane input resistance was calculated using Ohm’s law. Capacitance (pF) was calculated as (τ / input resistance) x 1000. Each trace was also fitted with a monoexponential decay curve, between 10% of the maximum voltage deflection and the minimum membrane potential during the response, and the membrane time constant was obtained from the fits. Lastly, spike trains were elicited by injecting rectangular current pulses from 0-550 pA in 50 pA incrementing steps, with a duration of 120 ms, followed by 4 seconds recovery at 0 pA. This protocol was repeated while the cell was locally perfused with 30 µM of the M-channel activator retigabine, prepared in extracellular recording solution (Tocris Biosciences, #2000), using a Picospritzer III (Parker Hannifin). The medium after burst-hyperpolarization (mAHP) was measured in the current injection step eliciting a 50-60 Hz firing frequency by measuring the peak voltage in the 20-100 ms post-burst interval^96^. Holding current and Rs were monitored, and cells were discarded if either of these parameters varied more than 20% or if Rs was above 25 MΩ.

### Seizure induction in stargazin V143L mice

#### Pentylenetetrazol (PTZ)-induced seizures

10-12-week-old male WT and stargazin V143L littermate mice were injected intraperitoneally with a single dose of 40 mg/kg PTZ (Tocris Bioscience, #2687), a GABAA receptor antagonist. The PTZ dose used was selected based on a pilot test. Mice were placed in an open field arena with transparent walls and continuous video monitoring immediately after the injection. The severity of the seizures was assessed by observing the behavior of the animals, and the latency to reach phase 5/6 according to the PTZ-adapted Racine scale^53^ was determined by a blinded observer. In this scale, phase 5 is defined as bilateral clonic and tonic-clonic seizures (lying on the belly), whereas phase 6 is characterized by both bilateral clonic and tonic-clonic seizures, as well as loss of righting reflex (lying on the side). The experiment was conducted in a room at 28-30°C with controlled humidity. Animals were acclimatized to the room for 2 hours before the experiment, with *ad libitum* access to food and water.

#### XE991-induced seizures

To provoke seizures by suppressing M-currents, 10-12-week-old male WT and stargazin V143L littermate mice were intraperitoneally injected with a single dose of 15 mg/kg XE911 (Tocris Bioscience, #2000), an M-channel blocker, under the same experimental conditions described for PTZ-induced seizures. The XE991 dose used was selected based on a pilot test. Since mice injected with XE991 exhibited behavioral manifestations similar to those described in phases 1-3 of the classical Racine scale^97^, the videos were analyzed by a blinded observer for scoring using a combination of the classical and the PTZ-modified Racine scales.

## Quantification and statistical analysis

### Protein identification and gene ontology analysis

The mass spectra were imported into MaxQuant software (version 1.5) and cross-referenced with the Uniprot mouse proteomic database (version 2013-01-06). The mass tolerances were set to 6 ppm and 0.5 Da for MS and MS/MS, respectively. Up to two missense cleavages were allowed, with trypsin as the selected protease. Cysteine alkylation with acrylamide was set as a fixed modification, while methionine oxidation and N-terminal acetylation were set as variable modifications. The false discovery rate (FDR) was set at 0.01. Only unique peptides were used for identification and protein hits were considered valid if they contained at least one unique peptide. Valid hits were considered putative interactors of TARP-γ2 if they co-immunoprecipitated with TARP-γ2 and were absent in control samples (immunoprecipitation performed with non-immune IgGs) in at least one of the duplicates from two independent experiments. To identify ontology groups for biological processes associated with the list of TARP-γ2 interacting proteins, gene ontology (GO) analysis was performed using PANTHER GO-slim (version 19.0, released 2024-06-19; https://www.pantherdb.org/panther/goSlim.jsp)^82^.

### Image analysis

#### PLA puncta analysis

8-10 fields on each coverslip were randomly selected, and stargzin/Kv7.2 PLA puncta were quantified using the Surfaces model in Imaris v10.1.0 software (Bitplane, Oxford Instruments). The number of PLA puncta was normalized for the area of the neurons visible in each image. The area was defined using phalloidin staining and quantified with ImageJ/Fiji software (National Institutes of Health). Data are presented as percentage of the respective control.

#### Analysis of axon initial distance from the soma

The distance from the soma to the axon initial segment (AIS) in cultured cortical neurons was measured manually using ImageJ/Fiji software (National Institutes of Health). Only neurons with a single axon originating from the soma were included in the analysis. The EGFP signal of transfected cells was used to identify the soma, while Ankyrin G labeling was used to visualize the AIS. All analyses were performed blind to the experimental conditions.

#### AMPA receptor surface and synaptic levels

All experimental conditions were prepared and stained simultaneously, with acquisition settings maintained across conditions within each independent experiment. GluA-AMPA receptor surface and synaptic puncta number, area, and intensity were analyzed using ImageJ/Fiji software (National Institutes of Health) with an in-house macro that automated the quantification steps. 11-12 fields on each coverslip were randomly selected, and dendritic regions of interest (ROIs) were defined based on the MAP2 staining. The dendritic length was measured, and background and intensity threshold settings were adjusted for each channel. The threshold was defined to include all visible puncta. The average background value for each image was subtracted from the thresholded mean intensity value to obtain the corrected intensity value. These corrected intensity values were then multiplied by the puncta area to determine the final integrated intensity. Synaptic GluA-AMPA receptor puncta were defined by their co-localization with PSD95. For this, binary masks were created for each channel and overlaid with the other to test for the co-localization of GluA-AMPA receptor puncta with PSD95 puncta.

#### Cortical neurons viability

Cortical neurons viability was assessed by analyzing nuclear morphology using Hoechst 33342 staining. 10-12 images were randomly captured per coverslip. The total number of cells per sample was manually counted using ImageJ/Fiji software (National Institutes of Health), and the percentage of cells with condensed and fragmented nuclei was determined.

#### SplitFAST fluorescence analysis

SplitFAST signal quantification was performed using ImageJ/Fiji software (National Institutes of Health). First, z-series images were corrected for small xy-drift using the TurboReg plugin^98^. Next, the mCherry signal was used to manually annotate regions of interest (ROIs) corresponding to individual cells. The integrated intensity of the mCherry and HMBR fluorescence signals was analyzed over time. For each ROI, the baseline signal for each channel was obtained by averaging the integrated intensity of the frames acquired during the first 20 seconds before the addition of the fluorogen HMBR. HMBR and mCherry signals were expressed as a percentage of the respective baseline. For the HMBR channel, since the baseline fluorescent signal was background, this was subtracted from all data points.

#### dSTORM image analysis

Super-resolution images were reconstructed with the Leica LAS software (Leica Microsystems) using the center of mass fitting algorithm. This algorithm determined the centroid-coordinates of individual molecules and fitted the point-spread-function (PSF) of each distinct diffraction-limited event to a Gaussian function. The resulting super-resolved images had a final spatial resolution of 20 nm. To correct lateral drift, multicolor fluorescent microspheres were employed (Invitrogen Thermo Fisher Scientific, #T7279). The SR-Tesseler software^84^ was then used to analyze protein clusters. Single molecule spatial coordinates were used to compute a Voronoï diagram, decomposing the image into polygons of different sizes, each centered on a localized molecule. First-rank densities (δ *^1^*) were calculated, and density maps were produced by applying the δ *^1^* values as textures to the Voronoï polygons. The Kv7.2 clusters were automatically segmented by applying a threshold of two times the average first-rank density of the entire dataset, with a minimum area of 0.04 pixel^2^ and requiring at least 50 localizations. The minimum area was defined by considering the size of a single Kv7.2 channel, which is 20 nm^48^. The number of detections was determined after a pilot experiment, which compared specific Kv7.2 clusters with non-specific clusters by omitting the primary antibody in the staining protocol. In the case of stargazin, clusters were detected by applying a threshold of two times the average density δ of the entire dataset, with a minimum area of 1 pixel^2^, based on the minimum size of stargazin clusters in low-resolution images. A threshold of one time the average density of each cluster, with a minimum area of 0.04 pixel^2^ and at least 50 localizations was used. Size parameters and local density (number of detections per cluster area) were extracted after cluster segmentation by principal component analysis.

For co-localization analyses, each fluorescence channel was segmented independently as described, and co-localization was computed using the PoCA software^85^. The total Kv7.2 expression levels were quantified from epifluorescence images acquired from the same samples under control conditions and when stargazin was overexpressed, using ImageJ/Fiji software (National Institutes of Health).

#### Total and surface staining of Kv7.2 in HEK 293T cells HEK 293T

In each independent experiment, 14-16 cells were randomly selected. FLAG-tag staining was used to confirm the expression of TARPs in the selected cells when co-expressed with Kv7.2. All experimental conditions were prepared and stained simultaneously, with acquisition settings maintained across conditions within each independent experiment. Surfaces were generated from z-stack images in imaris v10.1.0 software (Bitplane, Oxford Instruments) using the combined fluorescence signal from the total and surface Kv7.2 stainings. The intensity sum of the fluorescent signal for both the total (EGFP fluorescence) and surface (HA-tag immunostaining) Kv7.2 stainings for each cell was obtained from these volumetric reconstructions. For each independent experiment, the data were normalized to the average intensity sum of the respective control.

### Electrophysiology data analysis

All recordings were analyzed using pCLAMP v10.7 (Axon Instruments, Molecular Devices). Data acquired with the PATCHMASTER software (HEKA Elektronik) were converted to ABF file format before analyses. All electrophysiology data analyses were done blindly, without knowledge of the experimental conditions.

### Statistics

For each experiment, the normality of population distributions was assessed using the Shapiro-Wilk normality test. Based on this evaluation, either parametric or non-parametric tests were employed, as described in the figure legends. Accordingly, data are presented as mean ± SEM or in box and whiskers plots as median ± interquartile range, as indicated in the figure legends. Over time measurements are presented as mean ± SD. The number of independent experiments is indicated in the figure legends. For all tests, statistical significance was considered as *p <* 0.05. Variance analysis following one-way ANOVA was performed using the Brown-Forsythe test. Before running a *t*-test, variance analysis was performed using the F-test. When the variances of the two populations differed, a Welch’s *t*-test was applied. All statistical analyses were performed using GraphPad Prism 10.1.2 software (GraphPad Software). Additional information regarding the number of independent experiments, the statistical tests employed, and the corresponding *p*-values is available in Table S3.

The schemes in Figure 1E, 5A, and S1F were created with BioRender (BioRender.com).

## Acknowledgments

The pcDNA3.1_Kv7.2 and pcDNA3.1_Kv7.3 plasmids were a kind gift from Prof. Thomas J. Jentsch (Zentrum für Molekulare Neurobiologie Hamburg, Germany); the pcDNA3.1_TARPγ2 plasmid was a kind gift of Dr. Verity Letts (The Jackson Laboratory, USA); and the piRES2-EGFP_TARPγ3, piRES2-EGFP_TARPγ4, and piRES2-EGFP_TARPγ8 plasmids were a kind gift from Prof. Mark Farrant (University College London, UK). This work was funded by La Caixa Iniciativa Ibérica and FCT 4b initiative under the project LCF/PR/HP20/52300003, and by the European Regional Development Fund (ERDF), through the COMPETE 2020 — Operational Program for Competitiveness and Internationalization and Portuguese national funds via FCT, under projects UIDB/04539/2020, UIDP/04539/2020, LA/P/0058/2020. MVR was funded by FCT through the following fellowships: SFRH/BD/129236/2017 and CRM:0026298. The authors wish to thank Francesco Micelli for insightful discussions, and the MICC Imaging facility of CNC-UC.

## Notes

### Competing Interest Statement

The authors have declared no competing interest.

