## Supplemental information for "Type I TARPs regulate Kv7.2 potassium channels and susceptibility to seizures"

#### Supplemental Figures:

FIGURE S1

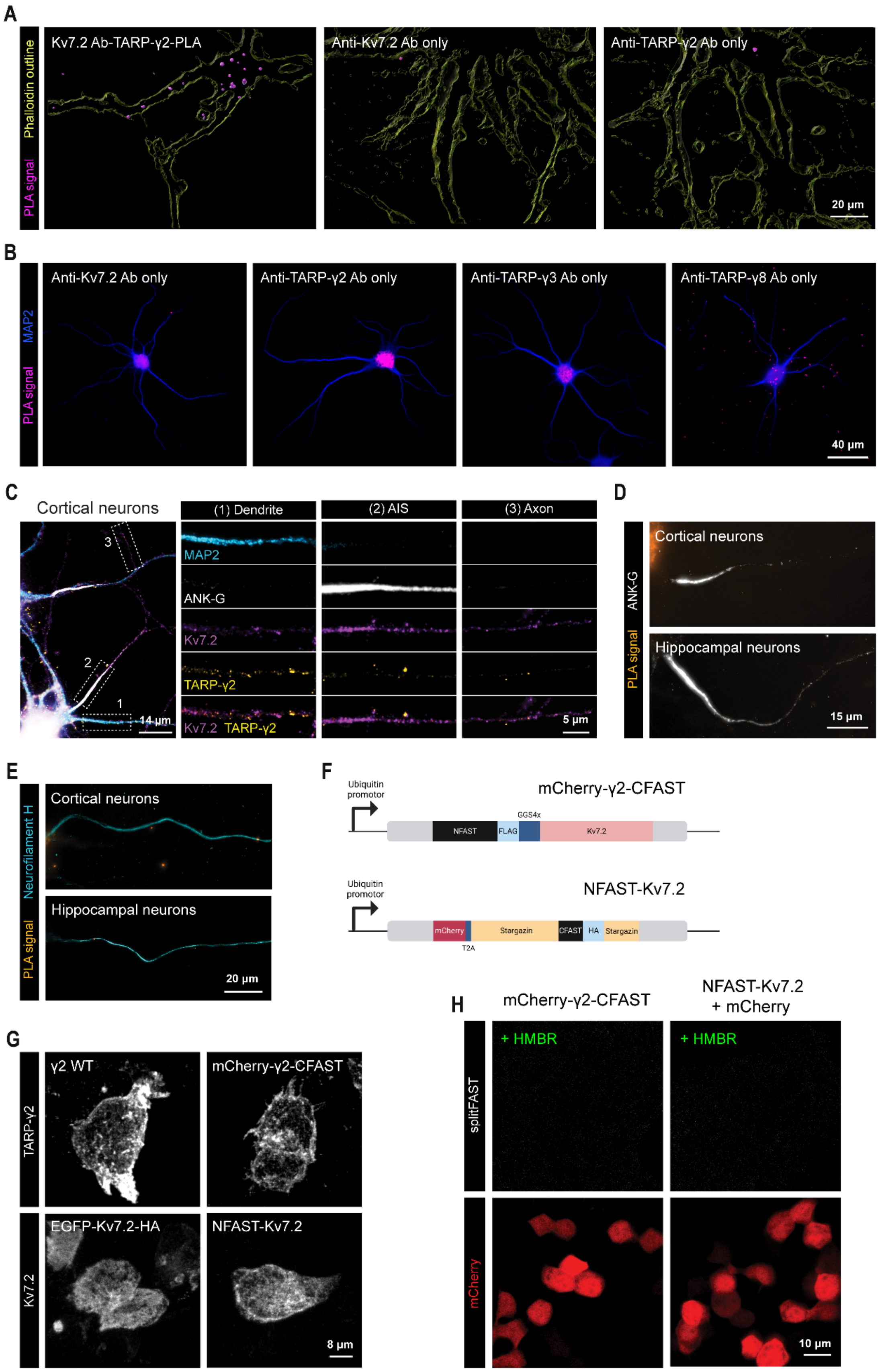
A
Phalloidin outline
PLA signal
Kv7.2 Ab-TARP-γ2-PLA
Anti-Kv7.2 Ab only
Anti-TARP-γ2 Ab only
20 µm
B
MAP2
PLA signal
Anti-Kv7.2 Ab only
Anti-TARP-γ2 Ab only
Anti-TARP-γ3 Ab only
Anti-TARP-γ8 Ab only
40 µm
C
Cortical neurons
(1) Dendrite
(2) AIS
(3) Axon
MAP2
ANK-G
Kv7.2
TARP-γ2
Kv7.2 TARP-γ2
14 µm
5 µm
D
ANK-G
PLA signal
Cortical neurons
Hippocampal neurons
15 µm
E
Neurofilament H
PLA signal
Cortical neurons
Hippocampal neurons
20 µm
F
Ubiquitin promotor
mCherry-γ2-CFAST
NFAST
FLAG
GGS4x
Kv7.2
NFAST-Kv7.2
mCherry
T2A
Stargazin
CFAST
HA
Stargazin
G
TARP-γ2
Kv7.2
γ2 WT
mCherry-γ2-CFAST
EGFP-Kv7.2-HA
NFAST-Kv7.2
8 µm
H
mCherry-γ2-CFAST
NFAST-Kv7.2 + mCherry
splitFAST
mCherry
+ HMBR
+ HMBR
10 µm

**Figure S1. Type I TARPs interact with the Kv7.2 subunit of the M-channels and localize to the same subcellular compartments, related to Figure 1.**

A) Representative image of proximity ligation assay (PLA) signal for TARP-γ2 and Kv7.2 (pink) in 15 DIV rat cortical neurons. Negative controls, where one of the primary antibodies was omitted from the PLA reaction, are also shown. The contour of the neurons was defined using phalloidin staining (yellow). B) Representative images of PLA signal (pink) negative controls in 15 DIV rat hippocampal neurons, where the specific primary antibody for each type I TARP tested or Kv7.2 was omitted from the reaction. Neurons were also labeled for MAP2 (blue). C) Subcellular distribution of Kv7.2 and TARP-γ2 in 15 DIV rat cortical neurons. Immunolabeling of both Kv7.2 (pink) and TARP-γ2 (yellow) is observed in the dendritic compartment (1), identified by MAP2 staining (cyan), in the axon initial segment (AIS; 2), delimited by ankyrin G staining (white), and in axons (3). D) No PLA signal (orange) was detected for TARP-γ2 and Kv7.2 in the AIS (ankyrin G, white) or in E) axons (neurofilament H, cyan) of rat cortical and hippocampal neurons. F) Schematic representation of the TARP-γ2 and Kv7.2 splitFast constructs. G) Representative fluorescence images showing the expression patterns of wild-type TARP-γ2 and Kv7.2, as well as the splitFast constructs of TARP-γ2 and Kv7.2, in transfected HEK293T cells. The splitFAST constructs show a similar expression pattern to that of the wild-type proteins. H) Representative live-cell confocal microscopy images of HEK239T cells transfected with either mCherry-γ2-CFAST or NFAST-Kv7.2 and mCherry, acquired 200 seconds after the addition of the HMBR fluorogen. The splitFast fluorescence signal is not detected when CFAST or NFAST are not co-expressed in the same cell.

FIGURE S2

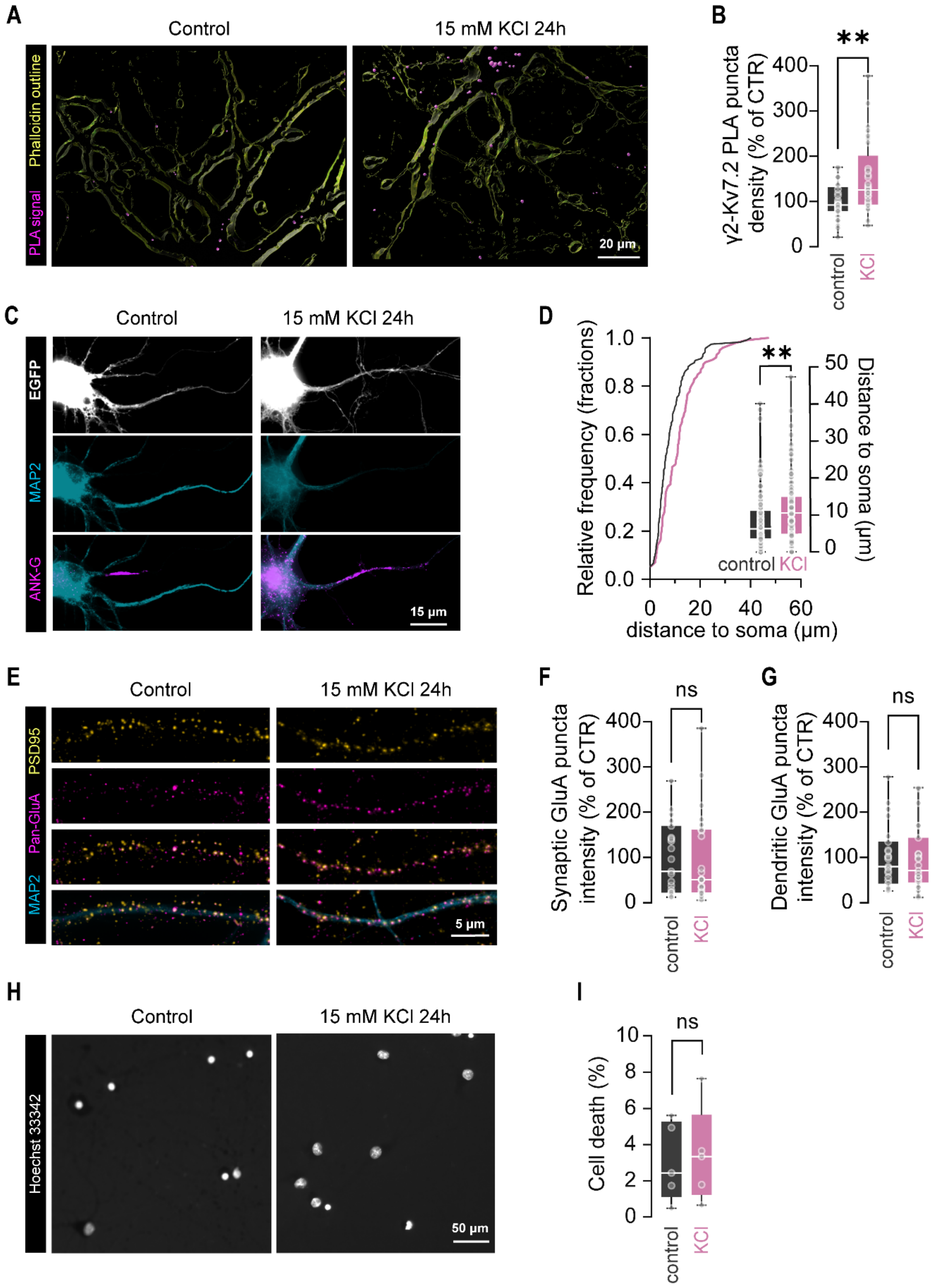


59
60

**Figure S2. Interaction between TARP-γ2 and Kv7.2 is potentiated by neuronal depolarization, related to Figure 1.**

A) Representative images of PLA signal (pink) for endogenous TARP-γ2 and Kv7.2 in 15 DIV rat cortical neurons after incubation with 15 mM KCl for 24 hours. The contour of the neurons was defined using phalloidin staining (yellow). B) Increasing the extracellular KCl concentration from 5 mM to 15 mM for 24 hours significantly enhanced the interaction between TARP-γ2 and Kv7.2 in rat cortical neurons. Data are presented in whisker plots as median ± interquartile range (IQR), with boxes showing $25^{th}$ and $75^{th}$ percentiles, whiskers ranging from the minimum to the maximum values, and the horizontal lines representing the median. Two-tailed Mann-Whitney test, $^{**}p=0.0072$. N=3 independent experiments, n=29 for control, n=29 neurons for KCl. C) Representative fluorescence images showing the position of the axon initial segment (AIS) in 15 DIV rat cortical neurons transfected with an EGFP plasmid, under control conditions and after exposure to 15 mM KCl for 24 hours. The AIS was visualized by ankyrin G staining (pink), the EGFP signal (white) was used to identify the soma, and dendrites were labelled with MAP2 (cyan). D) Cumulative distribution and whisker plots of the AIS distance from the soma under control and depolarizing conditions for 24 hours, showing a distal shift of the AIS from the soma in KCl-treated neurons compared to control neurons. Data presented in whisker plots are shown as median ± IQR. Two-tailed Mann-Whitney test, $^{**}p = 0.0016$. N=11 independent experiments, n=117 neurons for control, n=99 neurons for KCl. E) Representative images of total and synaptic levels of AMPAR (pink) under control conditions and after depolarization with 15 mM KCl for 24 hours. Synaptic sites were identified by immunolabeling PSD95 (yellow), and dendrites were visualized with MAP2 staining (cyan). F) The depolarizing KCl stimulus did not alter the number of synaptic AMPAR that colocalized with PSD95. Two-tailed Mann-Whitney test, ns $p=0.7940$. G) Total levels of AMPAR remained unchanged after 24 hours of incubation with 15 mM of KCl. Two-tailed Mann-Whitney test, ns $p=0.8109$. F) and G) Results are presented in whisker plots as median ± IQR. N=2 independent experiments, n=23 neurons for control, n=23 neurons for KCl. H) Representative images of Hoechst 33342 nuclear staining (white) used to assess neuronal cell viability after 24 hours of treatment with 15 mM KCl. I) Depolarization with KCl for 24 hours did not increase neuronal cell death. Data are presented in whisker plots as median ± IQR. Two-tailed unpaired *t*-test, ns $p=0.0836$. N=5 independent experiments, n ≥ 53 images per experimental condition.

**FIGURE S3**

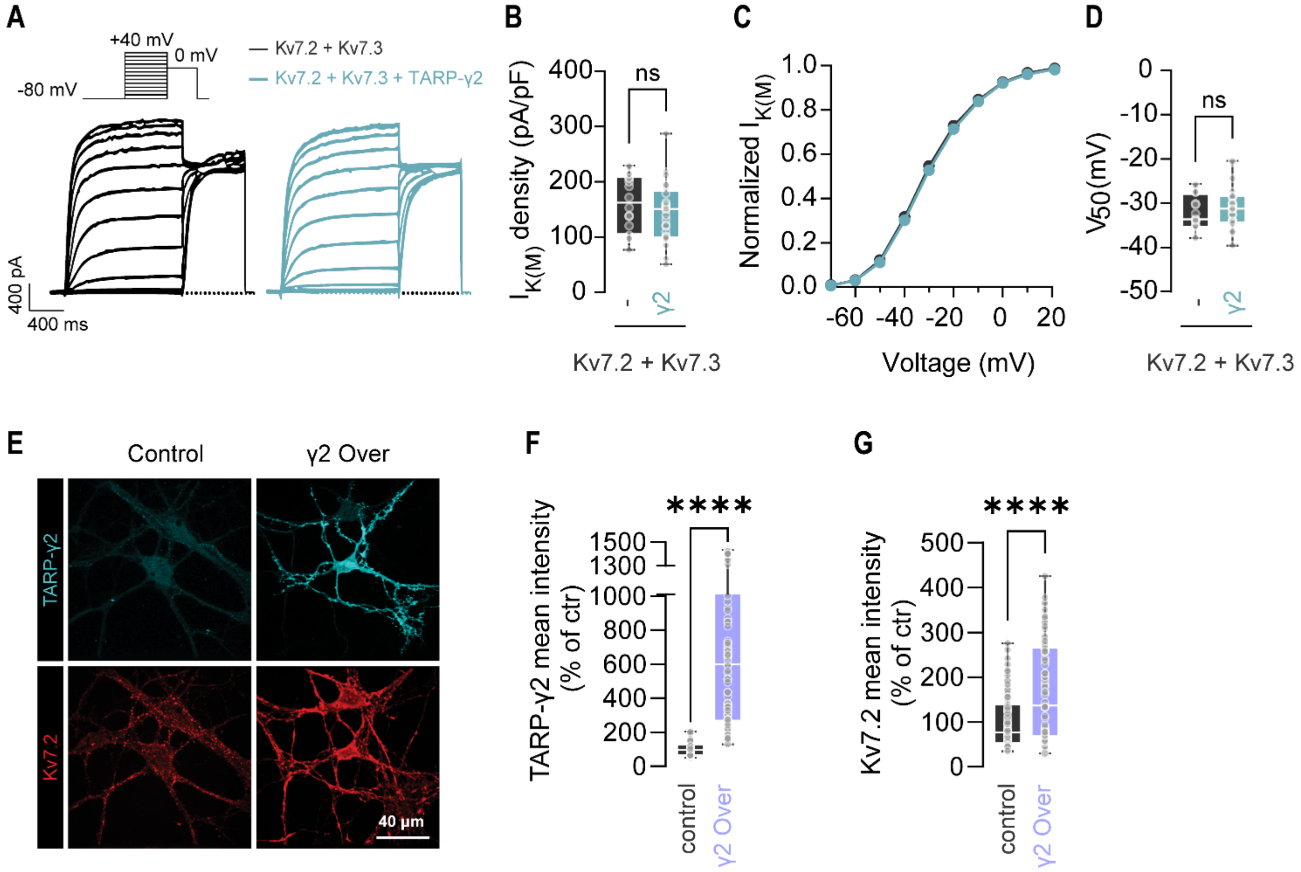


**Figure S3. TARP-γ2 regulates homomeric Kv7.2-M-currents but not heteromeric Kv7.2/Kv7.3-M-currents and increases the neuronal expression of Kv7.2, related to Figure 2.** A) Representative traces of heteromeric Kv7.2/7.3-M–currents measured in CHO cells transfected with Kv7.2 and Kv7.3 or co-transfected with Kv7.2, Kv7.3, and TARP-γ2. To generate conductance–voltage curves, cells were held at –80 mV and depolarized for 1.5 seconds from –80 to +40 mV in 20 mV increments, followed by an 800 ms isopotential pulse at 0 mV. B) TARP-γ2 co-expression did not potentiate the currents mediated by Kv7.2/7.3 heteromeric M-channels. Results are presented in whisker plots as median ± IQR (boxes show the 25$^{th}$ and 75$^{th}$ percentiles, whiskers range from the minimum to the maximum values, and the horizontal lines represent the median). Two-tailed unpaired *t*-test, ns $p$=0.3643. C) Current/voltage curves showing the impact of TARP-γ2 on heteromeric Kv7.2/7.3 voltage-gating properties. Data are presented as mean ± SEM. D) The half-activation potentials ($V_{50}$) were determined by fitting the current/voltage curves to a Boltzmann distribution. TARP-γ2 did not regulate the gating properties of heteromeric Kv7.2/Kv7.3-M-channels. Data are presented in whisker plots as median ± IQR. Two-tailed unpaired *t*-test, ns $p$=0.4240. B-D) N ≥ 3 independent experiments, n=16 cells for Kv7.2 + Kv7.3, n=25 cells for Kv7.2 + Kv7.3 + TARP-γ2. E) Representative fluorescence images showing endogenous Kv7.2 expression in 15 DIV rat cortical neurons following TARP-γ2 overexpression. Neurons were transfected with either a control EGFP plasmid or a wild-type TARP-γ2 plasmid (γ2 Over) and stained for Kv7.2 (red) and TARP-γ2 (cyan). F) TARP-γ2 was significantly overexpressed in neurons transfected with the TARP-γ2 plasmid compared to control-transfected neurons. Two-tailed Mann-Whitney test, ****$p$ < 0.0001. G) Endogenous Kv7.2 expression was significantly increased by overexpressing TARP-γ2 compared to control-transfected neurons. Two-tailed Mann-Whitney test, ****p < 0.0001. F and G) Data are presented in whisker plots as median ± IQR. N=4 independent experiments, n ≥ 88 neurons for each experimental condition.

**FIGURE S4**

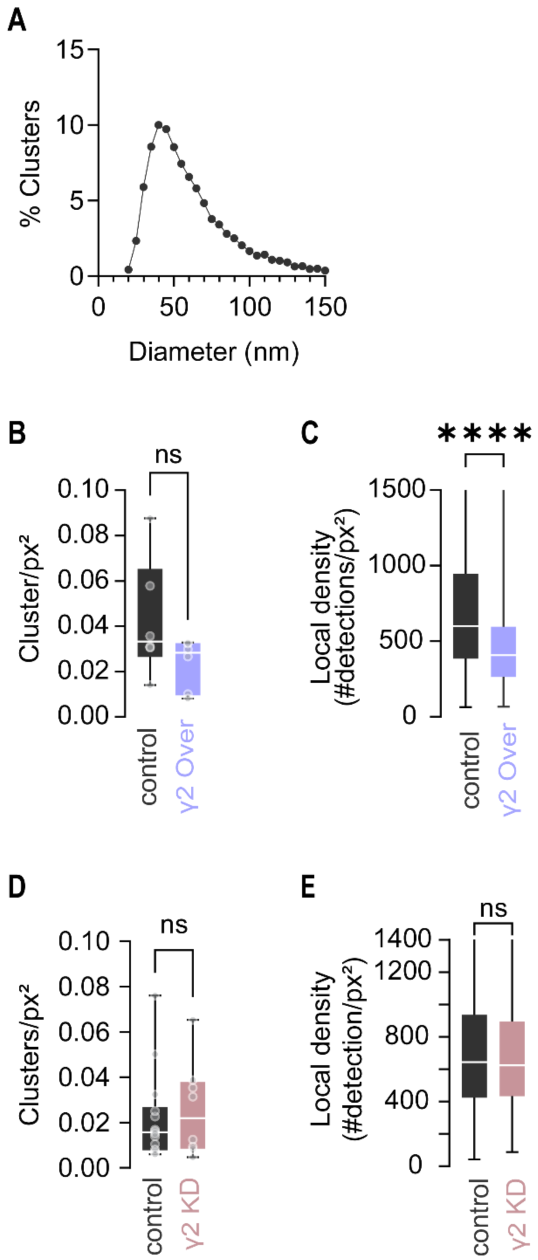

**Figure S4. Genetic manipulation of TARP-γ2 expression alters endogenous Kv7.2 nano-clustering at dendritic sites, related to Figure 3.**

A) Distribution of Kv7.2 cluster diameters in 15 DIV rat cortical neurons. N > 3 independent experiments, n=7387 clusters. B) Overexpressing TARP-γ2 (γ2 Over) did not impact dendritic Kv7.2 cluster density. Two-tailed Mann-Whitney test, $p$=0.1320. N=4 independent experiments, n=6 neurons per experimental condition. C) The local density of Kv7.2 dendritic clusters in cortical neurons was significantly decreased following TARP-γ2 overexpression. Two-tailed Mann-Whitney test, ****$p$ < 0.0001. N=4 independent experiments, n=6 neurons. N=4 experiments, n=787 clusters. D) TARP-γ2 knockdown (γ2 KD) did not change the density of Kv7.2 clusters in the dendrites of cortical neurons. Two-tailed Mann-Whitney test, p=0.7639. N=5 independent experiments, n ≥ 8 neurons for each experimental condition. E) The local density of Kv7.2 dendritic clusters remained unchanged in cortical neurons after TARP-γ2 silencing. Two-tailed Mann-Whitney test, p=0.6839. B-E) Results are presented in whisker plots as median ± IQR (boxes show 25th and 75th percentiles, whiskers range from the minimum to the maximum values, and the horizontal lines represent the median).

**FIGURE S5**

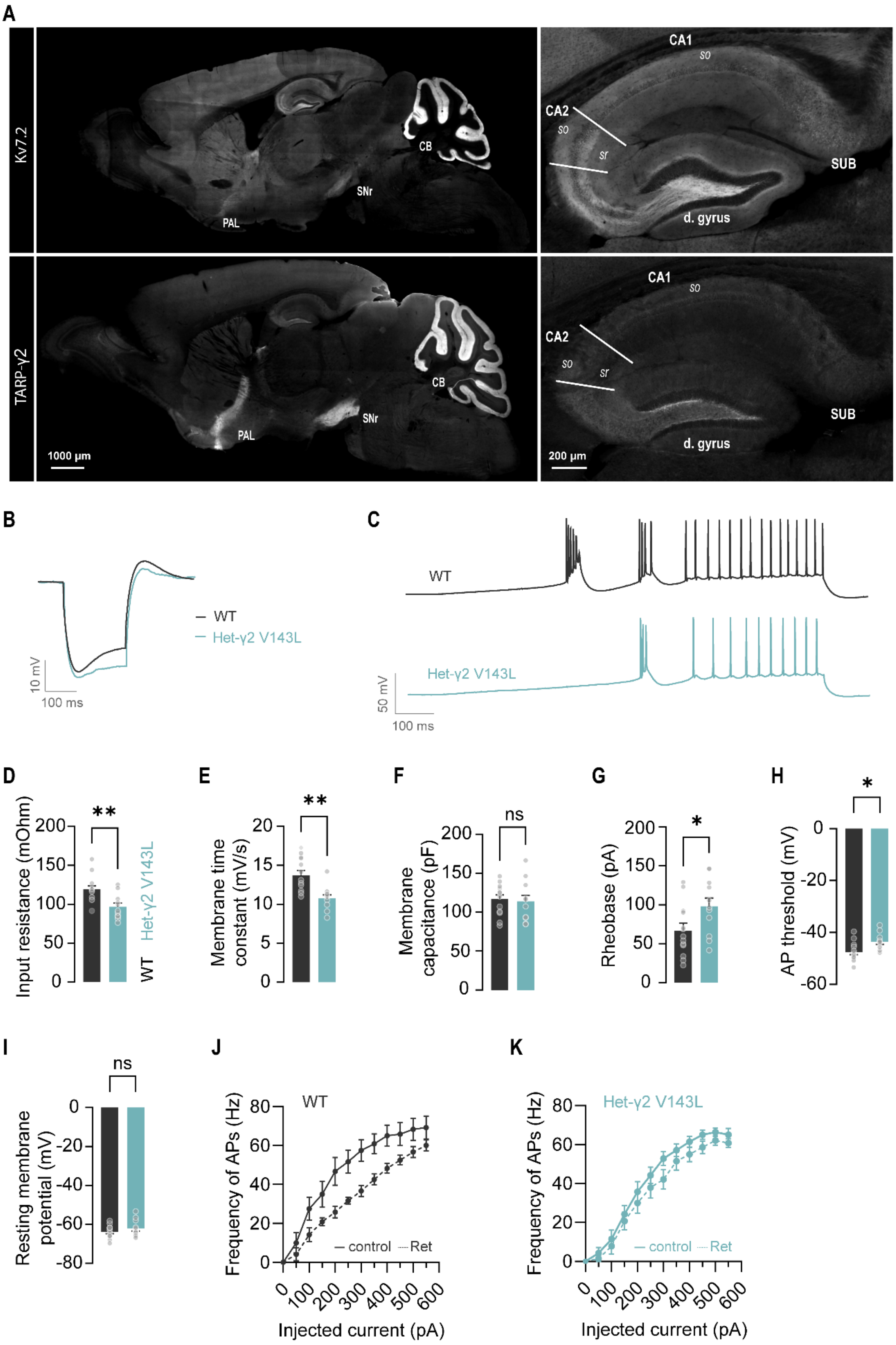
A
Kv7.2
TARP-γ2
CB
SNr
PAL
1000 µm
CA1
CA2
so
sr
SUB
d. gyrus
200 µm
B
WT
Het-γ2 V143L
10 mV
100 ms
C
WT
Het-γ2 V143L
50 mV
100 ms
D
Input resistance (mOhm)
**
WT
Het-γ2 V143L
E
Membrane time constant (mV/s)
**
F
Membrane capacitance (pF)
ns
G
Rheobase (pA)
*
H
AP threshold (mV)
*
I
Resting membrane potential (mV)
ns
J
WT
Frequency of APs (Hz)
control
Ret
Injected current (pA)
K
Het-γ2 V143L
Frequency of APs (Hz)
control
Ret
Injected current (pA)

**Figure S5. The disease-associated TARP-γ2 V143L mutant re-shapes intrinsic neuronal excitability in the subiculum of a knock-in mouse model, related to Figure 5.**

A) Representative fluorescence images showing the expression pattern of Kv7.2 and TARP-γ2 in the mouse brain. The proteins were immunolabeled in sagittal sections of 2-month-old mice, revealing a partial overlap in their expression patterns. TARP-γ2 and Kv7.2 were co-enriched in natural seizure-controlling brain areas, including the cerebellum (CB), pallidum (PAL), substantia nigra pars reticulata (SNr), dentate gyrus (d. gyrus), and subiculum (SUB). (so) stratum oriens; (sr) stratum radiatum. B) Representative traces showing the response of subicular burst-firing pyramidal neurons to a hyperpolarizing -200 pA current injection in wild-type (WT; dark grey) and heterozygous TARP-γ2 V143L mice (Het-γ2 V143L; cyan). Whole-cell patch-clamp recordings were performed in acute slices from 8-week-old mice in the presence of synaptic transmission blockers. Input resistance and membrane time constant (tau) were estimated from these voltage responses. C) Representative traces from the same neurons in acute slices from WT and Het-γ2 V143L mice showing their response to a ramp current injection (0 to +200 pA). The rheobase and the action potential (AP) threshold were determined from these voltage responses. D) The input resistance was significantly reduced in subicular burst-firing pyramidal neurons expressing TARP-γ2 V143L. Two-tailed unpaired *t*-test, $^{**}p = 0.0033$. E) In subicular neurons from Het-γ2 V134L mice, the membrane time constant was significantly decreased. Two-tailed unpaired *t*-test, $^{**}p=0.0011$. F) Membrane capacitance was unchanged in neurons expressing the TARP-γ2 V143L mutant. Two-tailed unpaired *t*-test, ns $p=0.7784$. G) Rheobase was significantly increased in Het-γ2 V134L animals compared to WT mice. Rheobase was defined as the minimal current of infinite duration required for the neuron to initiate an AP. Two-tailed unpaired *t*-test $^{*}p=0.0405$. H) The AP threshold was significantly increased in subicular neurons from Het-γ2 V143L mice. Two-tailed unpaired *t*-test, $^{*}p=0.0129$. I) The resting membrane potential was not significantly changed in subicular pyramidal neurons of Het-γ2 V143L mice compared to WT mice. Two-tailed unpaired t-test, p=0.2618. B-I) Data presented as mean ± SEM. N ≥ 4 animals and n ≥ 11 neurons for each genotype. J) Mean firing frequency of subicular neurons plotted as a function of the injected depolarizing current before (control) and during retigabine perfusion (Ret) in WT and K) Het-γ2 V143L mice. Data presented as mean ± SEM. N ≥ 4 animals and n=7 neurons for each genotype.

### Supplemental tables:

**Table S1. Gene ontology (GO) analysis of the stargazin interactome.** The stargazin interactome was investigated in cultured cortical neurons by immunoprecipitation of stargazin, followed by mass spectrometry. Newly identified putative stargazin interactors were analyzed using PANTHER GO-slim to identify ontology groups for biological processes.

| **PUTATIVE** | **MOLECULAR** | **BIOLOGICAL PROCESS** | **PROTEIN CLASS** |
|---|---|---|---|
| Potassium voltage-gated channel subfamily KQT member 2 (Kcnq2) | voltage-gated potassium channel activity | potassium ion transmembrane transport | voltage-gated ion channel |
| Glycogen phosphorylase, brain form (Pygb) | hexosyltransferase activity<br>heterocyclic compound binding<br>organic cyclic compound binding<br>anion binding | glycogen metabolic process<br>macromolecule catabolic process<br>carbohydrate catabolic process | *glycosyltransferase* |
| Acyl-CoA-binding protein (Dbi) | nucleotide binding<br>carbohydrate derivative binding<br>lipid binding<br>anion binding<br>amide binding | fatty acid metabolic process | transfer/carrier protein |
| Cysteine-tRNA ligase, cytoplasmic (Cars) | ATP binding<br>catalytic activity, acting on RNA<br>ligase activity | amino acid metabolic process<br>tRNA metabolic process<br>translation | aminoacyl-tRNA synthetase |
| Valine-tRNA ligase (Vars) | catalytic activity, acting on RNA<br>ligase activity | amino acid metabolic process<br>tRNA metabolic process<br>translation | aminoacyl-tRNA synthetase |
| Asparagine-tRNA ligase, cytoplasmic (Nars) | catalytic activity, acting on RNA<br>ligase activity | amino acid metabolic process<br>tRNA metabolic process<br>translation | *aminoacyl-tRNA synthetase* |
| Quaking protein (Qki) | mRNA binding | regulation of mRNA splicing, via spliceosome | RNA splicing factor |
| Myelin expression factor 2 (Myef2) | mRNA binding | - | RNA splicing factor |
| Large ribosomal subunit protein eL24 (Rpl24) | structural constituent of ribosome<br>mRNA binding | cytoplasmic translation | ribosomal protein |

| | | | |
|---|---|---|---|
| Small ribosomal subunit protein uS5 (Rps2) | structural constituent of ribosome | translation | ribosomal protein |
| Small ribosomal subunit protein uS3 (Rps3) | structural constituent of ribosome | - | ribosomal protein |
| Large ribosomal subunit protein uL10 (Rplp0) | structural constituent of ribosome<br>rRNA binding | cytoplasmic translation<br>ribosomal large subunit assembly | ribosomal protein |
| Large ribosomal subunit protein eL8 (Rpl7a) | RNA binding | maturation of LSU-rRNA | ribosomal protein |
| ATP-dependent RNA helicase A (Dhx9) | RNA binding<br>helicase activity | - | RNA helicase |
| Elongation factor 1-beta (Eef1b) | guanyl-nucleotide exchange factor activity | translational elongation | translation elongation factor |
| Elongation factor 1-gamma (Eef1g) | - | translational elongation | - |
| T-complex protein 1 subunit alpha (Tcp1) | unfolded protein binding | protein folding | Chaperonin |
| Dynactin subunit 1 (Dctn1) | - | - | Chaperone |
| Stress-70 protein, mitochondrial (Hspa9) | - | - | Hsp70 family chaperone |
| 3-ketoacyl-CoA thiolase, mitochondrial (Acaa2) | - | - | Acyltransferase |
| GTP-binding nuclear protein Ran (Ran) | GTPase activity | ribosomal subunit export from nucleus<br>protein import into nucleus | small GTPase |
| Signal-induced proliferation-associated 1-like protein 1 (Sipa1l1) | GTPase activator activity | - | GTPase-activating protein |
| ELKS/Rab6-interacting/CAST family member-1 (Erc1) | structural molecule activity | neuromuscular synaptic transmission<br>synapse organization<br>regulation of synaptic plasticity | membrane traffic protein |
| Beta-synuclein (Sncb) | copper ion binding | synapse organization<br>synaptic vesicle endocytosis<br>chemical synaptic transmission | membrane trafficking regulatory protein |
| AP-2 complex subunit mu (Ap2m1) | - | endocytosis | membrane traffic protein |
| Ras-related protein Rab-10 (Rab10) | myosin binding | exocytosis<br>protein secretion | - |

| | | | |
|---|---|---|---|
| Membrane-associated phosphatidylinositol transfer protein 2 (Pitpnm2) | lipid transfer activity<br>phosphatidylinositol binding<br>phosphatidylcholine binding<br>phosphatidylcholine transporter activity | | Transporter |
| Kinesin-like protein KIF21A (Kif21a) | ATP hydrolysis activity<br>microtubule motor activity<br>microtubule binding | microtubule-based movement | microtubule binding motor protein |
| CLIP-associating protein 2 (Clasp2) | microtubule binding | establishment of mitotic spindle localization<br>mitotic spindle assembly | non-motor microtubule binding protein |
| IQ motif and SEC7 domain-containing protein 2 (Iqsec2) | - | actin cytoskeleton organization | guanyl-nucleotide exchange factor |
| Armadillo repeat protein (Arvcf) | - | - | intermediate filament binding protein |
| DNA topoisomerase 2-beta (Top2b) | - | sister chromatid segregation<br>resolution of meiotic recombination intermediates | DNA metabolism protein |
| Histone deacetylase 2 (Hdac2) | histone deacetylase activity | heterochromatin formation | - |
| ADAMTS-like protein 2 (Adamtsl2) | metalloendopeptidase activity | proteolysis<br>extracellular matrix organization | metalloprotease |
| NF-kappa-B essential modulator (Ikbkg) | K63-linked polyubiquitin modification-dependent protein binding | positive regulation of canonical NF-kappaB signal transduction | - |
| Junction plakoglobin (Jup) | nuclear receptor binding<br>protein phosphatase binding<br>transcription coactivator activity<br>cell adhesion molecule binding | canonical Wnt signaling pathway<br>cell-cell adhesion<br>positive regulation of transcription by RNA polymerase II | - |
| Plexin-A1 (Plxna1) | transmembrane signaling receptor activity | regulation of cell migration | transmembrane signal receptor |
| Guanine nucleotide-binding protein G(I)/G(S)/G(T) subunit beta-2 (Gnb2) | signaling receptor complex adaptor activity | G protein-coupled receptor signaling pathway | heterotrimeric G-protein |

| | | | |
|---|---|---|---|
| Metabotropic glutamate receptor 5 (Grm5) | neurotransmitter receptor activity<br>adenylate cyclase inhibiting G protein-coupled glutamate receptor activity | G protein-coupled receptor signaling pathway<br>trans-synaptic signaling<br>regulation of synaptic transmission, glutamatergic<br>glutamate receptor signaling pathway | G-protein coupled receptor |
| DCC-interacting protein 13-alpha (Appl1) | - | signaling | scaffold/adaptor protein |
| Insulin receptor substrate 1 (Irs1) | phosphatidylinositol 3-kinase binding<br>signaling receptor binding | insulin receptor signaling pathway | scaffold/adaptor protein |
| Nucleoside diphosphate kinase B (Nme2) | nucleoside diphosphate kinase activity | regulation of apoptotic process | Kinase |
| Leucine zipper protein 2 (Luzp2) | - | - | leucine zipper protein |
| SOGA-3 (Soga3) | - | regulation of autophagy | - |

**Table S2. List of primers and synthetic DNA oligomers (gBlocks) used.**

| OLIGONUCLEOTIDES | SEQUENCE (5'-3') | |
|---|---|---|
| Primers | | |
| Genotyping: WT allele | AAGGGACCCTCCGTCCTCTC | *GGGCCCGGTGCAATACACGC* |
| Genotyping: mutant allele | CATCGGGCATGGATCCTCAGTTC | CATCGGGCATGGATCCTCAGTTC |
| pFUGW_TARPγ2-FLAG insert 1 | CTCTAGAGGATCCCCGGGTAGCC<br>ACCATGGGGCTGTTTGATCGAG | GGTCCACAACTGTATCCAGGACT<br>ACAAAGACGATGACGACA |
| pFUGW_TARPγ2-FLAG insert 2 | CTACAAAGACGATGACGACAAGA<br>AGGACAGCAAGGACTCTCTC | CGGACCACGCCCGTATGACGATA<br>TCAAGCTTATCGATA |
| pFUGW_TARPγ3-FLAG insert 1 | CTCTAGAGGATCCCCGGGTAGCC<br>ATGGATGAGGATGTGTGACAGAG | *GTTCCACAATTCCACACCCGACT*<br>*ACAAAGACGATGACGACA* |
| pFUGW_TARPγ3-FLAG insert 2 | CTACAAAGACGATGACGACAAGA<br>AAGAGTTCAAAGAGTCACT | CGCACCACGCCCGTCTGACGATA<br>TCAAGCTTATCGATA |
| pFUGW_TARPγ4-FLAG insert 1 | CTCTAGAGGATCCCCGGGTAGCC<br>ATGGATGGTGCGATGCGACCGC | AGGTGCATGACTTTTTCCAGGAC<br>TACAAAGACGATGACGACA |
| pFUGW_TARPγ4-FLAG insert 2 | CTACAAAGACGATGACGACAAGC<br>AGGACCTGAAGGAAGGTTT | CGGACGACCCCTGTGTGACGATA<br>TCAAGCTTATCGATA |
| gBlocks | | |
| pFUGW_TARPγ8-FLAG | CTCTAGAGGATCCCCGGGTAGCCATGGATGACCATCGCCATTAGCACT<br>GATTACTGGTTGTATACAAGAGCCCTCATCTGCAATACTACCAATCTCAC<br>TGCCGGTGGAGACGACGGACCCCACACCGTGGTGGCGGCGGTGCAT<br>CAGAGAAGAAGGATCCAGGTGGGTTGACCCACTCTGGCTTGTGGAGGA<br>TCTGCTGTTTGGAAGGGTTAAAAAGAGGAGTCTGTGTGAAAATCAATCA<br>TTTCCCGGAGGATACGGACTACGATCACGACAGCGCTGAGTATCTATTA<br>CGAGTTGTCCGGGCTTCCAGCATCTTTCCTATCCTTAGCGCCATTCTGC<br>TGCTGCTTGGGGGTGTGTGTGTGGCGGCCTCCCGAGTATACAAATCAA<br>AGAGGAACATCATTCTCGGTGCAGGGATACTGTTTGTGGCAGCAGGCC<br>TGAGCAACATAATTGGCGTGATCGTTTACATCTCAGCCAACGCAGGAGA<br>ACCTGGACCGAAGCGGGATGAGGAAAAGAAAAATCATTACTCTTATGGT<br>TGGTCATTCTATTTTGGCGGGCTATCTTTCATTCTGGCCGAGGTTATAG<br>GCGTGCTTGCAGTCAACATCTATATAGAAAGAAGCAGGGAGGCGCATT<br>GCCAGTCTAGAAGCGATCTGCTCAAAGCCGGCGGTGGAGCAGGCGGC<br>AGTGGAGGGTCAGGCCCCTCGGCCATTCTCAGGCTGCCAAGTTACCGC<br>TTCCGCTACCGCCGGCGATCCAGATCTAGTTCCAGGTCAAGCGAGCCT<br>AGTCCTTCGCGGGACGCCTCTCCAGGAGGACCCGGGGGCCCTGGGTT<br>TGCCTCCACGGACATTTCCATGTATACACTTAGCCGTGACCCCAGTAAG<br>GGCTCCGTGGCCGCAGGTCTGGCGGGGGCTGGAGGCGGCGGAGGAG<br>GAGCCGTGGGTGCTTTCGGCGGCGCTGCTGGGGGCGCAGGGGGCGG<br>AGGCGGAGGCGGCGGCGGGGCGGGTGCCGAGCGGGACCGCGGAGG<br>GGCTTCCGGATTTCTCACACTGCACAACGCCTTCCCCAAGGAAGCTGG<br>CGGGGGAGTCACAGTCACTGTAACCGGGCCTCCAGCCCCGCCCGCTC<br>CTGCACCACCAGCTCCCTCTGCTCCCGCCCCCGGGACCCTGGACTACA<br>AAGACGATGACGACAAGGCCAAGGAGGCCGCCGCCTCCAACACCAACA<br>CGCTCAACAGGAAAACCACGCCTGTGTAGCGATATCAAGCTTATCGATA | |

**Table S3. Statistical analysis of the data.** Details concerning the number of independent experiments, statistical tests used and *p*-values.

| FIGURE | PANEL | STATISTICAL TEST | *p*-value | POST-HOC TEST | *p*-value | n | N | F-test or Brown-Forsythe |
|---|---|---|---|---|---|---|---|---|
| Figure 1 | | | | | | | | |
| Manders coefficient for the co-localization of Kv7.2 and TARP γ-2 (dSTORM) | D | *n/a* | n/a | n/a | n/a | 7 | 4 | n/a |
| splitFAST fluorescence over time | G | *n/a* | n/a | n/a | n/a | 55 | 2 | n/a |
| Figure 2 | | | | | | | | |
| Effect of type I TARPs on Kv7.2 current density | B | Kruskal-Wallis | <0.0001 | Dunn's multiple comparison test (*p*-values corrected for multiple comparisons) | Kv7.2 vs Kv7.2+γ2<br>*p* = 0.0021<br>Kv7.2 vs Kv7.2+γ3<br>*p* = 0.0001<br>Kv7.2 vs Kv7.2+γ4<br>*p* < 0.0001<br>Kv7.2 vs Kv7.2+γ8<br>*p* > 0.9999 | ≥32 | ≥3 | n/a |
| Effect of type I TARPs on Kv7.2 voltage-dependent activation | C | n/a | n/a | n/a | n/a | ≥32 | ≥3 | n/a |
| Effect of type I TARPs on Kv7.2 half-activation potential ($V_{50}$) | D | One-way ANOVA | <0.0001 | Dunn's multiple comparison test (*p*-values corrected for multiple comparisons) | Kv7.2 vs Kv7.2+γ2<br>*p* = 0.0307<br>Kv7.2 vs Kv7.2+γ3<br>*p* < 0.0001<br>Kv7.2 vs Kv7.2+γ4<br>*p* < 0.0001<br>Kv7.2 vs Kv7.2+γ8<br>*p* < 0.0001 | ≥32 | ≥3 | *p* = 0.1990 |

| | | | | | | | | |
|---|---|---|---|---|---|---|---|---|
| Effect of type I TARPs on Kv7.2 surface expression | F | Kruskal-Wallis | <0.0001 | Dunn's multiple comparison test (*p*-values corrected for multiple comparisons) | Kv7.2 vs Kv7.2+γ2<br>*p* < 0.0001<br>Kv7.2 vs Kv7.2+γ3<br>*p* < 0.0001<br>Kv7.2 vs Kv7.2+γ4<br>*p* < 0.0001<br>Kv7.2 vs Kv7.2+γ8<br>*p* = 0.0457 | ≥57 | 4 | n/a |
| Effect of type I TARPs on Kv7.2 total expression | G | Kruskal-Wallis | 0.0296 | Dunn's multiple comparison test (*p*-values corrected for multiple comparisons) | Kv7.2 vs Kv7.2+γ2<br>*p* = 0.0457<br>Kv7.2 vs Kv7.2+γ3<br>*p* = 0.0162<br>Kv7.2 vs Kv7.2+γ4<br>*p* = 0.4397<br>Kv7.2 vs Kv7.2+γ8<br>*p* > 0.9999 | ≥57 | 4 | n/a |
| Figure 3 | | | | | | | | |
| Kv7.2 cluster density | E | *Two-tailed Welch's t-test* | 0.0459 | n/a | n/a | ≥6 | ≥3 | *p* = 0.0235 |
| Kv7.2 cluster area | F | *Two-tailed Mann-Whitney test* | <0.0001 | n/a | n/a | 1800 | ≥3 | n/a |
| Kv7.2 cluster local density | G | *Two-tailed Mann-Whitney test* | <0.0001 | n/a | n/a | 1800 | ≥3 | n/a |
| Effect of TARP γ-2 overexpression on Kv7.2 cluster area | I | *Two-tailed Mann-Whitney test* | <0.0001 | n/a | n/a | 787 | 4 | n/a |
| Effect of TARP γ-2 knockdown on Kv7.2 cluster area | K | Two-tailed Mann-Whitney test | 0.0031 | n/a | n/a | 1050 | 5 | n/a |
| Figure 4 | | | | | | | | |

| | | | | | | | | |
|---|---|---|---|---|---|---|---|---|
| Effect of TARP γ-2 knockdown on retigabine-induced hyperpolarizing shift of the membrane potential | B | *Two-tailed Unpaired t-test* | 0.0063 | n/a | n/a | ≥25 | 8 | *p* = 0.5904 |
| Effect of TARP γ-2 knockdown on neuronal membrane resting potential | C | *Two-tailed Unpaired t-test* | 0.0936 | n/a | n/a | ≥25 | 8 | *p* = 4411 |
| Effect of TARP γ-2 knockdown on Kv7.2 current density | E | Two-tailed Mann-Whitney test | 0.0389 | n/a | n/a | ≥22 | 10 | n/a |
| Figure 5 | | | | | | | | |
| Effect of TARP γ-2 V143L on Kv7.2 current density | C | *Kruskal-Wallis* | 0.0060 | Dunn's multiple comparison test (*p*-values corrected for multiple comparisons) | Kv7.2 vs Kv7.2+γ2 WT *p* = 0.0066<br>Kv7.2 vs Kv7.2+γ2 V143L *p* = 0.4811 | ≥15 | ≥3 | n/a |
| Retigabine-induced reduction in neuronal firing in Het-γ2 V143L mice | E | *Two-way Repeated Measures ANOVA* | 0.4055 (interaction)<br>0.0387 (genotype)<br><0.0001 (injected current) | n/a | n/a | 7 | ≥4 | n/a |

| | | | | | | | | |
|---|---|---|---|---|---|---|---|---|
| mAHP amplitude in subicular pyramidal neurons of Het-γ2 V143L mice | G | *Two-way Repeated Measures ANOVA* | 0.3398 (interaction)<br>0.0011 (genotype)<br>0.0020 (retigabine) | Šídák's multiple comparison test (*p*-values corrected for multiple comparisons) | Cont. WT vs Ret. WT<br>*p* = 0.0090<br>Cont. Het-γ2 V143L vs Ret. Het-γ2 V143L<br>*p* = 0.1156<br>Cont. WT vs Cont. Het-γ2 V143L<br>*p* = 0.0060<br>Ret. WT vs Ret. Het-γ2 V143L<br>*p* = 0.0006 | 7 | ≥4 | n/a |
| PTZ-induced seizures in Het-γ2 V143L mice | H | *Two-tailed Unpaired t-test* | 0.0097 | n/a | n/a | n/a | 5 | *p* = 0.3278 |
| XE991-induced seizures in Het-γ2 V143L mice | H | Two-tailed Unpaired *t*-test | 0.0166 | n/a | n/a | n/a | ≥6 | *p* = 0.1086 |
| Figure S2 | | | | | | | | |
| Effect of KCl-induced depolarization on γ2-Kv7.2 PLA puncta density | B | *Two-tailed Mann-Whitney test* | 0.0072 | n/a | n/a | 29 | 3 | n/a |
| Effect of KCl-induced depolarization on AIS plasticity | D | *Two-tailed Mann-Whitney test* | 0.0016 | n/a | n/a | ≥99 | 11 | n/a |
| Effect of KCl-induced depolarization on GluA synaptic clusters intensity | E | *Two-tailed Mann-Whitney test* | 0.7940 | n/a | n/a | 23 | 2 | n/a |
| Effect of KCl-induced depolarization on GluA total surface clusters intensity | G | *Two-tailed Mann-Whitney test* | 0.8109 | n/a | n/a | 23 | 2 | n/a |

| | | | | | | | | |
|---|---|---|---|---|---|---|---|---|
| Effect of KCl-induced depolarization on neuronal cell viability | I | *Two-tailed Unpaired t-test* | 0.8126 | n/a | n/a | n/a | 5 | *p* = 0.7024 |
| Figure S3 | | | | | | | | |
| Effect of TARP γ-2 on Kv7.2/Kv7.3 current density | B | *Two-tailed Unpaired t-test* | 0.3643 | n/a | n/a | ≥16 | ≥3 | *p* = 0.9509 |
| Effect of TARP γ-2 on Kv7.2/Kv7.3 voltage-dependent activation | C | *n/a* | n/a | n/a | n/a | ≥16 | ≥3 | n/a |
| Effect TARP γ-2 on Kv7.2/Kv7.3 half-activation potential ($V_{50}$) | D | *Two-tailed Unpaired t-test* | 0.4240 | n/a | n/a | ≥16 | ≥3 | *p* = 0.5312 |
| Effect of TARP γ-2 overexpression on TARP γ-2 total expression | F | *Two-tailed Mann-Whitney test* | <0.0001 | n/a | n/a | ≥88 | 4 | n/a |
| Effect of TARP γ-2 overexpression on Kv7.2 total expression | G | Two-tailed Mann-Whitney test | <0.0001 | n/a | n/a | ≥88 | 4 | n/a |
| Figure S4 | | | | | | | | |
| Distribution of Kv7.2 cluster diameter | A | *n/a* | n/a | n/a | n/a | 6 | 4 | n/a |
| Effect of TARP γ-2 overexpression on Kv7.2 cluster density | B | *Two-tailed Mann-Whitney test* | 0.1320 | n/a | n/a | 6 | 4 | *p* = 0.0891 |
| Effect of TARP γ-2 overexpression on Kv7.2 cluster local density | C | *Two-tailed Mann-Whitney test* | <0.0001 | n/a | n/a | 787 | 4 | n/a |
| Effect of TARP γ-2 knockdown on Kv7.2 cluster density | D | *Two-tailed Mann-Whitney test* | 0.7639 | n/a | n/a | ≥8 | 5 | n/a |

| | | | | | | | | |
|---|---|---|---|---|---|---|---|---|
| Effect of TARP γ-2 knockdown on Kv7.2 cluster local density | E | Two-tailed Mann-Whitney test | 0.6899 | n/a | n/a | 1050 | 5 | n/a |
| Figure S5 | | | | | | | | |
| Input resistance of subicular pyramidal neurons of Het-γ2 V143L mice | D | *Two-tailed Unpaired t-test* | 0.0033 | n/a | n/a | ≥11 | ≥4 | *p* = 8603 |
| Membrane time constant of subicular pyramidal neurons of Het-γ2 V143L mice | E | Two-tailed Unpaired *t*-test | 0.0011 | n/a | n/a | ≥11 | ≥4 | *p* = 2468 |
| Membrane capacitance of subicular pyramidal neurons of Het-γ2 V143L mice | F | Two-tailed Unpaired *t*-test | 0.7784 | n/a | n/a | ≥11 | ≥4 | *p* = 3523 |
| Rheobase of subicular pyramidal neurons of Het-γ2 V143L mice | G | Two-tailed Unpaired *t*-test | 0.0405 | n/a | n/a | ≥11 | ≥4 | *p* = 8021 |
| Action potential threshold of subicular pyramidal neurons of Het-γ2 V143L mice | H | Two-tailed Unpaired *t*-test | 0.0129 | n/a | n/a | ≥11 | ≥4 | *p* = 7110 |
| Resting membrane potential of subicular pyramidal neurons of Het-γ2 V143L mice | I | Two-tailed Unpaired *t*-test | 0.2618 | n/a | n/a | ≥11 | ≥4 | *p* = 3867 |
| Effect of retigabine on the firing frequency of subicular pyramidal neurons of WT mice | J | n/a | n/a | n/a | n/a | 7 | ≥4 | n/a |

| | | | | | | | | |
|---|---|---|---|---|---|---|---|---|
| Effect of retigabine on the firing frequency of subicular pyramidal neurons of Het-γ2 V143L mice | K | n/a | n/a | n/a | n/a | 7 | ≥4 | n/a |

n corresponds to the number of cells or images analyzed
N corresponds to the number of independent experiments or the number of animal
